# Phylogenetic reconciliation supports a methanogenic ancestor of the Archaea and a derived origin for host-associated lineages

**DOI:** 10.1101/2025.11.11.687807

**Authors:** Wen-Cong Huang, Nina Dombrowski, Tara A. Mahendrarajah, Noah Wahl, Alexandros Stamatakis, Gergely J. Szöllősi, Tom A. Williams, Anja Spang

## Abstract

The phylogeny of the Archaea continues to be revisited and revised as new groups are discovered and phylogenetic methods improve, but key questions about their early evolution remain. It has been suggested that the root of the Archaea may lie on, or potentially within, any of three major groups - the Euryarchaeota, TACK+Asgard clade, and DPANN, the last of which includes many host-associated and genome-reduced lineages. These root hypotheses make starkly different predictions about the nature of early archaeal evolution: for example, a root on or within DPANN might suggest a small-genome and host-associated ancestor, with methanogenesis, the hallmark metabolism of the Archaea, evolving later. Here, we investigate the position of the archaeal root and the nature of the last archaeal common ancestor using a range of phylogenetic approaches, including the best available site- and branch-heterogeneous substitution models, and new gene tree-species tree reconciliation models that capture changes in rates of gene duplication, loss and transfer across the phylogeny. Our analyses converge on a narrow archaeal root region at/near the base of the Euryarchaeota, supporting hypotheses in which the Last Archaeal Common Ancestor (LACA) was a complex, free-living (hyper-)thermophilic methanogen. We recover DPANN as the sister group to TACK and Asgard archaea, and suggest that their genome evolution has been characterised by episodes of genome streamlining and expansion, driven by gene loss and transfer.

## Introduction

Over the past 15 years, developments in environmental genomics have led to the discovery of major new lineages that have fuelled new interest in, and debates about, the deep structure of the tree of life^1–7^. In particular, the discovery of clades of genetically diverse, small-genome sized, host-associated archaea and bacteria - i.e. the DPANN archaea (an acronym of the first described phyla: Diapherotrites, Parvarchaeota, Aenigmarchaeota, Nanoarchaeota and Nanohaloarchaeota, now referred to as Nanobdellati^8^) and the CPR bacteria (Candidate Phyla Radiation bacteria, now termed Patescibacteria^9^/Minisyncoccota^10^) - have catalysed new controversies about the antiquity of symbiosis and parasitism in life’s history. Initial phylogenetic analyses placed DPANN and CPR at the base of their respective domains, potentially implying archaeal and bacterial ancestors of only moderate size and cellular and metabolic complexity. Subsequent analyses using improved phylogenetic methods resolved CPR as a derived clade related to Chloroflexota^11–15^ and suggested that the last bacterial common ancestor was likely a complex and free-living cell akin in complexity to modern bacteria^12^. However, resolving or approaching a similar degree of consensus on the position of DPANN within Archaea has proven more challenging, with published analyses recovering phylogenies that have starkly different consequences for our understanding of archaeal origins and the sequence of metabolic evolution within this domain^16–19^.

The primary question is whether DPANN branch at the base of the archaeal domain, or alternatively whether they occupy a derived position, as sister to or branching within one of the other major lineages, which include Euryarchaeota (also referred to as Methanobacteriati^8^), Thermoproteota (TACK Archaea^20^ / Thermoproteati^8^) or Asgard archaea (Asgardarchaeota^21^/ Promethearchaeota^22^)^4,14,17–19,23,24^. Resolving the placement of DPANN in the archaeal tree is critically important because it informs the nature of LACA and the tempo and mode of archaeal genome evolution. This is because DPANN predominantly, though not exclusively^24–28^, comprise organisms with reduced genomes and apparently limited metabolic capabilities. Indeed, the first cultivated representatives^29–36^ of the DPANN were shown to be obligate archaeal ectosymbionts depending on distinct archaeal^24–26,37^ and potentially even bacterial hosts^38^. If DPANN branch at the base of the Archaea, this could suggest that LACA was a relatively simple and small-genome sized organism, with methanogenesis - the hallmark metabolism of the Archaea^39–41^ - evolving later, after the divergence of DPANN from Euryarchaeota, TACK, and Asgard archaea. Alternatively, if the archaeal root lies within one of those alternative lineages with DPANN being derived, LACA may instead have been a *bona fide* complex, free-living organism.

Specifically, determining the position of DPANN within the archaeal phylogeny has proven challenging. Some recent studies have resolved DPANN as the earliest-diverging branch in the archaeal tree^13,14,18^, or with, or within, Euryarchaeota^17^, while others have suggested that DPANN might actually be polyphyletic, evolving multiple times from within the Euryarchaeota via parallel events of genome reduction^42–44^. Each of these inferred positions has been suggested to be induced by various phylogenetic artifacts. For example, the distinctive evolutionary mode of DPANN - which, as symbionts may both evolve fast and lose many genes otherwise conserved in archaea - might artifactually draw them to the root of the tree due to long branch attraction (LBA)^17,19,23,45^. On the other hand, alternative DPANN placements at derived positions within the archaeal tree have also been suggested to result from phylogenetic artifacts. For example, the placement of DPANN with Euryarchaeota in some analyses has been suggested to result from unmodelled horizontal gene transfer (HGT) between DPANN symbionts and their euryarchaeotal hosts^46–48^, or to result from systematic phylogenetic error^49^. A recent analysis using a newly-developed non-stationary phylogenetic model that captures variation in sequence composition across both the sites of the alignment and the branches of the tree - two pervasive but often un-modelled features of sequence evolution - provided support for a derived position of DPANN branching sister to Euryarchaeota, with the root of the archaeal tree lying between the TACK+Asgard archaea and all other archaea^17^.

The studies summarised above used a range of distinct approaches to root the archaeal phylogeny. The most widely-used rooting approach is to use an outgroup - that is, to include a bacterial outgroup to root the archaeal tree on the inter-domain branch. This approach is conceptually straightforward but may be problematic for rooting entire domains, because the long inter-domain branch might induce errors in the within-Archaea topology^50,51^. In particular, the basal position of DPANN in published trees might result from long branch attraction between the DPANN and inter-domain stems. As an alternative to outgroup rooting, non-reversible or non-stationary models can be used to root the tree based on capturing directional aspects of the evolutionary process, such as long-term trends in amino acid substitution patterns or changing amino acid compositions over time^52–56^. However, the power of these approaches to root trees depends on the strength of the time-dependent signal in the data and the ability of the model to adequately describe the underlying processes^54^. These signals are sometimes weak for anciently diverged sequences^52^. A third approach is to use phylogenetic reconciliation, which can assess distinctly rooted species trees based on inferred evolutionary events that have a clear direction in time, such as gene duplications, losses and transfers^12,18,57,58^. However, as with substitution models, the accuracy of the results depends on the adequacy with which the model captures the underlying processes^12^. In addition to these modelling concerns, different studies have used distinct sets of phylogenetic “marker” genes - genes that are thought to provide a reliable trace of vertical archaeal evolution - and protein families in the case of reconciliations with the differing properties of these datasets introducing additional variation into the results^13,17–19,23,45,49^. While none of the available rooting methods or datasets are consummate, the hope is that, as methods improve, they may converge on a consistent root position supported - or at least not strongly rejected - by distinct analytical approaches.

Here, we investigate the root of the archaeal tree, using the best-available outgroup-rooting methods and recently developed non-reversible and non-stationary substitution models^52,59,60^. For the first time, we also apply a new class of more flexible and better-fitting phylogenetic reconciliation methods that we recently developed^57^, and which allow the model of genome evolution to vary across the rooted species tree. The reconciliation method not only allows us to assess the support for various root hypotheses but also to reconstruct historical patterns of genome evolution across Archaea.

## Results

### Outgroup rooting has only limited power to resolve the root of Archaea

We re-visited suitable markers and outgroup rooting on an archaeal tree using a focal dataset of 257 Archaea and 103 Bacteria (Table S1) based on 25 vertically-evolving markers that did not violate the requirement of domain monophyly (Table S2)^13^. Specifically, we used several rounds of manual curation to assemble a dataset of 257 phylogenetically representative, high-quality (low contamination, high completeness) genomes sampled from across the known diversity of Archaea (Table S1) (see methods). The best-known maximum likelihood tree under the best-fitting site-heterogeneous model (LG+C60+F+R; chosen by Bayesian Information Criterion (BIC)) placed the archaeal root between DPANN archaea and all other archaea (Fig. S1, DPANN root, 96.2 and 88 ultrafast bootstrap support^61^ respectively for the monophyly of DPANN lineage and the monophyly of Euryarchaeota+TACK+Asgard), consistent with some previous analyses^4,13,18,24,62^. However, previous work has shown that there is sometimes limited information about the archaeal root position in such “tree of life”-scale analyses^12–14,18,63^, so that the best-known maximum likelihood tree is not necessarily significantly better than trees with alternative root positions. To test this possibility on our data, we generated a range of rooting hypotheses (Table S3), including key hypotheses from the literature as well as other plausible root positions (e.g., between each of the major archaeal taxonomic clades), by relocating the bacterial outgroup’s attachment point to the archaeal ingroup. For each candidate root, we inferred the best-known ML tree while remaining consistent with that root hypothesis using a constrained tree search in IQ-TREE 2^64^, and then assessed whether the likelihoods of these trees were significantly worse than the unconstrained ML tree using an approximately unbiased (AU) test^65^. This analysis rejected some hypotheses, such as a root between the DPANN Cluster 2 archaea (a DPANN clade that includes Nanohaloarchaeota, Aenigmatarchaeota, Nanoarchaeota, Woesearchaeales and Pacearchaeales)^62^ and all other Archaea (P < 0.05; Table S3). However, various other hypotheses, some of which received literature support e.g. DPANN root^18^, a root between TACK+Asgard archaea and all other archaea (hereafter, TACK+Asgard root)^17,45^, a root between a clade comprising Halobacteriota and Thermoplasmatota and all other archaea (hereafter, Halobacteriota+Thermoplasmatota root)^19,23^, and a root between Euryarchaeota and all other archaea (hereafter, Euryarchaeota root), remained statistically indistinguishable (Table S3). These results are consistent with another recent analysis of the tree of life using site-heterogeneous models^63^, in which the maximum likelihood analysis supported TACK+Asgard on one side of the archaeal root, and Euryarchaeota+DPANN on the other (“Topology 1” in^63^); however, this root was statistically indistinguishable from a root between DPANN and all other Archaea based on the AU test.

### Inferring an unrooted species tree of Archaea

Given the lack of resolution in the outgroup rooting analyses, we next built an unrooted phylogenetic tree for subsequent use of outgroup-free rooting using non-reversible, non-stationary, and phylogenetic reconciliation models. Since tree search is computationally prohibitive under these models, we followed a two-step procedure in which we initially estimated an unrooted archaeal species tree using a concatenation of phylogenetic marker genes, then searched for the root position on this fixed tree using a variety of approaches. We extracted the sequences for a set of 51 marker genes previously suggested to evolve vertically within the Archaea^62^ from our archaeal focal dataset of 257 representative genomes; manual inspection (see methods) suggested that 45 of those markers were suitable for concatenation (i.e. they had no indication for inter-domain HGT and phylogenetic incongruence) (Table S4). We then inferred an unrooted tree using the best-fitting model (LG+C60+F+R; chosen by BIC; see Methods), which accounts for different rates of change among amino acids and variation in evolutionary rate and sequence composition across the sites of the sequence alignment. This phylogeny recovered the clanhood (that is, monophyly conditional on a root outside the group^66^) of three archaeal groups with maximal or near-maximal support based on the ultrafast bootstrap (UFBOOT)^61^ and Shimodaira–Hasegawa approximate likelihood ratio test (SH-aLRT)^67^ metrics: Euryarchaeota (100/99), TACK+Asgard archaea (100/100) and DPANN archaea (100/100) (Fig. 1). To evaluate whether the inferred topology might be affected by across-branch variation in amino acid composition - a feature of sequence evolution not accounted for by the LG+C60+F+R model - we incrementally filtered out 10%-80% of the most compositionally-biased sites from the alignment and re-inferred the species trees, which produced highly congruent species trees that all recovered the three archaeal groups (Fig. S2). This suggests that key features of our topology are not artifacts of biased sequence composition. The inferred topology was in good agreement with previous analyses - for example, recovering Euryarchaeota, TACK+Asgard and DPANN clans, as well as many of the major relationships within each of those groups^14,17,18,62,63^.

**Fig. 1.**
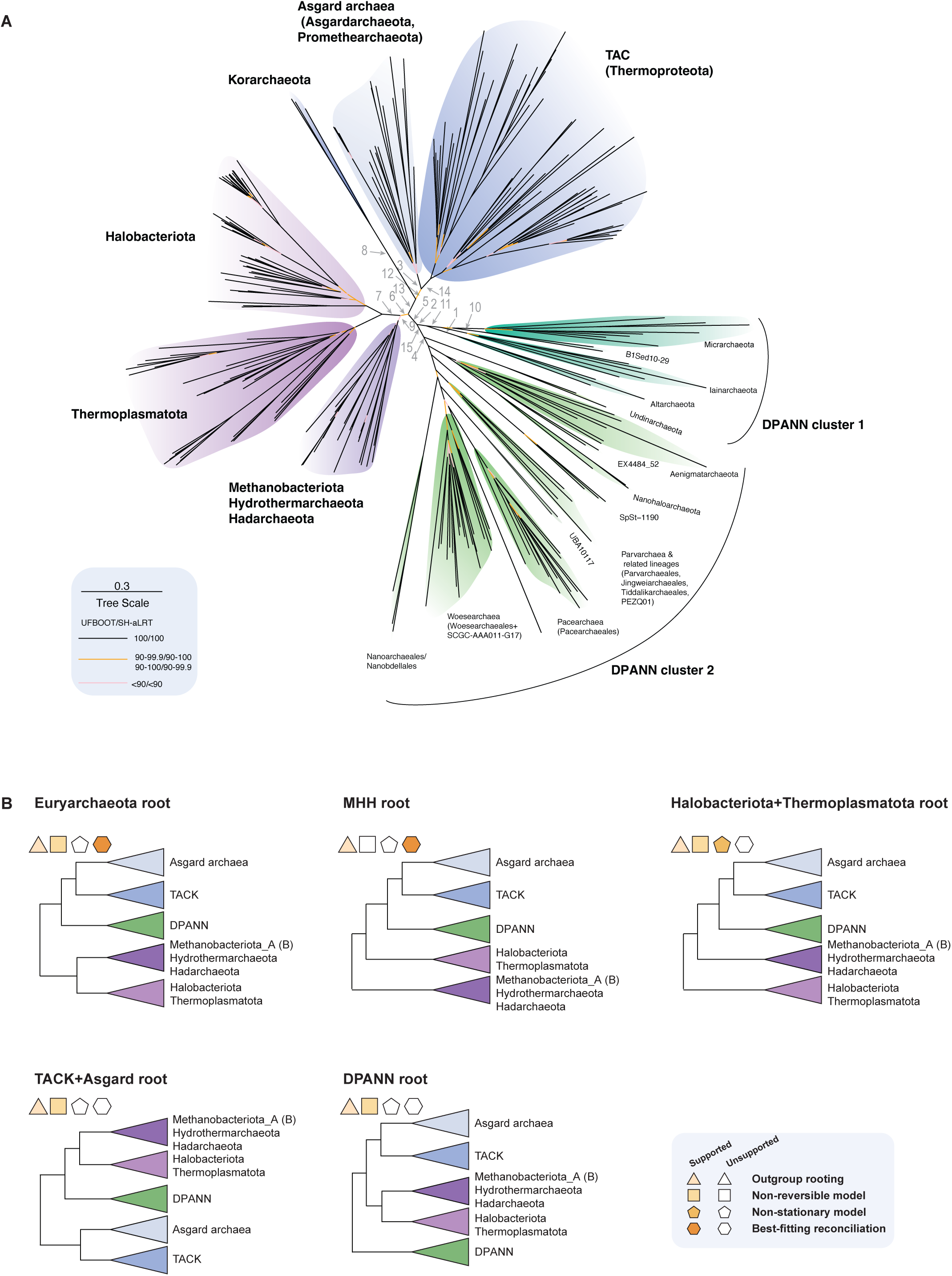
An unrooted phylogeny of the archaeal tree of life and depictions of alternative root hypotheses. (**A**) An unrooted best-known maximum likelihood phylogeny of Archaea, inferred from a supermatrix of 45 vertically-evolving marker genes under the best-fitting site-heterogeneous model, LG+C60+F+R. The numbers indicate the various root positions tested. (**B**) Schematic represen-tation of the most supported root positions using different root inference methods, as indicated by the color-coded shapes in the figure inlet: outgroup rooting, non-reversible, non-stationary (GFmix) and the best-fitting reconciliation models (See Methods) (i.e., that cannot be rejected by an approximately unbiased (AU) test; P>0.05). AU test results of outgroup rooting, non-reversible model, non-stationary (GFmix) model and best-fitting reconciliation model are provided in Tables S3, S6, S7, and S12. Among these, the best-fitting reconciliation model supports a root between Euryarchaeota and all other Archaea and a root between a clade compris-ing Methanobacteriota_A(B), Hydrothermarchaeota and Hadarchaeota on one side, and all other Archaea on the other. Scale Bar: average substitutions per site.

One lineage that has proven difficult to place in previous analyses has been the Altiarchaeota, which branch with DPANN but are less genome-reduced and appear capable of free-living growth^27^. We recovered Altiarchaeota at the base of DPANN Cluster 1 (Fig. 1), in agreement with various previous studies^62,68–70^. By contrast, other recent studies placed the Altiarchaeota as the earliest-diverging branch within DPANN as a whole^17,19^. These studies made use of an alternative set of marker genes with a range of functions beyond information processing, including metabolic genes such as subunits of ATP synthase. While an expanded marker set is valuable, one challenge is the higher rate of horizontal gene transfer in non-ribosomal proteins^13^. We re-analyzed these genes and, after filtering out marker genes with instances of HGT within Archaea and between Archaea and Bacteria, recovered Altiarchaeota at the base of Cluster 1 (see Supplementary Material for further details, Fig. S3, Table S5).

### Analysis with non-reversible and non-stationary substitution models supports a root within Euryarchaeota

We next used non-reversible and non-stationary substitution models to infer the root position on the fixed unrooted species tree. We first performed analyses under the non-reversible “NONREV+G” model^52^ implemented in IQ-TREE 2^64^. In this model, exchangeabilities between amino acid states are non-reversible; for example, the probability of change from D (aspartic acid) to E (glutamic acid) is different to the probability from E to D; as a result, trees inferred under NONREV+G are intrinsically rooted, with the root information derived from long-term trends in protein evolution (a long-term difference in the rate at which exchanges occur between pairs of amino acids; e.g., the relative rate of change of amino acid from D to E was 0.43, while the rate from E to D was 0.20, in our dataset). When applying such considerations to the 257 archaeal species, the 45-marker gene dataset inferred the best known maximum likelihood root between Korarchaeota and all other archaea (Korarchaeota root, rootstrap support: 61.5)^52^ (Fig. S4). As in the outgroup-rooting analyses, we used an AU test^65^ on site-wise likelihoods to evaluate whether the Korarchaeota root provided a significantly better fit to the data than alternative positions that are rerooted at every branch of the inferred maximum likelihood tree. As with outgroup rooting, the analysis indicated that a variety of root hypotheses could not be rejected under the NONREV+G model for these data, including roots on Korarchaeota, Thermoproteota, DPANN, Euryarchaeota, and TACK+Asgard archaea (Table S6, Fig. S4).

Non-stationary models can also be used to infer the root of a tree accounting for changes in the substitution process over time. One such model is the GFmix model^17,59,60^, which models changing nucleotide or amino acid sequence compositions over the branches of the phylogenetic tree. This kind of compositional change is manifest in real data. For example, halophilic archaea were reported to be enriched in D and E while depleted in I (isoleucine) and K (lysine) compared to non-halophilic archaea^49,71^. GFmix is particularly appealing for analyses of deep microbial evolution because it combines the modelling of across-site compositional heterogeneity (as in the site-heterogeneous C60 model described above) with across-branch compositional variation - both pervasive features of these datasets - in a single analysis. When applied to our 45-gene supermatrix, a root within the Euryarchaeota, i.e., the Halobacteriota+Thermoplasmatota root^19,23^ had the highest likelihood with all other tested roots being rejected by an AU test (P< 0.05, Table S7). By contrast, a recent analysis of deep archaeal phylogeny using GFmix and a different marker gene set, tested a range of root hypotheses and reported that a root between TACK+Asgard and all other Archaea had the highest likelihood among the trees tested, with DPANN being derived from within a paraphyletic Euryarchaeota^17^. We repeated this analysis on the original dataset, but including a broader set of root hypotheses (9 more positions in addition to the DPANN root and TACK+Asgard root) (Table S8) and found that several of these had significantly higher likelihoods. Among those we tested, the Halobacteriota+Thermoplasmatota root^19,23^ had the highest likelihood in agreement with our new results from the 45-gene concatenate (being 8.6 likelihood units higher than the TACK+Asgard root; see Table S7). All other tested roots, including a root between TACK+Asgard and all other Archaea^17^, were rejected by the AU test (P < 0.05; see Table S8 and Supplementary Material).

Overall, our analyses of these datasets using outgroup rooting, non-reversible and non-stationary models suggest that the root of the Archaea remains a challenging problem, yielding rather similar likelihoods for a variety of plausible root hypotheses, most of which are within 3 nodes of one another on the unrooted species tree. To complement these analyses, we explored rooting using gene tree-species tree reconciliation, which makes use of a substantially larger fraction of the available genome data and might therefore have greater resolving power to assess contrasting root hypotheses^12,18,72^.

### Improved modelling of archaeal gene family evolution using branch-heterogeneous phylogenetic reconciliation models

Early reconciliation analyses of the archaeal root made use of a limited set of archaeal diversity (e.g. 60 genomes in Williams et al.,^18^), and a model that made the simplifying assumption that rates of duplication (D), transfer (T) and loss (L) do not vary across the branches of the species tree. However, various lines of evidence suggest that D, T and L rates do vary among archaeal clades. For example, duplications appear to be more common in Asgard archaea^18,73^, while losses are more common among DPANN lineages^18^. In principle, a failure to accommodate for these varying rates might mislead phylogenetic analyses that rely on reconciliation methods, just as failure to accommodate varying evolutionary rates or compositions among alignment sites can induce high support for the incorrect trees in traditional sequence-based phylogenetics^59,74,75^. Both of these limitations can now be overcome: the genome sampling of the archaeal domain is far richer than previously, and we and others have recently developed more flexible phylogenetic reconciliation models that relax the assumptions of constant D, T and L rates across the species tree. Our new model is implemented in AleRax^57^.

To begin, we fit a three-parameter branch-homogeneous reconciliation model (one D, T and L rate, respectively, common to all branches, referred to as global in what follows) to the dataset of 257 archaeal genomes and 7059 gene family phylogenies, and used an AU^65^ test to compare support for a range of potential root positions that were also assessed in outgroup rooting analyses (Fig. 1, 1-15). The result provided strong support for a root between DPANN Cluster 2 - a clade within DPANN - and all other archaea (referred to as DPANN Cluster 2 root, Fig. 1A) (Table S9). To gain insight into the nature of the signal for the DPANN Cluster 2 root, we investigated the properties of the gene families that favoured the DPANN Cluster 2 root over alternative root positions, using gene family filtering to explore root signals^12^. We noticed that the preference of DPANN Cluster 2 root came from gene families with a small number of species and a biased taxonomic representation (Fig. S5A-B), consistent with a “small genome attraction (SGA)” artifact, where the model prefers a root that separates smaller genomes from larger ones on the tree^18^. In principle, such a bias might arise in analyses under a branch-homogeneous DTL model if gene absences in genome-reduced lineages are over-interpreted as evidence that a given lineage never had a gene, overlooking the possibility that genes might be absent due to secondary loss. Consistent with this possibility, DPANN Cluster 2 clades predominantly comprise symbionts with reduced genomes, suggesting that they might exhibit a higher-than-average gene loss rate^26^. To test whether the assumption of a single, global loss rate might drive support for the DPANN Cluster 2 root, we added one more parameter to allow for a different loss (L) rate for DPANN archaea (DPANN_L), and reassessed root supports using an AU test^65^. Adding a DPANN-specific loss rate substantially improved reconciliation model fit (average BIC difference: 1388.46 lower, Table S10), and, as predicted, the DPANN loss rate was inferred to be higher than that for the other clades (DPANN: 0.40-0.45, other clades: 0.26-0.27). Interestingly, this analysis expanded the set of plausible roots that could not be rejected by the AU test under the global model, including two alternative root positions that are located nearby on the unrooted tree: Euryarchaeota root or TACK+Asgard root (P>0.174; Table S11), suggesting an amelioration of SGA in the analysis.

### The best-fitting reconciliation model supports a root at/near the base of Euryarchaeota

Given the shift in root support when explicitly modelling the higher loss rate in DPANN, we next explored a series of more complex reconciliation models with additional lineage-specific D, T and L parameters. Our aim was to identify a model which adequately captured lineage-specific variation in D, T and L rates across Archaea by using the BIC to assess model fit and prevent overfitting, while also evaluating how root support varies when different aspects of genome evolution are taken into account (see Supplementary Material).

These analyses indicated that the best-fitting reconciliation model for these data was a model in which each of the major clades of Archaea had their own gene family D, T and L rates (Table S10, assessed by BIC). The final model had 72 parameters, with lineage-specific DTL rates for various Euryarchaeota, TACK+Asgard, and DPANN archaeal lineages (see Methods). Under this best-fitting model (Table S10), an AU test rejected all roots except the Euryarchaeota root and a root within Euryarchaeota (between a clade comprising Methanobacteriota_A(B), Hydrothermarchaeota and Hadarchaeota, hereafter MHH root), both with a monophyletic DPANN emerging as sister to the Asgard+TACK group (Fig. 1B, P ≥ 0.48, Table S12). Support for the Euryarchaeota root and the MHH root under the branch-heterogeneous model was broad and particularly concentrated in the most widespread and taxonomically balanced families (Fig. S5I), with the Euryarchaeota/MHH root remaining the maximum likelihood root until the top 2% most widespread and balanced families had been filtered out of the analyses (Fig. S5I), in contrast to the gene family-removal analysis described above which found support for the DPANN Cluster 2 root derived from gene families with limited taxa representation present mainly on one side of the root (Fig. S5B). Both the improved fit of this model (as measured by BIC) and the congruent signal across gene families with balanced taxa representation indicate that these are the best-supported roots from our reconciliation analyses (see Supplementary Material).

Given the improved model fit and change in root support observed with the best-fitting, branch-heterogeneous reconciliation model on the 257-genome dataset, we also reanalysed the 60-genome dataset from Williams et al.,^18^, which had provided support for a DPANN root when analyzed with an earlier branch-homogeneous reconciliation model, by including a few additional rooting hypotheses. As in our focal dataset, allowing D, T and L parameters to vary across clades substantially improved the dataset’s model fit (average BIC: 463958.631, Table S13), and the best-fitting branch-heterogeneous model supported a root between Euryarchaeota and all other Archaea (P=1, Table S14, Supplementary Material). Taken together, our results on new and previous datasets demonstrate that improved modelling of variation in D, T and L rates across the tree substantially improves reconciliation model fit, and provides support for the root at/near the base of Euryarchaeota (Fig. 1B).

### Dynamics of archaeal genome evolution

Our analyses based on the best-fitting branch-wise reconciliation model suggest that archaeal gene family evolution has been mostly vertical, with a mean verticality (i.e., the proportion of genes at the bottom of a branch whose ancestry trace straight up through the same branch) estimated to be 79% (Table S15). Transferability of gene families varies significantly across functional categories under both the Euryarchaeota root and the MHH root (H=885.19, P=1.13x10^-174^ and H=878.0, P=3.75x10^-173^, degree of freedom: 20, Kruskal–Wallis test, Table S16-S17), with genes involved in defence mechanisms (such as restriction endonuclease, COG category V), and cell wall biogenesis (such as glycosyltransferase, COG category M), being the most frequently transferred, and those involved in cell cycle and informational processing the least (Fig. S6, Table S18), in line with previous findings^18,76^. Gene transfer has a strong impact on archaeal evolution: it outnumbers duplication events across nearly all (507/501 out of 513 branches for the Euryarchaeota/MHH root), and loss events on many branches (177/154 out of 513 branches). Specifically, we found that gene transfers are on average 10.87-11.18 times more frequent than duplications events in archaeal evolution (Table S19), in agreement with the finding that transfers are more frequent than duplications in prokaryotic genomes^77,78^. In line with previous studies^18,73^, Asgard archaea had the highest inferred duplication rate compared to the other archaeal clades (Fig. 2AB), whose duplication events (normalised by speciations) appeared significantly higher than its sister lineage Thermoproteota (Conoveŕs test, P<0.012, Table S20), while the transfer and loss events of Asgard archaea (normalised by speciations) did not appear to be significantly different from what we observe for other archaea with the exception of DPANN lineages (Fig. 2AB, Conoveŕs test, Table S21-22). This suggests that the eukaryote-like propensity to fix duplicate genes had already begun to develop in our Asgard ancestors prior to the origin of eukaryotes.

**Fig. 2.**
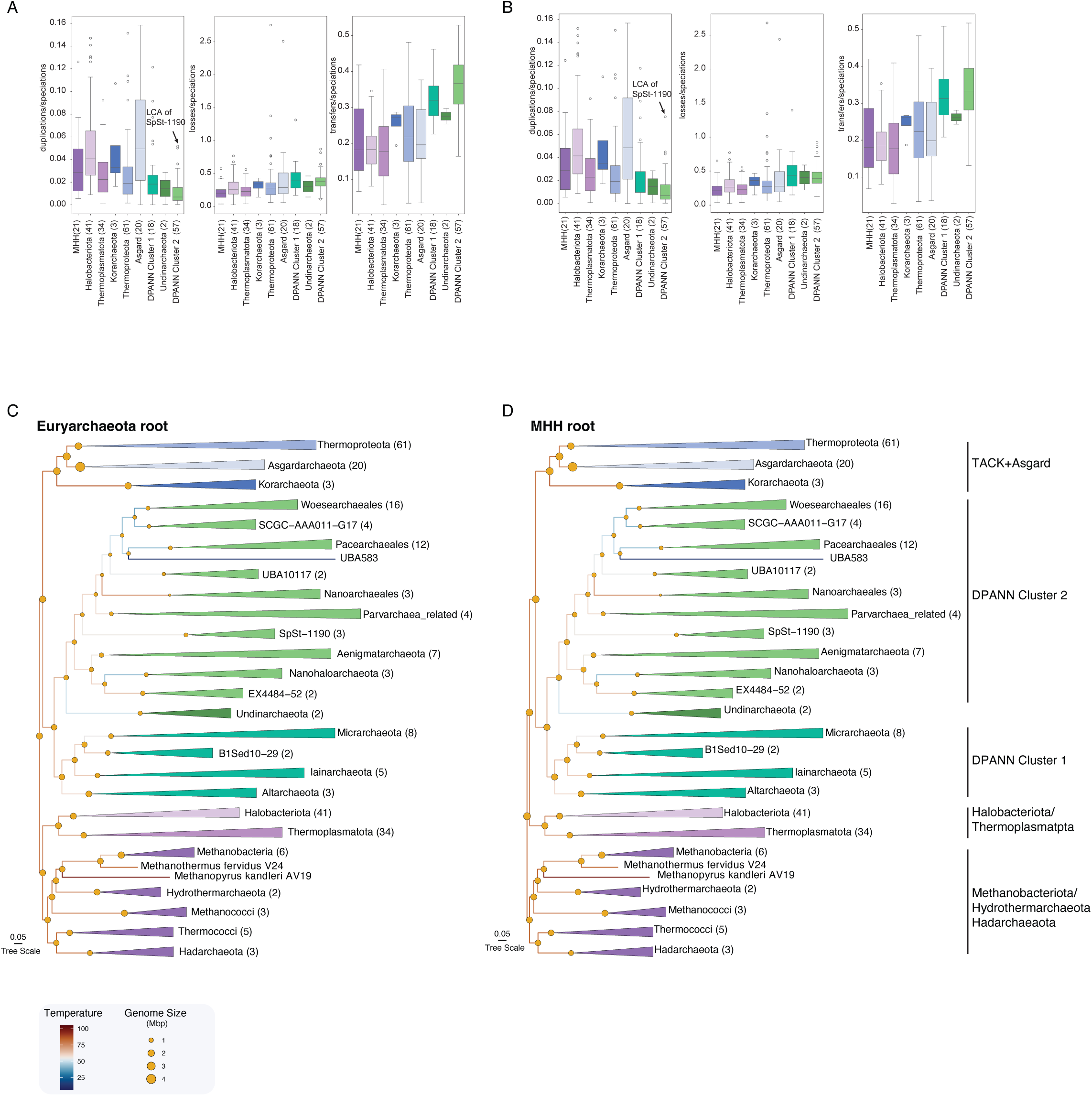
Evolution of gene family repertoires and temperature preferences in the Archaea. A comparison of normalised duplication, loss and transfer events inferred at all branches of different archaeal groups under (A) Euryarchaeota root and (B) MHH root. Genome sizes appear to have remained roughly constant in Euryarchaeota (including Halobacteriota, Thermoplasmatota, Methanobacteriota_A(B), Hydrothermarchaeota and Hadarchaeota) and TACK, expanded in Asgard archaea, and generally reduced in DPANN (including DPANN Cluster 1, Undinarchaeota and DPANN Cluster 2 archaea), with some lineage-specific exceptions. The last common ancestor of Archaea might have been a hyperthermophilic organism, encoding reverse gyrase under both the (C) Euryarchaeota and (D) MHH root hypotheses. The inferred temperature is provided at Table S28. A Kruskal–Wallis rank sum test was performed to assess differences among archaeal groups. Post hoc pairwise comparisons were conducted using the Conover test. Significance results are provided in Table S20-S22. The number in brackets corresponds to the number of included genomes per taxonomic group.

Our analyses inferred significantly higher normalised loss and transfer events in DPANN Cluster 1 and DPANN Cluster 2 archaea compared to euryarchaeal and thermoproteotal lineages (Conoveŕs test, P<0.05, Table S21-22), suggesting that genome evolution of DPANN was shaped by gene exchanges and losses, consistent with their reduced genomes, predicted symbiotic lifestyle and host dependencies. In fact, the branch leading to the common ancestor of DPANN archaea (LDCA) experienced 639.82 /592.64 loss events, followed by another 490.19/588.91 events on the branch leading to Undinarchaeota and DPANN Cluster 2 archaea under the Euryarchaeota/MHH root (Table S19, Fig. S7-8). Notably, while the duplication rate is significantly lower in DPANN Cluster 2 archaea, SpSt-1190 lineages appeared to have the highest normalised duplication events among DPANN Cluster 2 archaea (Fig. 2AB). Additionally, we observed a high number of duplication and acquisition events (transfers and originations) at the branch leading to Haloferacaceae, whose members are suggested to have evolved from a methanogenic ancestor into aerobic/light harvesting heterotrophs^79^ and are reported to be polyploid^80^ with duplicated genes predicted to be involved in the osmostic stress response for adaptation to hypersaline environments (Fig. S7-8).

### The Last Archaeal Common Ancestor was a thermophilic methanogen

Branch-wise reconciliation analysis provides an inference of gene family evolution and an estimate of ancestral genome content across the tree, allowing the reconstruction of the metabolic potential of the last archaeal common ancestor (LACA). Compared to previous work^18^, this reconstruction sampled broader archaeal diversity and made use of a more realistic, better fitting reconciliation model to reconstruct archaeal genome evolution according to the BIC (Table S10). On the basis of the relationship between number of arCOG gene families and genome size in modern archaeal genomes (Asgard archaea: Pearson’s correlation coefficient: 0.967, P=3.27x10^-12^, all other Archaea: 0.9587, P=2.57x10^-130^, Fig. S9, Table S23), we estimated the genome size of LACA to be 1.58 Mb (95% bootstrap confidence interval, BCI: 1.54 Mb to 1.61 Mb) for the Euryarchaeota root and 2.06 Mb (95% BCI: 2.01 Mb to 2.11 Mb) for the MHH root (Fig. 2CD). Based on the inferred presence probabilities (PP) of all arCOG families, we further predicted that LACA encoded between 800 to 1219 arCOG gene families for the Euryarchaeota root, and 1190 to 1709 gene families for the MHH root (Table S24-25). This estimate is obtained by integrating over the LACA PPs for all gene families: some gene families have LACA PPs equal to zero, suggesting that they originated later in archaeal evolution (e.g., ammonia monooxygenase subunit A, arCOG08676; for succinctness, PP values of Euryarchaeota/MHH root are reported as x/y unless specified otherwise; here, 0/0); others have moderate root PPs (≥ 0.5), suggesting they may have been present early in archaeal evolution (e.g. glucokinase, arCOG04280 with PP = 0.54/0.54), while others have high root PPs (≥ 0.75), representing strong evidence that the gene was already present in LACA (e.g. DNA-directed RNA polymerase subunit A, arCOG04527, PP = 0.99/1). Due to phylogenetic uncertainty, stochastic error, and the mosaic ancestry of genomes, PP support for the constituent genes of a metabolic pathway is often variable. For example, while most canonical genes encoding proteins involved in replication^81,82^ (Fig. S10, Table S24-25) and translation machinery^83^ (Fig. S11, Table S24-25) were inferred to having been present in LACA, some had low support. This incompleteness of certain modern metabolic pathways is expected, given the evolutionary changes through time and limitations of our analysis pipeline including phylogenetic noise. Hence, we interpret our reconstruction of LACA’s metabolism based on the inference of pathways using the same PP cutoff (≥ 0.5) as in previous analyses^12,18,63^.

The inferred gene content of LACA suggests that it already had most of the components of modern archaeal DNA replication, transcription, translation and protein quality control machinery in agreement with previous work^18^ (Fig. S10-S14). We also inferred an FtsZ-based cell division system, components of secretion systems, and various transporter genes (Table S24-25). Additionally, we identified most of the proteins required for archaellum biosynthesis^84^ and an archaeal/vacuolar-type ATP synthase (Table S24-25, Fig. S15). The improvement of the reconciliation model and expanded archaeal diversity furthermore allowed us to map the presence of many additional components to LACA, including proteins involved in the biosynthesis for key cellular building blocks, such as nucleotides and lipids, which have been missed previously^18^. For example, a complete mevalonate pathway (MVA) via anhydromevalonate 5-phosphate along with the genes for archaeol biosynthesis, including glycerol-1-phosphate dehydrogenase (GldA, arCOG00982, PP=1/0.99) (Fig. 3, Fig. S15, Table S24-25), were mapped back to LACA. The improvement also included many genes in *de novo* purine and pyrimidine and cofactor biosynthesis (See Supplementary Material).

**Fig. 3.**
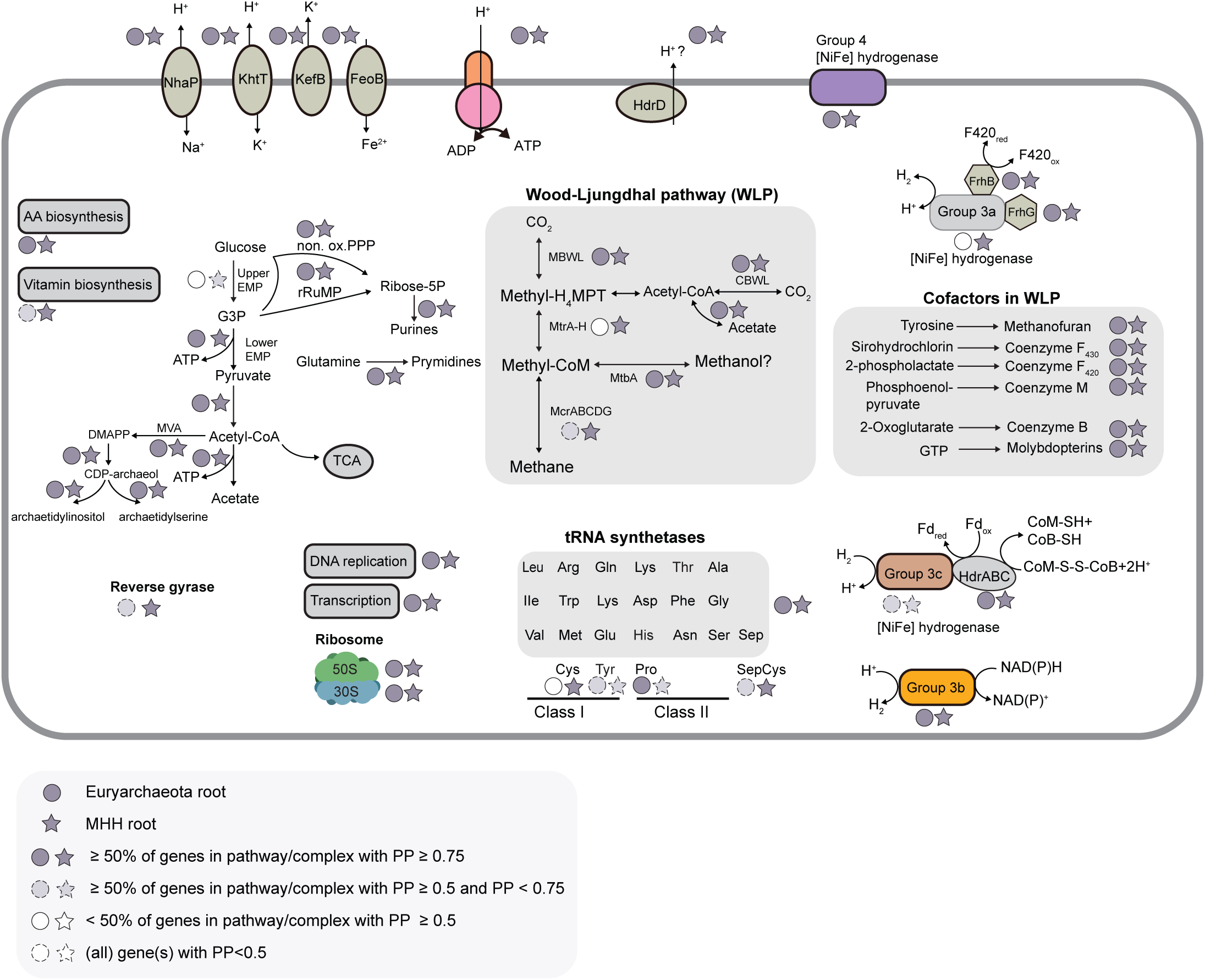
Ancestral reconstruction of the last archaeal common ancestor (LACA) under the two most plausible root hypotheses. The reconstruction is based on genes that are mapped to LACA with a presence posterior probability (PP) ≥ 0.5 under the two most plausi-ble root positions. Pathway is shown by arrows, and complexes/genes by boxes. We recovered strong support for methyl-coenzyme M reductase in LACA, along with the carbonyl- and methyl-branch of the Wood-Ljungdhal pathway, suggesting LACA was a methano-gen. Posterior probability for genes used in reconstruction is provided in Table S24-27. A more detailed figure is provided in Fig. S15 and Fig. S16. MBWL: methyl branch of Wood-Ljungdahl pathway; CBWL: carbonyl branch of Wood-Ljungdahl pathway. The complete-ness of amino acid and vitamin biosynthesis genes are counted from gene families that are inferred to have a presence probability (PP ≥ 0.5) in at least two ancestors (Table S24-25).

Further, we identified almost all components of the lower glycolytic pathway, non-oxidative pentose phosphate pathway and ribulose monophosphate with high PP support (Fig. 3, Fig. S15, Table S24-25). Several genes in the tricarboxylic acid (TCA) cycle were also mapped to LACA. (Fig. S15, Table S24-25). Among the various different carbon fixation pathways operating in archaea^85–89^, the Wood-Ljungdahl pathway (WLP) has been suggested to represent an ancient metabolic pathway^18,90,91^ potentially tracing back to the last universal common ancestor^63,92,93^. In line with this, our results traced almost all components of the tetrahydromethanopterin methyl- and carbonyl-branch of the WLP to LACA (Fig. 3, Fig. S16, Table S24-25). Our reconstructions also indicated the presence of nearly all subunits of the key enzyme of methanogenesis/methane oxidation, i.e. methyl(acyl)-coenzyme M reductase (M(A)cr) (McrA, arCOG04857, PP=0.5/0.58, McrB, arCOG04860, PP=0.61/0.56, McrG, arCOG04858, PP=0.39/0.82, McrD, arCOG04859, PP=0.78/0.96, and McrC, arCOG03225, PP=0.74/0.94) (Fig. 3, Table S24-25) in LACA.

Moreover, we identified many methanogenesis marker proteins and genes involved in methanofuran, coenzyme F_430_, coenzyme F_420_, coenzyme B, coenzyme M, and molybdopterins biosynthesis in LACA (Fig. 3, Fig. S16, Table S24-25). This provides the first support for a methanogenic LACA based on ancestral genome reconstructions, in agreement with evidence from the geological record^94^. LACA was furthermore inferred to have encoded two out of eight subunits of the tetrahydromethanopterin S-methyltransferase (Mtr) complex (MtrA, arCOG03221, PP=0.92 and MtrH, arCOG04336, PP=0.98) under the Euryarchaeota root (Fig. 3, Fig. S16, Table S24), and six out of eight subunits under the MHH root (MtrA, arCOG03221, PP=0.99; MtrB, arCOG04867, PP=0.65; MtrC, arCOG04868, PP=0.58; MtrE, arCOG04870, PP= 0.6; MtrG, arCOG03380, PP=0.71; MtrH, arCOG04336, PP=0.97) (Fig. 3, Fig. S16, Table S25). This enzyme is used by many hydrogentrophic methanogens for sodium ion translocation^40,95^. High support for the presence of methanol-corrinoid protein: coenzyme M methyltransferase (MtbA, arCOG03325, PP=0.96/0.94 and methanogenic corrinoid protein (MtbC, arCOG02028, PP=0.77/0.99) at the root suggest that LACA might have been able to use methylated compounds for methanogenesis. In addition, acetate-CoA ligase (AcdA, arCOG01340, PP=0.60/0.85) and acetyl-CoA synthetase (Acs, arCOG01529, PP=0.94/0.97) were also recovered at the root, indicating the potential to metabolise acetate. Thus, our analyses indicate the antiquity of the WLP and methanogenesis in Archaea^39,96,97^, with the potential for CO_2_-reducing methanogenesis being root-dependent.

Our reconstruction indicated high PP support for a group 3 [NiFe]-hydrogenase (PP=0.98/0.99) (Table S26-27, Fig. S16, Fig. S17, See Supplementary Material), among which we were able to map a group 3c [NiFe]-hydrogenase (PP=0.66/0.68) (Table S26-27) and heterodisulphide subunits (HdrA, arCOG02236, PP=0.73/0.81; HdrC, arCOG00964, PP=0.87/0.99) to LACA with at least moderate support (Table S24-25). This suggests that LACA might have had the ability to use a group 3c [NiFe]-hydrogenase to bifurcate electrons from H_2_ to reduce ferredoxin and heterodisulphide of coenzyme B and M, coupled with the Mcr complex to form methane. Furthermore, our analyses indicated that an ancestral group 4 [NiFe] hydrogenase might have been present in LACA (PP=0.9/0.99; Table S26-27, Fig S16). Group 4 [NiFe]-hydrogenases comprise various respiratory membrane-bound enzymes in modern organisms and may have been involved in energy conservation in the last universal common ancestor^98,99^. These results are consistent with the view that [NiFe]-hydrogenases are ancient enzymes^99^. Additionally, we recovered a subunit of heterodisulfide reductase (HdrD, arCOG00333, PP=0.99/0.97), which is present in various modern archaea, including methanogens^100,101^, and might facilitate electron transfers.

Finally, we estimated the optimal growth temperature of LACA based on the observation that the amino acid composition of our 45-gene supermatrix showed a significant correlation with the optimal growth temperature (Pearson R: -0.87, P=8.951x10^-20^) of modern Archaea (Fig. S18). We used the branch-heterogenous CoaLA substitution model^102^ to sample 100 ancestral sequences reconstructed at nodes of the trees and inferred the temperature optima for each node^18^. The analysis suggested that LACA may have been a (hyper)-thermophile (median 78.78℃, 95% CI: 74.15 ℃ - 83.52 ℃ under the Euryarchaeota root; median 79.40 ℃, 95% CI: 74.18 ℃ - 84.30 ℃ under the MHH root; Table S28), consistent with previous work^18,102^ and with the inference that reverse gyrase, a specific hallmark of (hyper)-thermophiles, may have been present in LACA under the Euryarchaeota/MHH roots (arCOG01526, PP=0.68/PP=0.75, Table S24-25).

### The DPANN ancestor was a free-living organism

Our rooted species tree resolves DPANN as a monophyletic sister clade of TACK and Asgard. The last DPANN common ancestor (LDCA) under the Euryarchaeota root and MHH root, respectively, was estimated to have encoded a genome of 1.32 Mb/1.68 Mb (95% CI: 1.28 Mb to 1.36Mb / 95% CI: 1.64 Mb to 1.72 Mb), with 546-827/820-1152 arCOG families (Table S23). This is comparable to modern Altiarchaeota genome sizes and gene repertoires (1.40 Mp to 2.48 Mp; 1691 to 2841 total coding sequences) (Table S1), suggesting that the LDCA may have been a free-living organism similar to modern Altiarchaeota^27,103^. Indeed, our inference of the ancestral metabolic content of the LDCA is compatible with a free-living lifestyle. For instance, our analyses indicate the presence of almost all components of archaeal ether lipid biosynthesis, and *de novo* purine and pyrimidine biosynthesis pathways in the LDCA (Fig. 4, Table S24-25). A number of genes involved in vitamin and amino acid biosynthesis were also recovered (Fig. 4, Table S24-25). This contrasts with many DPANN Cluster 2 representatives including co-cultivated symbionts such as members of Nanoarchaeales^34–36,104^ like *Nanoarchaeum equitans*^29,105^, and members of Nanohaloarchaeota^30,31^, such as *Nanohaloarchaeum antarcticus*^30^, that do not encode lipid, nucleotide, amino acid or cofactor biosynthesis pathways. We also identified components of central carbon metabolic pathways, including the lower glycolytic pathway, pentose phosphate pathway (PPP) and TCA cycle in the LDCA (Fig. S15). Notably, although the M(A)cr complex was lost in LDCA and the last DPANN Cluster 1 common ancestor (LDC1CA), both ancestors were predicted to have encoded many genes in the H_4_MPT methyl and carbonyl branch of WLP as well as an acetate-CoA synthetase (Fig. S16). However, genes of the WLP pathway were lost on the branch leading to the last common ancestor of Undinarchaeota+DPANN Cluster 2 archaea and are absent in most modern DPANN archaea except Altiarchaeota and SpSt-1190^27,106^.

**Fig. 4.**
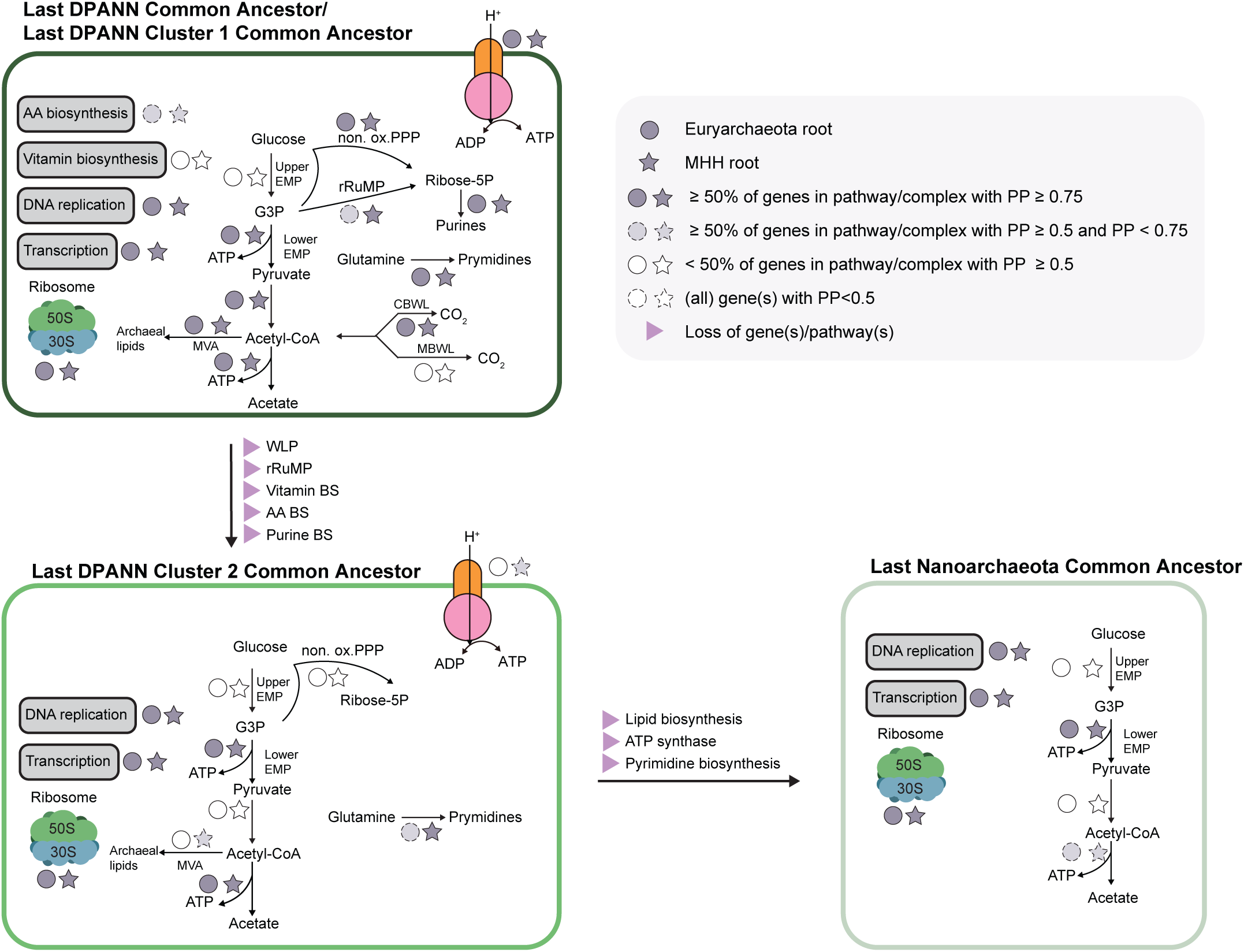
Ancestral reconstruction of the last common ancestor of DPANN archaea, DPANN Cluster 1 archaea, DPANN Cluster 2 archaea, and Nanoarchaeota. The reconstruction is based on genes that could be mapped to these ancestors with a presence posterior probability (PP ≥ 0.5). Pathway is shown by arrows, and complexes/genes by boxes. We recovered strong support for the carbon-yl- and methyl-branches of the Wood-Ljungdahl pathway (WLP), in addition to lipid-, vitamin- and amino acid-biosynthesis genes in the last common ancestor of DPANN and the last common ancestor of DPANN Cluster 1, suggesting a free-living lifestyle. However, genes related to the WLP, ribulose monophosphate pathway, vitamin and amino acid biosynthesis pathways were lost in the DPANN Cluster 2 archaea, followed by a loss of genes comprising the ATP synthase, and lipid and nucleotide biosynthesis genes in the last common ancestor of Nanoarchaeota ancestor, increasing their host-dependency (Supplementary Material). Posterior probability for genes used in reconstruction is provided in Table S24-27.

We inferred an ongoing trend of convergent genome reduction during the evolution of DPANN archaea (Fig. 2CD, Fig. S19, Table S23). In particular, DPANN Cluster 2 archaea are characterised by genome streamlining that appears to have proceeded in parallel in different lineages^62,107^. Specifically, our reconstructions inferred a reduction in genome size from 1.32/1.68 Mb in LDCA to 0.92/1.26 Mb in the DPANN Cluster 2 common ancestor (LDC2CA, Table S23). The reduction in genome content included genes encoding enzymes of the WLP, ribulose monophosphate pathway (RuMP), vitamins, amino acids, nucleotides and lipid biosynthesis pathways, which indicates dependency on external sources likely evolved by the LDC2CA (Fig. 4, Fig. S15, Table S24-25, Supplementary Material). However, we identified most genes of the lower glycolytic pathway and acetate-CoA ligase in the LDC2CA, in line with experimental support indicating that this machinery may be indispensable for these symbionts in the absence of host-derived ATP^108^. We also inferred that most of the archaellum subunits^84^ of the motor complex (ArlHIJ) were present in the LDC2CA, which were found in extant representatives of DPANN Cluster 2 archaea (Table S24-S25)^108,109^. Similarly, we identified most of the components of DNA replication, transcription and translation machinery in LDC2CA and last common ancestor of Nanoarchaeota, in line with extant representatives^24,25,110,111^. In contrast to this general trend, we inferred that some clades within Woesearchaeales and Pacearchaeales experienced genome expansion (Fig. S19).

For example, we inferred an increase of 0.34 to 0.39 Mb of one Woesearchaeales clade, and an increase of 0.26 to 0.27 Mb in one Pacearchaeales clade, compared to the last common ancestor of Woesearchaeales and Pacearchaeales, respectively (Fig. S19, Table S23). The genome content expansion of the Woesearchaeales clade includes genes involved in amino acid transport and metabolism, as well as energy production and conversion, in line with a previous inference^107^. For example, several subunits (1-4) of the archaeal/vacuolar-type ATP synthase were inferred to have been re-acquired during this expansion (Table S29). In contrast, the majority of genes that contributed to genome expansion in Pacearchaeales were inferred to be involved in translation, transcription and replication machinery (Table S30). For example, acquisitions of transcriptional regulators were inferred under both the Euryarchaeota and MHH root (Table S30) for the latter lineage.

## Discussion

The archaeal phylogeny continues to be revisited and revised as genomes from new groups become available, and phylogenomic methods improve^1,3,5,6,73,112,113^. While many studies consistently support the clanhood of DPANN and TACK+Asgard archaea^13,14,18,19,24,25,39,63^, the clanhood of Euryarchaeota remains debated. Some analyses recover Euryarchaeota as a monophyletic group^14,18,24,25,39,63,112^, whereas others suggest it to be paraphyletic, with (hyper-)thermophilic lineages, Thermococci and/or Hadarchaeota, branching basal to TACK+Asgard^17,19,49,114^. These two lineages were proven difficult to place in previous work^13,14,17,114–116^. Our analyses consistently recovered the clanhood of Euryarchaeota under the progressive removal of compositionally biased sites, in line with a previous observation in which support for the clanhood of Euryarchaeota emerged upon site filtering^17^. While some studies placed Altiarchaeota at the base of all DPANN archaea^17,19,49^, our analyses supported the placement of Altiarchaeota within DPANN as a member of DPANN Cluster 1 once gene families strongly affected by horizontal evolution (including interdomain transfer) were removed, suggesting that alternative placement was induced by including non-vertical markers^62^.

We applied various methods to root the archaeal tree and assessed confidence of root placements^52^. Our results reveal that outgroup rooting and non-reversible models appear to be unable to distinguish among a broad set of potential root placements (Table S3, S6). By contrast, the GFmix model, which accounts for variation in amino acid frequencies across sites and branches, supported a Halobacteriota+Thermoplasmatota root, in line with recent analyses^19,23^. While these analyses are typically based on information from a small set of marker genes, phylogenetic reconciliation allows to incorporate additional information based on gene evolutionary histories^12,18,58,72,117^. Earlier attempts to resolve the root of Archaea using gene tree-species tree reconciliation approaches were based on a smaller taxon set and relied upon a simpler reconciliation model that did not incorporate variation in duplication, transfer and loss rates between different lineages^18^ - a limitation that can induce artifacts, especially when including a diverse set of reduced-genome from symbionts. Analyses using a better-fitting model placed the archaeal root on or within Euryarchaeota, with DPANN emerging as a sister group to TACK+Asgard. The root recovered in the GFmix analysis was one node away from these two adjacent branches, defining a small “root region” on the archaeal tree at or near the base of the Euryarchaeota. These results emphasise the importance of integrating realistic models of evolution that improve model fit and resolve deep archaeal phylogeny.

Our reconciliation approach simultaneously allowed reconstructing ancestral gene sets along the rooted archaeal tree, resolving a relatively complex LACA that was likely a (hyper-)thermophilic methanogen^18,81–83,118,119^. The inferred metabolic content of LACA is consistent with methyltrophic methanogenesis or potentially CO_2_-reducing methanogenesis^39,91,96,120^. We hypothesize that such a hyperthermophilic ancestor could have thrived in environments analogous to hydrothermal vents or geothermal subsurface systems with strong reducing conditions abundant in H_2_, CO_2_, and methylated compounds such as methanol^121,122^.

While various studies have recovered a deep split between DPANN and all other archaeal lineages^14,18,24^, more recent work provided support for DPANN forming a derived lineage within Euryarchaeota in a tree rooted between TACK + Asgard and all other archaea^17^. Our analyses also recovered DPANN as a derived lineage, albeit branching sister to the TACK + Asgard (Fig 1). These results suggest that the symbiotic lifestyle of DPANN archaea is a secondary adaptation rather than an ancestral trait. In fact, our analysis suggests the LDCA might have been a free-living organism, with the subsequent diversification of DPANN lineages having been shaped by high rates of gene loss and transfer. While genome reduction in both DPANN and CPR can thus be considered a derived trait, their evolutionary trajectories differ. Reduction in CPR likely occurred early on the stem lineage^12^, whereas DPANN gene loss was more episodic and, in part, occurred convergently in different lineages. However, genome reduction in the stem leading to DPANN Cluster 2 exhibits similar patterns as observed for CPR, involving the loss of genes coding for proteins involved in central carbon and energy metabolism including the ATP synthase and amino acid biosynthesis^12^. The loss of metabolic pathways also shows a common pattern in reductive genome evolution in the bacterial endosymbionts of insects^24,25,123^. In contrast to those endosymbionts, DPANN archaea appear to retain genes for proteins that are involved in DNA repair and transcription. This difference may reflect their distinct symbiotic strategies: all DPANN symbionts that have been successfully co-cultivated have episymbiotic stages^29–31,33,108,124^. Therefore, they likely experience stronger selection to maintain key informational machinery than strictly vertically inherited intracellular endosymbionts. Furthermore, the ectosymbiotic lifestyle of DPANN archaea might explain the observed genome expansion in certain DPANN lineages, such as Woesearchaeales and Pacearchaeales, through high rates of horizontal gene transfer. The prospective analysis of the sources and mechanisms of these gene transfers may allow identifying host organisms for the vast diversity of uncultivated DPANN.

## Methods

### Taxa selection

Contigs and protein files of archaeal genomes were downloaded from NCBI in August 2021 (n = 7,967) . If protein files were unavailable, gene calling was performed using Prokka (v1.14, settings: --kingdom Archaea --addgenes --increment 10 --compliant --centre UU --norrna --notrna)^125^. Taxonomic classifications were assigned to all genomes using GTDB-Tk using GTDB release 202^126^ and the original NCBI tax ID was converted into a NCBI taxonomy string using ncbitax2lin (https://github.com/zyxue/ncbitax2lin). Genome completeness was estimated using CheckM2 (v0.1.3)^127^ and the output was used to calculate the missing fraction per genome as an input for the ancestral reconciliations.

Genomes with completeness >50% and contamination <5% were used for an initial phylogenetic analysis using the PhyloSift marker set^128^. In brief, homologs of 34 PhyloSift marker proteins were identified in each genome using the PhyloSift v1.0.1 ‘find’ mode (settings: –besthit). Sequences were aligned with MAFFT (v7.407, settings: –reorder)^129^, trimmed with BMGE (v1.12, settings: -t AA -m BLOSUM30 -b 2 -h 0.55)^130^. The trimmed alignments were concatenated using catfasta2phyml.pl (https://github.com/nylander/catfasta2phyml) and the concatenated protein alignment was used to reconstruct a phylogenetic tree using IQ-TREE (v1.6.7, settings: -m LG+G -nt AUTO -bb 1000 -alrt 1000)^131^. Based on this tree, a sub-selection of 1,711 archaeal reference genomes were selected based on high genome completeness, low contamination, and taxonomic diversity.

To further thin out the taxa selection, two rounds of species tree inference were performed using 51 universal, single-copy marker proteins suitable for archaeal phylogenetic analyses^62^. Specifically, a customized TIGR database was queried against a database generated from all archaeal proteins using hmmsearch (v3.1b2)^132,133^. The output was parsed to only retain hits with a e-value ≤ 1E-3 and then to only include the best-hit per protein selected using lowest e-value and highest bit score. Selected sequences were aligned with MAFFT (v7.407, settings: –reorder)^129^ and alignments were trimmed with BMGE (v1.12, settings: -t AA -m BLOSUM30 -b 2 -h 0.55)^130^. Individual alignments were concatenated using catfasta2phyml.pl and a phylogenetic tree was generated using IQ-TREE (v1.6.7, settings: -m LG+C20+F+R -bb 1000 -alrt 1000)^131^ and visualized using FigTree (v1.4.4). The tree was manually inspected and an additional subset of taxa was selected based on genome completeness and contamination statistics while maintaining a taxonomically diverse set of genomes with a specific focus on the DPANN archaea. This workflow was repeated resulting in a selection of 513 taxa that were further reduced to the focal taxa set of 257 genomes with balanced taxa representation of Euryarchaeota, TACK+Asgard and DPANN (Table S1).

### Genome annotations

For further functional annotation, proteins from 257 archaeal genomes were compared against several databases, including the COGs (using the 2020 update)^134^, arCOGs (version from 2020)^135^, the KO profiles from the KEGG Automatic Annotation Server (KAAS; downloaded April 2019)^136^, the Pfam database (Release 31.0)^137^, and the TIGRFAM database (Release 15.0)^132^. Individual database searches were conducted as follows by querying these databases against all archaeal genomes using hmmsearch v3.1b2 (settings: --tblout --domtblout --notextw)^133^. The output was parsed to only include the best hit for each protein using the best e-value and bit score value. Additionally, all proteins were scanned for protein domains using InterProScan (v5.29-68.0; settings: --iprlookup --goterms)^138^.

### Inferring an unrooted archaeal species tree

The focal taxa selection was used to infer a maximum likelihood phylogeny using 51 universal, single-copy marker proteins as described above but with minor adjustments to improve the quality of the species tree^62^- i.e. markers with more than 10% duplicated proteins were discarded leaving 45 markers. Sequences were aligned using MAFFT or MAFFT-L-INS-i for sequences more than or less than 1000 sequences, respectively, and trimmed with BMGE. Single gene trees were generated with IQ-TREE 2 (v2.1.2, settings: -m LG -T AUTO -keep-ident --threads-max 2 -B 1000 -bnni)^64^. A custom script (sequence_correler.py, https://zenodo.org/records/17360806) was run to identify multi-copy proteins^15^. Each single gene tree was visually inspected to identify the correct protein sequences per marker and remove long branching sequences. After removing problematic sequences, trimmed alignments were generated using the procedure described above and concatenated using catfasta2phyml.pl (https://github.com/nylander/catfasta2phyml). The best fitting phylogenetic model was estimated using IQ-TREE 2 (v2.1.2, settings: -m MF -mset LG -madd LG+C10,LG+C10+G,LG+C10+R,LG+C10+F,LG+C10+R+F,LG +C10+G+F,LG+C20,LG+C20+G,LG+C20+F,LG+C20+ G+F,LG+C20+R,LG+C20+R+F,LG+C30,LG+C30+G,LG+C30 +R,LG+C30+F,LG+C30+R+F,LG+C30+G+F,LG+C40,LG+C40+G,LG+C40+R,LG+C40+F,LG+C40+R+F,LG+C40+G +F,LG+C50,LG+C50+G,LG+C50+R,LG+C50+F,LG+C50+ R+F,LG+C50+G+F,LG+C60,LG+C60+G,LG+C60+R,LG +C60+F,LG+C60+R+F,LG+C60+G+F --score-diff all), selected based on Bayesian Information Criterion^139^, and a phylogenetic tree was inferred using IQ-TREE 2 (v2.1.2, settings: -m LG+C60+F+R -B 1000 -alrt 1000)^64^ and visualized using FigTree (v1.4.4). The species tree was re-rooted at 15 different key positions (positions from literature and between each major lineages) using a custom script (root_w_ete3.py, https://zenodo.org/ records/17360806).

To assess the clade support for major archaeal lineages, compositionally biased sites were filtered from the alignment in a progressive manner (from 10-80%, in 10% increments) using alignment_pruner.pl (https://github.com/novigit/davinciCode/blob/master/perl/alignment_pruner.pl; settings: --chi2_prune). For each pruned alignment a phylogenetic tree was inferred with IQ-TREE 2 (v2.1.2, settings: -m LG+C60+F+R -bb 1000 -alrt 1000)^64^. All tree topologies were similar overall so we used the phylogenetic tree based on the full alignment for all further analyses (Fig. S2).

### Reciprocal outgroup rooting of an archaeal species tree with bacteria

To perform outgroup rooting of archaea, we selected 25 vertically-evolving marker proteins shared between bacteria and archaea^13^ (Table S2). COG marker sequences were identified using hmmsearch v3.1.2 from 257 archaeal and 103 bacterial genomes. Alignments were generated using MAFFT-L-INS-i and trimmed using BMGE (-m BLOSUM30 -h 0.55 -t AA). Single gene trees were inferred based on LG+G using IQ-TREE 2 (v2.1.2) and inspected to remove long branching sequences and false paralogues. Archaeal and bacterial sequences were then realigned and trimmed separately as described above to make a concatenation. These two concatenations were merged using MAFFT-L-INS-i (--merge) options. This procedure was chosen because it retained 2,122 additional alignment sites relative to aligning all sequences jointly followed by trimming. Finally, we inferred a best-known ML tree under LG+C60+F+R model which recovered a DPANN root (Fig. S1). Subsequently, we generated 14 additional root hypotheses by relocating the outgroup attachment point to the archaeal ingroup (Table S3), and conducted maximum likelihood based tree inferences under these topological constraints under LG+C60+F+R using IQ-TREE 2 (v2.1.2) (-g root -wlsr). AU tests were performed using CONSEL v0.2^140^ using 1,000,000 bootstrap samples.

### Rooting of archaeal species tree using a non-reversible model

Additionally, we inferred a maximum likelihood phylogeny from the same concatenated alignment using a non-reversible model implemented in IQ-TREE 2 (v2.3.6, setting: -m NONREV+G). The likelihood of trees being rooted on every branch of the ML tree was evaluated using the –root-test option and AU tests were performed for those trees using the option: -zb 10000 -au. P values of the AU test were shown in Fig. S4. The results representing the 15 root positions that were tested with outgroup-rooting and reconciliation analysis were provided in Table S6.

### Rooting of archaeal species tree using a non-stationary model

We also used the GFmix model (v1.2)^17,59,60^ with LG+C60+F+G model to assess the 15 different rooting hypotheses based on the same concatenated alignment. Site-wise likelihood was generated using the: -l options, and AU tests were performed using CONSEL v0.2^140^ using 1,000,000 bootstrap samples. The GFmix rooting analyses were also performed the alignment and treefile from a recent study^17^ (settings: -d -l). Afterwards, an AU test was performed.

### Gene tree/species tree reconciliations to root archaeal species tree and reconstruct ancestral proteins

#### arCOG gene families for ancestral gene content reconstruction

Gene families for ancestral reconciliations were generated based on the arCOGs. To this end, proteins from an initial taxa set with 513 archaeal genomes (relatively large representation of DPANN) were combined in a protein database that was used in an hmmsearch against the arCOG database (settings: --tblout --domtblout --notextw)^134,135^. Hits with e-values above 1e-3 were discarded from the domain results table and the output further parsed to only include the best hit for each protein based on the lowest e-value and highest bit score.

To identify and split potentially fused proteins (from now on called ‘splits’), proteins were further investigated for secondary domain hits after subtracting the position of the first domain hit from the full protein using bedtools subtract (v2.26.0)^141^. Secondary domains were only considered if they were assigned to a different arCOG than the first domain. This process was repeated until proteins were investigated for up to four domains and the original protein was split into individual domains using the positional domain information with bedtools getfasta. For reference, unsplit proteins were given the extension ‘a0’ and the split proteins were given the extension ‘a1’ to ‘a4’, indicating the primary to quarterly domain, in the generated trees.

All protein sequences were combined and sequences with unambiguous amino acids identified and discarded using a custom python script (remove_seq_with_specific_char.py, https://zenodo.org/records/17360806). Afterwards, a first round of alignments was generated using MAFFT or MAFFT-L-INS-i for sequences more than or less than 1000 sequences, respectively (v7.407, settings: --anysymbol --reorder)^129^. Alignments were trimmed using trimAl (v1.4.rev15, settings: -gappyout)^142^. Sequences with equal to or greater than 50% of gaps were identified and discarded using custom python scripts (faa_drop.py and screen_list_new.pl, https://zenodo.org/records/17360806). Only clusters with at least four sequences were considered for further analysis. Based on these initial selections, a second round of alignments were generated with MAFFT and MAFFT-LINS-i as described above and sequences trimmed with BMGE (v1.12, settings: -t AA -m BLOSUM30 -b 2 -h 0.55)^130^. The trimmed alignments were used to generate a first round of single gene trees using IQ-TREE 2 (v2.1.2, settings: -m LG+G -T 1)^64^. Single sequences or clusters of sequences on long branches were identified using a custom script and using a branch length cutoff of two (cut_gene_tree.py, https://zenodo.org/records/17360806)^15^. The validity of this cutoff was confirmed on basis of a random number of single gene trees that were manually inspected after adding the KO mapping, PFAM and arCOG mappings as well as taxa information to the tip labels. Single sequences on long branches were discarded and clusters of sequences of at least four sequences were separated into a separate cluster using a custom script (screen_list_new.pl, https://zenodo.org/records/17360806). Finally, sequences within each cluster were subset to the focal taxa set of 257 archaeal genomes with balanced representation of Euryarchaeota, TACK+Asgard and DPANN lineages, amenable to complex gene-tree and species-tree reconciliation models (https://zenodo.org/records/17360806).

A third round of alignments was generated with MAFFT and/or MAFFT-LINS-i and trimmed with BMGE as described above. The trimmed alignment was used to run a model test with IQ-TREE 2 (v2.1.2, settings: -m MF -mset LG -madd LG+C10, LG+C10+G, LG+C10+R, LG+C10+F, LG+C10+R+F, LG+C10+G+F, LG+C20, LG+C20+G, LG+C20+F, LG+C20+G+F, LG+C20+R,LG+C20+R+F --score-diff all) as well as guide trees (settings: -m LG+G+F --score-diff all -nt 1 -wbtl -B 1000 -bnni). Depending on the best model chosen, single gene trees were run using either with posterior mean site frequency (PMSF) model (settings: -s trimmed_aln.faa -m best_model -ft guide_tree.treefile -T 1 -wbtl -B 1000 -pers 0.2 -nstop 500)^143^ or the the default model settings of IQ-TREE 2 (v2.1.2, settings: -s trimmed_aln.faa -m best_model -T 1 -wbtl -B 1000 -pers 0.2 -nstop 500) depending on whether or not a C-series profile mixture model was selected as the best model, respectively. In total 7,059 single gene trees were generated for the arCOG-based clusters, respectively, and used as input for the ancestral reconciliations as described below.

#### Gene tree-species tree reconciliation

To account for the incomplete nature of some MAGs, genome completeness estimates from CheckM v2 were used in reconciliation^127^ (Table S1). Bootstrapped single gene trees, inferred from 7,059 arCOG-based clusters (see arCOG gene families for ancestral gene content reconstruction), were reconciled with 15 different rooted archaeal species trees (Figure 1A) using the distinct reconciliation models implemented with AleRax v1.1.1^57^: These models include a global rate model (3 parameters, one D, one T and one L homogeneous to all branches), a per-family DTL rate model (21177 parameters, one D, one T and one L per gene family), and different branch-wise DTL models to tackle differences in genome-evolving rates across archaeal lineages via the --model-parametrization option. All optimisations were performed using limited-memory BFGS algorithm following recommendations from Williams et al., 2024^72^. We built a series of branch-heterogeneous gene family reconciliation models on top of a global-rate model as baseline (Table S31). First, we introduced DPANN-specific rate classes: a different loss rate (DPANN:L; 4 parameters), different duplication+loss rates (DPANN:DL; 5 parameters), different transfer+loss rates (DPANN:TL; 5 parameters), and different duplication+transfer+loss rates (DPANN:DTL; 6 parameters). We then extended the DTL heterogeneity to major clades—Euryarchaeota, TACK+Asgard, DPANN, and three DPANN sublineages (Woesearchaeales+SCGC-AAA011-G17, Pacearchaeales, Micrarchaeota), totaling in 21 parameters (DTL_br1). Building on this, we allowed clade-specific origination rates for DPANN, “Euryarchaeota”, and TACK+Asgard (25 parameters, DTL_br1_O). Next, we let D, T, and L vary freely for “Euryarchaeota” lineages (i.e. for each Halobacteriota; Thermoplasmatota; and a clade comprising Methanobacteriota_A(B), Hydrothermarchaeota, and Hadarchaeota), for TACK+Asgard lineages (i.e. for each Korarchaeota, Thermoproteota, Asgardarchaeota), and for DPANN lineages (i.e. for each Micrarchaeota; B1Sed10-29; Iainarchaeota; Altiarchaeota; Undinarchaeota; Aenigmarchaeota; PWEA01; EX4484_52; Nanohaloarchaeota; SpSt-1190; parvarchaeal-related lineages; UBA10117; Pacearchaeales; Woesearchaeales+SCGC-AAA011-G17; Nanoarchaeales; plus two internal nodes representing the last common ancestor of all DPANN and of DPANN Cluster 1 archaea, respectively), yielding 72 parameters in total (DTL_br2). In summary, based on the BIC, the model with 72 parameters provided the best model fit across different roots (Table S10). Likelihoods of 7059 families under all these models were compared and tested using AU tests as implemented in CONSEL v0.20 with 1,000,000 bootstrap samples^140^. In addition, we reanalysed data generated by Williams et al, 2017^18^ using a global model, per-family DTL model, and a branch-wise model allowing Altiarchaeota, Micrarchaeota, Iainarchaeota, Nanohaloarchaeota, Aenigmarchaeota, Parvarchaeota, Nanoarchaeales, Woesearchaeales, Pacearchaeales, Halobacteriota, Thermoplasmatota, Asgardarchaeota, Korarchaeota, Thermoproteota to evolve at a different DTL rate, while we used different DTLO rates for nodes representing LCA of DPANN, “Euryarchaeota”, and TACK+Asgard, respectively (Table S31). For ancestral genome content reconstruction, we allowed heterogeneous origination by estimating independent rates for the root, its two immediate descendants, and the DPANN lineage using --origination OPTIMIZE, based on the best-fitting reconciliation model.

#### Gene family filtration

To evaluate the underlying support for competitive root candidates, we performed gene family filtration similar to Colemean etl., 2021^12^. We ordered gene families by species number and taxonomic bias score (defined as |⅓-%DPANN|+|⅓-%TACK+Asgard|+|⅓-%Euryarchaeota|), which quantifies deviation from equal representation of three archaea groups in a gene family. To assess potential of gene families to induce the SGA, we compared the differences in likelihoods between competing root hypotheses and the best-supported maximum-likelihood root as gene families were sequentially excluded from the analysis (Supplementary Material).

### Ancestral sequence reconstruction and temperature estimates

To estimate the ancestral optimal growth temperature of LACA, we reconstructed ancestral sequences based on the supermatrix using the LG+G4+COaLA[2] model (best-fitting model from LG+G4+COaLA[1-3]^102^ based on AIC and BIC; [x] refers to number of independent axis positions per branch and on root) implemented in Bio++ (v2.3.1)^144^. We sampled 100 sequences from the posterior distribution of ancestral states (i.e., amino acid) at each node to account for reconstruction uncertainty. Correspondence analyses on amino acid compositions were performed using the Prince^145^ package (version 0.13.0) Python3^146^. Axis 2 of amino acid compositions were found to be significantly correlated with the optimal growth temperatures in extant archaeal taxa (Pearson R: -0.87, p-value: 8.951x10^-20^, Fig. S18). A linear regression model was fitted using the OLS function implemented in statsmodels (v 0.13.5)^147^, and the fitted (R^2^: 0.75) regression line was used to predict the OGT for ancestral nodes. Confidence intervals were computed to take into account the uncertainty in the parameter estimates of the regression line.

### Genome size estimation

The numbers of arCOG families present at ancestral nodes of interest were obtained by counting families with posterior probabilities (PP) ≥ 0.5 (moderate support) and ≥ 0.75 (strong support). To estimate ancestral genome size, we used all 7,059 gene families. Specifically, we applied locally weighted scatterplot smoothing (LOESS) to model the relationship between protein number and genome size across extant genomes and predicted ancestral genome sizes from the fitted curve. Prediction uncertainty was quantified by bootstrap resampling (1,000 replicates).

### Statistical tests

Statistical significance of transfer propensity and gene duplication, loss and transfer events of different groups were tested using a Kruskal–Wallis test implemented in scipy.stats (v1.16.2), followed by a post hoc pairwise test for multiple comparisons of mean rank sums (Conoveŕs test) using scikit_posthocs (v 0.11.2) with P values adjusted by Holm–Bonferroni method.

## Data Availability and Code availability

Genomic data, workflows for inferring phylogenies, performing reconciliation, and ancestral gene content reconstructions, and custom scripts are deposited in the repository (https://zenodo.org/records/17360806).

## Acknowledgements

This project has received funding from the European Research Council (ERC) under the European Union’s Horizon 2020 research and innovation programme (grant agreement No. 947317, ASymbEL to A.Sp. and grant agreement No. 714774, GENECLOCKS to G.J.S.). A.St. and N.W. are funded by the EU ERA Chair (HORIZON-WIDERA-2022-TALENTS-01: 2023-2028) program under project ID 101087081 (Comp-Biodiv-GR). Our research was also funded by the John Templeton Foundation (63451 to G.J.Sz., T.A.W. and A.Sp.). The opinions expressed in this publication are those of the authors and do not necessarily reflect the views of the John Templeton Foundation.

## Author contributions

A.Sp. conceived the study. N.D. and W.H. designed experiments and assembled the initial and focal datasets, respectively. N.D. performed annotations and developed the phylogenetic analysis workflow. N.D., T.A.M and W.H. performed the phylogenetic analyses and reconciliations. W.H. performed ancestral reconstructions and rooting analyses. N.W., A.St., T.A.W, and G.J.Sz. contributed novel methods and software development. W.H., A.Sp., T.A.M, T.A.W. and G.J.Sz. interpreted the data. W.H., A.Sp. and T.A.W. wrote and all authors edited and approved the manuscript.

