## Supplementary Material for "Phylogenetic reconciliation supports a methanogenic ancestor of the Archaea and a derived origin for host-associated lineages"

CONTENT

**Phylogenetic placement of the Altiarchaeota** **3**
**Exploration of the reconciliation model and gene family filtration** **4**
**Re-analysis of the dataset from Williams et al., 2017** **6**
**Metabolism of the Last Common Ancestor of Archaea (LACA)** **6**
DNA replication 7
Transcription 8
Translation 8
Protein quality control 10
Central carbon metabolism 10
Wood-Ljungdahl pathway 11
Biosynthesis of cofactors related to WLP 12
Methanogenesis 12
[NiFe] hydrogenases 13
**Biosynthetic potential of the LDCA, LDC1CA, LDC2CA and LNCA.** **14**
**References** **15**
**Supplementary Figures** **22**

### Phylogenetic placement of the Altiarchaeota

Altiarchaeota are members of the DPANN Archaea, but unlike other characterised representatives, they are less genome-reduced and are capable of free-living growth[1]. As a result, their phylogenetic position within DPANN may help to inform our understanding of the early evolution of the group. Interestingly, a recent study[2] recovered Altiarchaeota at the base of DPANN - potentially suggesting they preserve an intermediate state of genome reduction from free-living ancestors to host-associated DPANN - while our tree and several others recover Altiarchaeota at a slightly more derived position, at the base of "Cluster 1" DPANN (Fig. 1)[3–5].

To investigate the cause of these contradictory results, we examined the gene marker set used by Baker et al.[2], specifically the NM126 markers, for their level of verticality and domain monophyly. First, we assigned those markers to the Cluster of Orthologs (COG) database using HMMSEARCH v3.1.2b[6]. We used the COG information of these NM126 to extract bacterial homologs from a bacterial dataset with 487 taxa (Table S5, https://zenodo.org/records/17360806). Bacterial homologs were then added to the corresponding NM marker set, i.e. the archaeal homologs from the original study. Sequences were then aligned using MAFFT-L-INS-i (v7.453)[7], and poorly aligned regions were removed using BMGE (v1.12; settings: -m BLOSUM30 -h 0.55)[8]. Single-gene trees were inferred under the LG+G model in IQ-TREE v2.1.2[9] and manually inspected. This revealed that 34 of these markers violated domain monophyly, i.e. archaeal and bacterial homologs did not form two separate clades, indicating that the evolution of these gene families was affected by interdomain horizontal gene transfer (Table S5). To determine whether these markers in the concatenation of Baker et al.[2] contributed to the conflicting phylogenetic results with regard to the placement of the Altiarchaeota, we also re-generated the trimmed alignments of the NM126 markers for concatenation using the method described in Baker et al. (2025), employing MAFFT-L-INS-i (v7.453) and BMGE (v1.12; settings: -m BLOSUM30 -b 3 -g 0.2 -h 0.5). Next, we inferred a species tree of the 34 markers under LG+C60+G using IQ-TREE v2.1.2[9] with mixture weights optimised using –mwopt. We also concatenated the remaining 89 markers that did not violate domain monophyly to infer a species tree using the same approach. Additionally, we inferred single-gene trees for the original NM126 markers under the best-fitting models (settings: -m MF -mset LG -madd LG+C10+G, LG+C20+G, LG+C30+G, LG+C40+G, LG+C50+G, LG+C60+G --score-diff all -cmax 4) with mixture weights optimised using --mwopt flag. We then applied our recently developed marker ranking approach[4,10] to select those marker gene families that have the strongest signal of vertical evolution: that is, we assessed verticality by the extent to which class-level monophyly (GTDB release 220) was conserved.[11] This metric of marker gene quality is motivated by the hypothesis that marker genes that better resolve shallow relationships may also provide more accurate inferences of deeper, and typically more controversial, relationships. For instance, previous work indicated that marker genes which recover the reciprocal monophyly of Archaea and Bacteria - a relatively uncontroversial deep split - also tend to recover a higher proportion of more recent within-domain relationships than do genes that show mixing of archaeal and bacterial clades.[10] We then divided these marker sets into 2 subsets based on their ranking score — the top 50% ranked markers represent those gene families that have the strongest signal for vertical evolution, while the bottom 50% ranked markers have the lowest signal for verticality. Respective species trees were inferred under the LG+C60+G model using IQ-TREE v2.1.2 with mixture weights optimised for these two subsets.

Our results indicate that 34 of the NM126 markers are affected by inter-domain HGT with Bacteria. In line with this, the concatenated phylogeny inferred from the 34 genes possessed some heterodox features, such as the paraphyly of Aenigmatarchaeota, the sisterhood of Methanobacteriota_B and Undinarchaeota (Fig. S3A), as well as the placement of Altiarchaeota at the base of the DPANN archaea. By contrast, a concatenated phylogeny inferred from the remaining 89 genes recovered Altiarchaeota within Cluster 1 DPANN, in agreement with our phylogenetic analyses (Fig. 1A) and previous studies[3–5], albeit with only moderate (81%) ultrafast bootstrap and (96) SH-aLRT support (Fig. S3B). Furthermore, the phylogeny inferred from the 50% top-ranked marker genes (0.24 phylogenetic difficulty score[12]) recovered Altiarchaeota within DPANN Cluster 1 with maximal ultrafast bootstrap and SH-aLRT support (100/100) (Fig. S3C), while the phylogeny inferred from the 50% bottom-ranked markers recovered Altiarchaeota at the base of DPANN (Fig. S3D). This latter tree also possessed some other unexpected features, such as the paraphyly of Asgard archaea and Aenigmarchaeota, and the polyphyly of Methabacteriota_B (Fig. S3D). Overall, these analyses confirm that vertically evolving marker genes support a placement of Altiarchaeota as a member of DPANN Cluster 1 archaea, rather than as a sister clade of the rest of DPANN.

### Exploration of the reconciliation model and gene family filtration

To explore the effect of the various marker gene families on the rooted phylogenies, we ranked each gene family by two metrics: the number of species (the number of species with at least one gene in the gene family) and the taxonomic bias score (representation of DPANN, Euryarchaeota and TACK+Asgard lineages in the gene family; that is defined as taxonomic bias score =|⅓-%(DPANN species in the gene family)|+|⅓-%(TACK+Asgard species in the gene family)|+|⅓-%(Euryarchaeota species in the gene family)|). The motivation for the taxonomic bias score is to assess whether taxon sampling is balanced in the gene family. The smaller the score, the more representative it is, and the larger the score, the more biased the representation becomes. A set of gene families with widespread species and balanced taxon sampling is expected to contain more information to resolve the root of the archaea and be less prone to the potential 'small genome attraction artefact' in this kind of analysis[13]. That is, the model interprets the absence of a gene family in a lineage as evidence that they never had this gene, while underestimating the possibility of secondary loss. Based on these rankings, we progressively filtered out the highest-ranked gene families and re-evaluated the likelihood difference between the maximum-likelihood root (i.e., ΔLL) and competing root hypotheses under different reconciliation models (Fig. S5 A-I). For example, Fig. S5B.a illustrates the effect of progressively filtering out families with the largest number of species (Fig. S5B.a, upper) on the support of distinct roots under the global model (one D, T, and L for all branches and families). The bottom panel of each plot (e.g., Fig. S5B.a) shows the threshold values corresponding to the percentile of families removed for likelihood comparison. For example, removing the top 10% most widespread gene families corresponds to removing gene families with more than 111 species. The top panel of each plot (e.g. Fig. S5B.a) shows the change in the difference of likelihood between a competing root and the maximum-likelihood root when the same set of gene families was removed for likelihood comparison. Through this gene family filtration analysis, we can link the support for different roots (i.e., whether they received better support; ΔLL>0) with the gene families' properties and assess potential biases.

We found that when filtering out the least widespread and taxonomically biased families, the support of DPANN Cluster 2 root (i.e. the root on the DPANN clades with the smallest genomes) shifted towards alternative rooting candidates, including TACK+Asgard root (a root between TACK+Asgard and all other archaea), Euryarchaeota root (a root between Euryarchaeota and all other archaea) and DPANN root (a root between DPANN archaea and all other archaea) (Fig. S5B.a-d). This observation suggests a possible 'small-genome attraction' artefact, prompting us to explore a series of models that more accurately capture the loss, transfer, and duplication rates of DPANN based on model fit, and to investigate support for root candidates using gene family filtration. Our analyses indicated that accounting for a different loss rate for DPANN genomes helps to mitigate the small genome attraction artefact (Fig. S5C-D). However, we also observed that when a different transfer rate was assigned to the DPANN clade, the DPANN Cluster 2 root received support again (Fig. S5E-F). Another potential bias is that DPANN archaea, which are hypothesised to primarily comprise symbionts, may have a higher-than-average gene exchange rates[4]. If this is not accurately modelled, it could be another source of bias. Hence, to disentangle this, we next used a model that accounted for differential duplication, transfer, and loss (DTL) rates for DPANN, TACK+Asgard, and Euryarchaeota lineages, respectively, as well as for the largest sublineages within DPANN (i.e., Woesearchaeales+SCGC-AAA011-G17, Pacearchaeales, and Micrarchaeota). This model provides a better fit compared to previous models (Table S9). Various roots, including two roots within Euryarchaeota (MHH root, between a clade comprising Methnobacteriota_A(B), Hydrothermarchaeota and Hadarchaeota, and all other archaea; Halobacteriota+Thermoplasmatota root, between a clade comprising Halobacteriota +Thermoplasmatota and all other archaea), and DPANN Cluster 2 root received support. However, the difference in support for these different roots was not large (Fig. S5F.a-d). In addition, we modelled different origination rates for the DPANN, TACK+Asgard and Euryarchaeota. Under this model, we obtained the highest support for a root between "Euryarchaeota" and all other archaea from a set of widespread and taxonomically balanced families that are robust to filtration of up to 20% of the most widespread gene families (Fig. S5G.a-d). Because previous analyses have suggested that some other archaeal lineages may have a higher-than-average DTL rate too (e.g. Asgard archaea were suggested to have a higher duplication rate in previous studies[13,14]), we tested a final, even more complex model using differential DTL rates for the various lineages in our dataset (see Methods). Indeed, this model provided the best fit to our data among all models we tested (Table S9): it could reject all but two roots of the archaeal tree, namely a Euryarchaeota root and a root between Methanobacteriota_A(B), Hydrothermarchaeota, and Hadarchaeota, as well as all other archaea. Gene family filtration analyses under this model suggested that the support for the Euryarchaeota roots (a root between Euryarchaeota and all other archaea, and a root between a clade comprising Methanobacteriota_A(B), Hydrothermarchaeota and Hadarchaeota and all other archaea) derived from the most widespread and taxonomically balanced families; however, the root is sensitive to the filtration of these families, i.e. the progressive removal of balanced gene families shifted root support (Fig. S5I.a-d).

#### Re-analysis of the dataset from Williams et al., 2017

A previous study, based on a branch-homogeneous but family-heterogeneous reconciliation model using amalgamated likelihood estimation, ALE[15], placed the root of archaea between all DPANN archaea and the rest (DPANN root)[13]. The shift of support in the root support observed in the dataset assembled in the current study motivated us to re-analyse this dataset using a global, branch-wise, and branch-homogeneous yet family-heterogeneous reconciliation model (used also in ref[13]) and test additional rooting hypotheses overlapping with this study (e.g., a root between the DPANN Cluster 2 archaea and all other archaea). We observed consistent results with the current analyses: both the branch-homogeneous model (i.e., global and family-wise) support the genomically-reduced DPANN symbionts (i.e., DPANN Cluster 2 root) as the optimal root (P = 1). Conversely, when applying a model with clade-specific DTL rates (for Altiarchaeota, Micrarchaeota, Iainarchaeota, Nanohaloarchaeota, Aenigmarchaeota, Parvarchaeota, Nanoarchaeales, Woesearchaeales +SCGC-AAA011-G17, Pacearchaeales, Halobacteriota, Thermoplasmatota, Asgardarchaeota, Korarchaeota, and Thermoproteota) and a clade-specific DTLO rate for DPANN, “Euryarchaeota”, and TACK+Asgard, we obtained a better model fit (Table S13) and the results were consistent with our focal dataset, supporting a Euryarchaeota root.

### Metabolism of the Last Common Ancestor of Archaea (LACA)

Note that in the following, we discuss the proteome of LACA under the most highly supported roots, i.e. the Euryarchaeota and MHH roots, resolved in this study. We compare these results to LACA reconstructed under the TACK+Asgard and DPANN roots, even though these were rejected using our reconciliation approach with an improved model of gene evolution and are likely to be the result of phylogenetic artefacts. The posterior probability values listed in the following paragraphs report the presence probability of the discussed proteins under the Euryarchaeota root/MHH root/TACK+Asgard root/DPANN root, respectively, unless specified otherwise. Note that the values for posterior presence probabilities of arCOG families for the Euryarchaeota root and the MHH root are provided, respectively, in Tables S24 and S25. The posterior presence probabilities for TACK+Asgard and DPANN root can be found in the Zenodo repository (https://zenodo.org/records/17360806).

### DNA replication

The proposed hypothetical components of the LACA's replication machinery include the origin recognition complex for replication, the MCM helicase complex, both subunits of archaeal primase, the sliding clamp PCNA and both subunits of its loader RFC, the polymerases PolB and PolD, the Okazaki fragment processing flap endonuclease Fen1 and RNaseH II, the ATP-dependent DNA ligase, and both subunits of the topoisomerase VI (TOPO VI)[16,17]. Similar to previous inferences, our results inferred the presence of a rather complete DNA replication machinery in LACA (Fig. S10). Specifically, under the Euryarchaeota root, MHH root, TACK+Asgard root and DPANN root, we inferred that LACA harbours genes encoding, CDC6/OrC1 (arCOG00467_01, PP=0.93/1/0.99/0.93), DNA replicative helicase MCM (arCOG00439_01, PP=1/1/1/1), DNA replication factor GINS15 (arCOG00551_01, PP=0.87/1/1/1), single-stranded DNA specific exonuclease RecJ/Cdc45 (arCOG00427_01, PP=1/1/1/0.96), single-stranded DNA binding protein A RPA1 (arCOG01510_01, PP=0.93/0.99/0.95/0.96), both subunits of polymerase D (large subunit, arCOG04447_01, PP=1/1/0.87/1; small subunit arCOG04455_01, PP=1/1/1/1), sliding clamp PCNA homologs (arCOG00488_01, PP=0.95/0.98/1/0.75) and its loader RFC (large subunit, RFCL, arCOG00470_01, PP=0.83/0.79/0.95/0.9; small subunit, RFCS, arCOG00469_01, PP=1/1/0.99/0.99), the large subunit of archaeal primase (PriL, arCOG03013_01, PP=0.98/0.9/0.81/0.76), DNA primase (DnaG, arCOG04281_01, PP=1/1/1/1), the flap endonuclease Fen1 (arCOG04050_01, PP=0.94/0.96/0.64/0.79) and RNaseH II (arCOG04121_01, PP=0.96/0.98/0.98/0.94), ATP-dependent DNA ligase (arCOG01347_01, PP=0.99/1/0.97/0.98), DNA topoisomerase VI subunit A (arCOG04143_01, PP=0.76/0.88/1/0.94) and DNA topoisomerase VI subunit B (arCOG01165_01, PP=0.71/0.74/0.96/0.42) (Fig. S10).

Our large-scale ancestral reconstruction is necessarily incomplete due to phylogenetic uncertainty, stochastic error and mosaic ancestry of genes. For example, while we mapped the large subunit of the archaeal primase back to LACA with high confidence across all of these four roots, the support for the small subunit of archaeal primase at LACA varies across different roots (arCOG04110_01, PP=0.29/0.69/0.12/0.04). Similarly, while DNA polymerase D was suggested to be a derived form of DNA polymerase[16,18], and various studies inferred the presence of DNA polymerase B at LACA[16–18], only one DNA polymerase B arCOG family (arCOG00329_01) was inferred with sufficient PP support for the presence at LACA at the MHH root (PP=0.75).

Nevertheless, our reconstruction under three different roots, Euryarchaeota root, the MHH root and TACK+Asgard root, consistently indicated that reverse gyrase was likely present in LACA with moderate to high posterior support (respectively, arCOG01526_01, PP=0.68/0.75/0.83/0.46), in line with the optimal growth temperature (median 78.78 ℃, 95% CI: 74.15℃-83.52℃ under the Euryarchaeota root; median 79.40 ℃, 95% CI 74.18 ℃-84.30 ℃) inferred via ancestral sequence reconstruction.

### Transcription

Our ancestral gene content reconstructions indicate that LACA possessed a complete transcriptional machinery as described below:

Archaea utilise a single type of RNA polymerase (RNAP) to transcribe their gene repertoire, with which transcription factors interact to modulate activities in transcription cycles by facilitating initiation, elongation, and termination[19–21]. In summary, we identified the presence of a relatively complete RNAP at LACA under four different roots (Euryarchaeota root/MHH root/TACK+Asgard root/DPANN root) (Fig. S12). Specifically, our reconstructions inferred the presence of the two largest RNAP subunits in LACA with high support (RpoA arCOG04257_01, PP=0.99/1/1/1 and arCOG04256_01, PP=0.82/1/0.77/0.68; RpoB, arCOG01762_01, PP=0.98/1/1/0.98) (Fig. S12). In agreement with this, we also inferred the presence of all genes necessary for the efficient assembly and stability of archaeal RNAP at LACA[22] (RpoD, arCOG04241_01, PP=0.98/0.88/1/1; RpoL/RPB11, arCOG04111_01, PP=0.69/0.87/0.97/0.85; Rpo/RPB10, arCOG04244_01, PP=1/1/0.99/1), except Rpo12, which only received moderate support under the MHH root (arCOG04341_01, PP=0.6) (Fig. S12, Table S25). Our reconstruction also suggested that the stalk complex in archaeal RNAPs is likely present in LACA ( Rpo7, arCOG00675_01, PP=1/0.99/1/1; Rpo4, arCOG01016_01, PP=0.3/0.59/0.89/0.5), in agreement with previous inference[20].

In vitro characterisation of the archaeal transcription mechanism demonstrates that the initiation proceeds by step-wise assembly of the pre-initiation complex (PIC)[23] (Fig. S12). The PIC forms by TBP and TFB binding to the promoter (TATA/BRE), followed by the recruitment of RNAP, with TFE enhancing RNAP recruitment and open-complex formation[24,25]. Our reconstructions suggested that these initiation factors were present in LACA under different roots (Euryarchaeota root/MHH root/TACK+Asgard root/DPANN root): i.e., the TATA-box binding protein (TBP/SPT15, arCOG01764_01, PP=0.92/0.76/1/0.97), transcription initiation factor II B (TFB/SUA7, arCOG01981_01, PP=0.89/0.93/0.82/1) and transcription initiation factor II E (TFE/TFA1, arCOG04270_01, PP=0.89/0.97/0.91/0.92). During elongation, pauses arise due to sequence context and the presence of DNA-bound proteins, which are mitigated by elongation factors[26–28]. We were able to trace a core set of these elongation factors back to LACA, including NusA (arCOG01760_01, PP=1/0.99/0.98/0.96; arCOG01761_01, PP=0.98/0/99/0.97/0.96), the processivity factor Spt4 (arCOG04077_01, PP=0.97/1/0.93/0.74), and antiterminator NusG (arCOG01920_01, PP=1/0.97/0.15/1).

### Translation

Our ancestral gene content reconstructions suggest a LACA with a translation machinery comparable to that of modern archaeal taxa. See below:

***Ribosomal proteins****:* Our ancestral gene content reconstructions traced the presence of an extensive ribosomal repertoire in LACA with at least moderate PP support (18/23/21/17 out of 28 30S ribosome subunits, and 27/24/18/22 out of 39 50S ribosome subunits) (Fig. S11). Our results suggest that the complexity of the LACA ribosome is comparable to that of modern organisms[29–31]. Interestingly, the number of ribosomal proteins inferred to be present in the last common ancestor of DPANN archaea (LDCA), the last common ancestor of DPANN Cluster 1 (LDC1CA), the last common ancestor of DPANN Cluster 2 (LDC2CA), and the last common ancestor of Nanoarchaeota (LNCA) is almost identical (<3 differences), suggesting ribosomal proteins were not affected by genome streamlining processes of DPANN archaea.

***Translation factors****:* We traced the presence of subunits of various known archaeal initiation factors that participate in the start codon selection in the 30S ribosome units and promote the joining of the 50S ribosome subunits[32], back to LACA (Euryarchaeota root/MHH root/TACK+Asgard root/DPANN root) with PP≥0.5 (aIF1, arCOG04223_01, PP=0.9/1/0.97/0.94; aIF2 gamma arCOG01563_01, PP=1/1/1/1; aIF2 beta arCOG01640_01, PP=0.97/1/0.92/0.96; aIF1A arCOG01179_01 PP=0.7/0.96/0.7/0.67; aIF2 alpha arCOG04107_01, PP=0.98/0.99/0.98/0.98; aIF5B, arCOG01560_01, PP=0.92/1/1/1; aIF6, arCOG04176_01, PP=1/0.99/0.98/1) (Fig. S11). In agreement with various studies that suggest the antiquity of the elongation factors[33,34], we confidently traced the presence of elongation factors to LACA, i.e., the elongation factor 1A (EF-Tu, arCOG01561_01, PP=0.97/1/1/1), elongation factor G (arCOG01559_01, PP=0.99/1/1/1), and elongation factor P (arCOG04277_01, PP=1/1/1/1). Additionally, peptide release factors were inferred to be present at LACA (arCOG01742_01, eRF1, PP=0.98/0.97/0.97/0.99; arCOG01741_01, PelA, PP=0.65/0.72/0.86/0.52) (Fig. S11).

***tRNA synthetase:*** Our reconstruction suggests that the complexity of canonical tRNA synthetases in LACA is comparable to that of modern archaeal taxa[35]. Specifically, under the Euryarchaeota root/MHH root/TACK+Asgard root/DPANN root, we found strong PP support (PP≥0.75) for 19/18/19/15 canonical tRNA synthetases and moderate support (P≥0.5) for 1/3/1/3 canonical tRNA synthetases for having been present in LACA (Fig. S13). The support for the presence of cysteinyl-tRNA synthetase at LACA was only inferred under the MHH root (arCOG00486_01, PP=0.9). Rather, it appears that under the other three roots, the charging of Cysteine to tRNA may have proceeded by two steps: O-phosphoseryl-tRNA synthetase (SepRS, arCOG00411_01, PP=0.98/0.98/0.95/0.92), first charges tRNACys with O-phosphoserine, which Sep-tRNA: which is then converted to cysteine in a rRNA-dependent manner by Cys-tRNA synthase (arCOG00091_01, PP=0.71/0.85/0.79/0.64). This inference thus suggests that tRNA-dependent cysteine biosynthesis is ancient in archaea. Additionally, this pathway is present in various Euryarchaeota and utilised by methanogens[36–39], which is consistent with the hypothesis that LACA may have been a methanogen. Finally, our analysis did not identify pyrrolysinyl-tRNA synthetase in LACA (arCOG00413_01).

### Protein quality control

Our inferences suggest that LACA likely already possessed various strategies to handle misfolded proteins. Specifically, we found strong support for the presence of prefoldin (arCOG01341_01, PP=0.98/1/0.92/0.86, arCOG01342_01, PP=0.94/0.98/0.9/0.9), archaeal chaperonin (GroEL, arCOG01257_01, PP=0.99/1/1/0.98), and cdc48/valosin-containing protein (arCOG01308_01, PP=0.75/0.9/0.99/0.95) in LACA (under the Eury/MHH roots/TACK+Asgard root/DPANN root), which might assist in the correct folding of newly synthesised/denatured proteins (Fig. S14). Additionally, our reconstruction also suggests a route of degrading misfolded proteins in LACA (Fig. S14). For example, we inferred the presence of 20S proteasome (arCOG00970_01, PP=0.97/0.97/1/0.97; arCOG00971_01,

PP=0.96/0.99/0.95/0.99) with its assembly and regulatory factors (arCOG01306_01, PP=0.95/0.94/0.85/0.77; arCOG00348_01, PP=1/1/0.98/0.99), and various proteases including Zn-dependent protease with chaperone function HptX (arCOG01331_01, PP=0.9/0.95/0.86/0.91), membrane protease subunit HflC, arCOG01915_01, PP=1/0.98/0.97/1) and ATP-dependent protease LonB (arCOG02160_01, PP=1/0.98/0.97/0.98).

### Central carbon metabolism

Our reconstruction indicated that LACA likely did not encode key enzymes of the Entner-Doudoroff (ED) pathway, such as the gluconate dehydratase (Gad, arCOG01168_01, PP=0.02/0.03/0/0.01), or that of the upper glycolytic pathway, e.g., glucose-6-phosphate isomerase (pgi, arCOG00052_01, PP=0/0.03/0.03/0). This suggests LACA was likely unable to utilise complex carbohydrates as a carbon or energy source. However, LACA likely contained candidate genes encoding enzymes for most steps of the lower-glycolytic pathway, catalysing reactions from glyceraldehyde-3-phosphate to pyruvate (Fig. S15). For example, we found strong support that genes encoding enzymes such as glyceraldehyde-3-phosphate dehydrogenase (GapA, arCOG00493_01, PP=0.93/0.99/0.87/0.95), which converts glyceraldehyde-3-phosphate into 1,3-bisphosphoglycerate, generating NADH, and 3-phosphoglycerate kinase (Pgk, arCOG00496_01, PP=0.98/1/0.99/0.99), which then converts that into 3-phosphoglycerate and produces ATP, were present in LACA. We also found that LACA had the ability to convert pyruvate into acetyl-CoA using a pyruvate: ferredoxin oxidoreductase (PorA, arCOG01608_01, PP=1/0.93/0.98/0.93; PorD, arCOG01604_01, PP=0.98/0.98/1/0.96; PorG, arCOG01602_01, PP=0.93/0.93/0.53/0.29) while generating reduced ferredoxin. Acetyl-CoA could have been metabolised to acetate via the ADP-forming acetyl-CoA synthetase in LACA (AcdA: arCOG01340_01, PP=0.66/0.85/0.96/0.83; AcdB, arCOG01338_01, PP=0.97/0.99/0.99/0.98), allowing the production of ATP.

LACA (under the Euryroot/MHHroot/TACK+Asgard root/DPANN root) was not predicted to encode the key genes for enzymes mediating gluconeogenesis, e.g., the bifunctional fructose-1,6-bisphosphate aldolase/phosphatase FBPA/FBPase (arCOG04180_01, PP=0.17/0.15/0.05/0.03) (Fig. S15), consistent with a previous reconstruction [13]. Our analyses predicted however that some components of the tricarboxylic acid cycle (TCA) were present in LACA, including a citrate synthase (arCOG02093_01, PP=0.7/0.73/0.67/0.5), the alpha subunit of pyruvate: ferredoxin oxidoreductase (KorA, arCOG01607_01, PP=0.64/0.87/0.69/0.47), and subunits of succinyl-CoA synthetase (SucC, arCOG01337_01, PP=0.98/1/0.97/0.95 and SucD, arCOG01339_01, PP=0.98/0.99/0.99/0.95). However, enzymes such as malate dehydrogenase (Mdh, arCOG00246_01) and succinate dehydrogenase (SdhC, arCOG02244_01; SdhD, arCOG04162_01) likely were not encoded by LACA.

We also inferred the presence of a rather complete non-oxidative pentose phosphate pathway (PPP) at LACA, including transketolase (arCOG01051_01, PP=0.6/0.94/0.21/0.42), ribose-5-phosphate isomerase (arCOG01122_01, RpiA, PP=0.97/0.99/0.94/0.99), and pentose-5-phosphate-3-epimerase (arCOG05046_01, PP=0.85/0.99/0.65/0.7). Additionally, we inferred the presence of formaldehyde-activating enzyme (Fae)/hexulose-6-phosphate synthase (Hps) (arCOG00103_01,

PP=0.79/0.89/0.6/0.53) and D-arabinose,5-phosphate isomerase (GutQ/HxlB, arCOG00068_01, PP=0.94/1/0.99/0.47) at LACA, which could have allowed the biosynthesis of ribulose-5-phosphate to be further converted to ribose-5-phosphate via RpiA, providing substrates for purine biosynthesis (Fig. S15).

### Wood-Ljungdahl pathway

The Wood-Ljungdahl pathway (WLP) consists of methyl and carbonyl branches, which respectively catalyse the reversible reduction of $CO_2$ into methyl and carbonyl moieties of acetyl-coenzyme A (Acetyl-CoA)[40,41]. Two variants of methyl branches exist: one uses tetrahydrofolate (H4F) while the other uses methanofuran (MF) and tetrahydromethanopterin (H4MPT) as cofactor carriers of reduced carbon. Our reconstructions indicate that all subunits of the H4F methyl branch were likely absent in LACA (Fig. S16, PP < 0.5), consistent with the observation that the H4F methyl branch is prevalent in bacteria but not archaea[42] and a previous analysis[13]. However, we inferred that the H4MPT branch was likely present in LACA, consistent with previous studies[40,43,44]. Ancestral reconstructions based on arCOG families showed moderate to high support for formylmethanofuran dehydrogenase subunit A (arCOG04461_01, FwdA, PP=0.79/0.94/0.76/0.6), subunit B (arCOG01499, FwdB, PP=0.62/0.86/0.45/0.31), subunit C (arCOG00097_01, FwdC, PP=0.91/0.97/0.95/0.92), subunit D (arCOG02674_01, FwdD, PP=0.98/1/0.95/0.8), formylmethanofuran: tetrahydromethanopterin formyltransferase (arCOG02695_01, Ftr, PP=0.92/0.95/0.75/0.7) and methenyltetrahydromethanopterin cyclohydrolase (arCOG02675_01, Mch, PP=0.89/0.83/0.83/0.66) (Fig. S16).

The carbonyl branch of the WLP is linked with the methyl branch by the key enzyme CO dehydrogenase/acetyl-CoA synthase (ACS/CODH), which synthesises acetyl-CoA from carbon monoxide and methyl-H4MPT via the methylated corrinoid Fe-S protein CH3COFeSP[45]. This enzyme complex generally consists of five subunits and often co-occurs in the genome with a complete H4MPT methyl branch[40]. Our ancestral reconstructions show high PP support for four subunits (PP≥0.75), including CdhB (arCOG04408_01, PP=0.92/0.93/0.89/0.81), CdhC (arCOG04360_01, PP=0.88/1/0.92/0.77), CdhD (arCOG01980_01, PP=0.86/0.98/0.98/0.93), and CdhE (arCOG01979_01, PP=0.74/0.88/0.75/0.53) (Fig. S16). The high support for these four subunits and the presence of a complete H4MPT methyl branch suggest that a complete ACS/CODH enzyme was likely present in LACA, in line with a previous study[45]. Our reconciliation also inferred that LACA may have encoded an acyl-CoA synthetase (arCOG01529_01; PP=0.94/0.97/0.92/0.86) and the ADP-forming acetyl-CoA synthetase that can activate acetate into acetyl-CoA.

In summary, our reconstruction suggests that LACA might have encoded the H4MPT methyl-branch and carbonyl-branch of the WLP, in agreement with various inferences that suggest the antiquity of this pathway[43–46].

### Biosynthesis of cofactors related to WLP

We found support for many genes encoding enzymes involved in the biosynthesis of various cofactors present in LACA, in agreement with the inference of a complete WLP. LACA under the Euryarchaeota root and MHH root likely encoded almost all genes for the biosynthesis of Coenzyme F430 (cfbA, arCOG02246_01, PP=0.49/0.96/0.41/0.25; cfbB, arCOG00106_01, PP=0.76/0.99/0.54/0.47; cfbD, arCOG04888_01, PP=0.87/0.96/0.75/0.53), Coenzyme F420 (CofC, arCOG04472_01, PP=0.96/0.97/0.86/0.76; CofD, arCOG04395_01, PP=0.48/0.62/0.79/0.3; CofE, arCOG02714_01, PP=1/0.98/0.97/0.92), Coenzyme B (AksD, arCOG01698_01, PP=0.97/1/0.99/0.92; AksE, arCOG02230_01, PP=0.95/0.98/0.86/0.75; AksF, arCOG01163_01, PP=0.03/0.25/0.03/0.01) and Coenzyme M (ComA, arCOG04896_01, PP=0.68/0.93/0.45/0.23; ComB, arCOG04871_01, PP=0.78/0.94/0.71/0.55; ComC, arCOG04874_01, PP=0.94/1/0.98/0.88, ComE/D, arCOG01614_01, PP=0.67/0.97/0.49/0.41), given that more than half of those genes in the biosynthesis pathway were found to have PP ≥ 0.5.

In methanogenic archaea, formylmethanofuran dehydrogenase has a molybdopterin dinucleotide cofactor and catalyses the first step in the WLP[47]. Our reconstructions also indicate that LACA likely encoded all genes of the molybdopterin biosynthesis, consistent with the inference of a complete formylmethanofuran dehydrogenase complex. These subunits include GTP-3',8-cyclase (arCOG00930_01, MoaA, PP=0.86/0.99/0.92/0.85), cyclic pyranopterin monophosphate synthase (arCOG01530_01, MoaC, PP=0.85/0.96/0.74/0.51), Molybdopterin converting factor (large subunit, arCOG00534_01, MoaE, PP=0.95/0.95/0.83/0.85; small subunit, arCOG00536_01, MoaD, PP=0.89/0.97/0.82/0.48), molybdopterin adenylyltransferase (arCOG00214_01, PP=0.71/0.95/0.64/0.47) and molybdopterin molybdotransferase (arCOG00216_01, PP=0.98/1/0.97/0.91).

### Methanogenesis

Our reconstruction provided support for methyl-coenzyme M reductase under the Euryarchaeota root and the MHH root. Furthermore, under the MHH root, LACA was suggested to have encoded almost all subunits of a membrane-bound, sodium-translocating methyltransferase complex: the $N^5$-tetrahydromethanopterin: CoM-S-methyltransferase (Mtr complex) (Fig. S16). A methylcobalamin: coenzyme M methyltransferase (MtbA, arCOG03325_01, PP=0.96/0.94/0.97/0.85) was inferred under all four different rooting scenarios (i.e., Euryarchaeota root/MHH root/TACK+Asgard root/DPANN root). Taken together, our results provide evidence for a methanogenic LACA under the most likely roots, i.e. the Euryarchaeota and MHH root. However, given that most of the Mtr complex was only inferred under the MHH root, our results suggest that LACA would have had the ability to perform methylotrophic methanogenesis under the Euryarchaeota root[46] and both $CO_2$-reducing and methyltrophic methanogenesis under the MHH root[43,46]. These two substrates have been suggested to have been abundant in early Earth environments[48,49].

In accordance with this, various methanogenesis marker proteins that were previously defined as being specific to class I/II methanogens[50] were also traced back to LACA with strong support, e.g., methanogenesis marker 3 (arCOG04900_01, PP=0.82/0.96/0.06/0.53), methanogenesis marker 9 (arCOG04853_01, PP=0.59/0.78/0.57/0.31), and methanogenesis marker 2 (Selenophosphate synthetase related protein, arCOG00640_01, PP=0.96/0.99/0.96/0.89).

### [NiFe] hydrogenases

Given the challenge to accurately assign sequences to arCOGs for the large and small subunits of the [NiFe]-hydrogenase, we instead retrieved homologues for these subunits using the their key domain, i.e. IPR006137 for the small subunit and IPR029014 for the large subunit and performed additional phylogenetic analysis to rule out false homologs, in order to classify them into different groups of [NiFe]-hydrogenases[51,52]. Our phylogenetic analysis only recovered a small number of archaeal sequences affiliated with group 1 [NiFe] hydrogenase (29 sequences) and group 2 [NiFe]-hydrogenase (9 sequences), consistent with the previous observation that group 1 and group 2 [NiFe]-hydrogenases were primarily found in Bacteria[53] (Fig. S17). Therefore, we chose to focus on group 3 [NiFe]-hydrogenase and group 4 [NiFe]-hydrogenase for reconciliations.

Our reconciliation predicted the presence of a group 3 [NiFe]-hydrogenase and group 4 [NiFe]-hydrogenase in LACA with strong PP support (group 3 [NiFe]-hydrogenase, PP=0.98/0.99/0.92/0.43, group 4 [NiFe]-hydrogenase, PP=0.9/0.99/0.9/0.76). We further divided Group 3 [NiFe]-hydrogenase into three largest subgroups: Group 3a, Group 3b, and Group 3c, whose marked diversity might stem from an ancient origin and a long evolutionary history in Archaea. Interestingly, we also found support for the presence of group 3b [NiFe]-hydrogenase (PP=0.84/0.89/0.8/0.54) and group 3c [NiFe]-hydrogenase in LACA (PP=0.66/0.68/0.72/0.64). Group 3c [NiFe]-hydrogenase is functionally linked to heterodisulfide reductase and is widespread in methanogens[54]. Since the group 4 [NiFe]-hydrogenase has many small subgroups that are poorly defined so far, we did not further subdivide those. Note that posterior presence probabilities for the Euryarchaeota root and MHH root can be found in Tables S26-S27, while that for the TACK+Asgard root and DPANN root can be accessed from the Zenodo repository (https://zenodo.org/records/17360806).

### Biosynthetic potential of the LDCA, LDC1CA, LDC2CA and LNCA.

Because extant DPANN archaea are known to have lost various biosynthetic pathways, we investigated the evolution of biosynthesis genes throughout DPANN diversification.

***Nucleotide biosynthesis***: The *de novo* biosynthesis of the purine pathway has 17 genes (Fig. S15). Our analyses traced 12-14 purine biosynthesis gene families back to the Last DPANN Common Ancestor (LDCA) and the Last DPANN Cluster 1 Common Ancestor (LDC1CA) under the two most likely roots, i.e., the Euryarchaeota and MHH root with PP≥0.5. However, only 1-3 of them were traced back to the Last

DPANN Cluster 2 Common Ancestor (LDC2CA) and the Last Nanoarchaeota Common Ancestor (LNCA). This reduction is similar but less apparent in the *de novo* biosynthesis of the pyrimidine pathway. Among 14 genes investigated, 13-14 of them were still traced back to the LDCA and LDC1CA with PP≥0.5 under the two most likely roots. Rooting with the Euryarchaeota and MHH roots, respectively, maps 7 and 6 and 10 and 4 of these gene families back to the LDC2CA and LNCA. Hence, our analyses suggest that the loss of the de novo purine biosynthesis pathway occurred in the LDC2CA prior to the loss of the de novo pyrimidine biosynthesis pathway under both scenarios. LNCA already lacked both of these pathways. The prior loss of the de novo purine biosynthesis pathway we observed here is in broad agreement with many bacterial symbionts. For example, the loss of the purine biosynthesis pathway is more pronounced than the pyrimidine biosynthesis pathway in many insect bacterial symbionts[55,56]. This loss is potentially constrained by the energetic demand of synthesising purine *de novo*, as this pathway is longer and more ATP-intensive[57].

***Lipid biosynthesis***: In summary, we recovered the modified mevalonate pathway (MVA) in LACA[58]. Out of 8 genes of interest from the MVA pathway, we inferred the putative presence of 7-8 of these genes in LDCA and LDC1CA under both the Euryarchaeota and MHH roots, suggesting that the LDCA and LDC1CA were capable of synthesising isoprenoids. Nevertheless, this MVA pathway appeared to have been lost already in the LDC2CA under the Euryarchaeota root (only 2 genes with PP≥0.5 were mapped to this ancestor), while it was inferred to potentially have been present under the MHH root (5 genes mapped back to LDC2CA, including mevalonate kinase (arCOG01028)). This pathway was subsequently entirely lost in the LNCA, as suggested under both the Euryarchaeota root and the MHH root, i.e. none of the gene families were inferred to be present in the LNCA (Fig. S15).

We can confidently assign 7 genes to the glycerophospholipid biosynthesis pathway (from glyceraldehyde-3P to archaetidylinositol/archaetidylserine). Five of these could be traced back to LDCA and LDC1CA (with PP ≥ 0.5). While 4/5 of these could still be traced back to LDC2CA under the Euryarchaeota root and the MHH root (PP≥0.5), only CDP-diglyceride synthetase (arCOG04106_01) was inferred to be present at LNCA. Together, these data indicate a stepwise reduction in lipid biosynthesis potential throughout the evolution of DPANNs.

***Vitamin biosynthesis pathway***: Because this gene category encompasses multiple pathways, we summarise the major observations here, with details reported in Tables S24-S25. Specifically, we mapped only 16 and 18 genes associated with vitamin biosynthesis back to the LDCA and LDC1CA (PP ≥ 0.5), respectively, under the Euryarchaeota root; whereas for LDC2CA and LNCA, this number was reduced to 4. Similarly, under the MHH root, 33 and 29 genes were mapped back to the LDCA and LDC1CA (PP≥0.5), respectively; 10 and 5 genes were mapped back to the LDC2CA and LNCA. In comparison, 43 and 66 genes were traced back to LACA, respectively, under the Euryarchaeota root and the MHH root. This suggests that the early loss of the vitamin biosynthesis pathway occurred in LDCA and that these pathways were subsequently streamlined throughout DPANN evolution.

***Amino acid biosynthesis***: Similarly, multiple pathways are included under this category, and details are provided in Table S24-S25. Here, we summarise the number of genes associated with amino acid biosynthesis under the Euryarchaeota and MHH root. Specifically, 32, 34, 5, and 3 genes, respectively, were mapped back to the LDCA, LDC1CA, LDC2CA, and LNCA under the Euryarchaeota root, while under the MHH root, the corresponding numbers are 45, 47, 10, and 5 genes. In comparison, 41 and 51 genes were traced back to LACA, respectively, under the Euryarchaeota root and the MHH root. This is similar to the vitamin biosynthesis pathway.

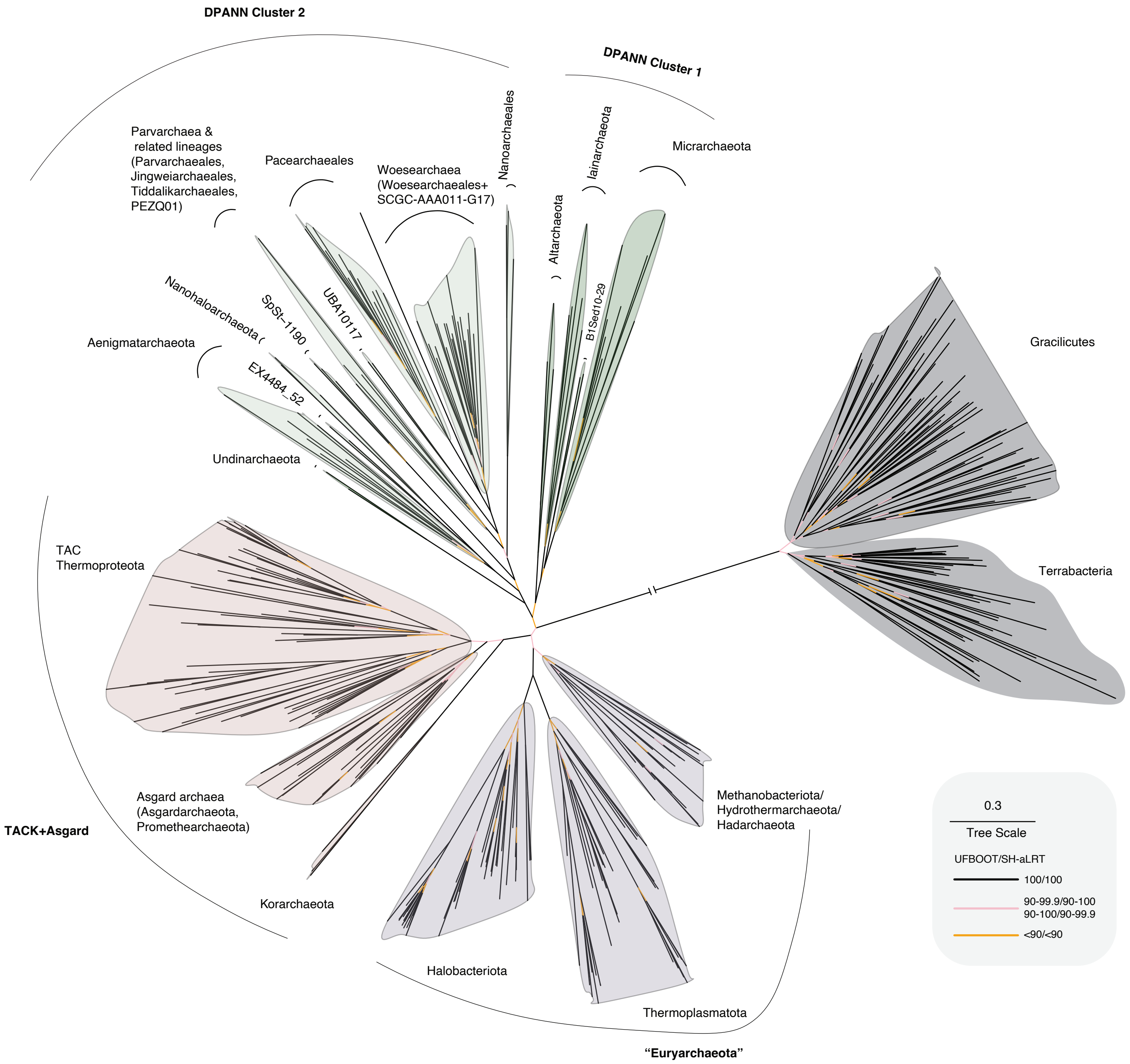


**Fig. S1: Maximum likelihood outgroup-rooted archaeal phylogeny.** The maximum likelihood phylogeny was inferred under the best-fitting LG+C60+F+R4 model on a concatenation of 25 vertically evolving marker genes shared between Bacteria and Archaea. The inferred archaeal root places DPANN as sister to the remaining Archaea (DPANN root), but this placement is weakly supported by ultrafast bootstrap and SH-aLRT metrics, and several alternative rootings cannot be rejected by an Approximately Unbiased (AU) test (Table S3). Scale bar indicates the mean number of substitutions per site.

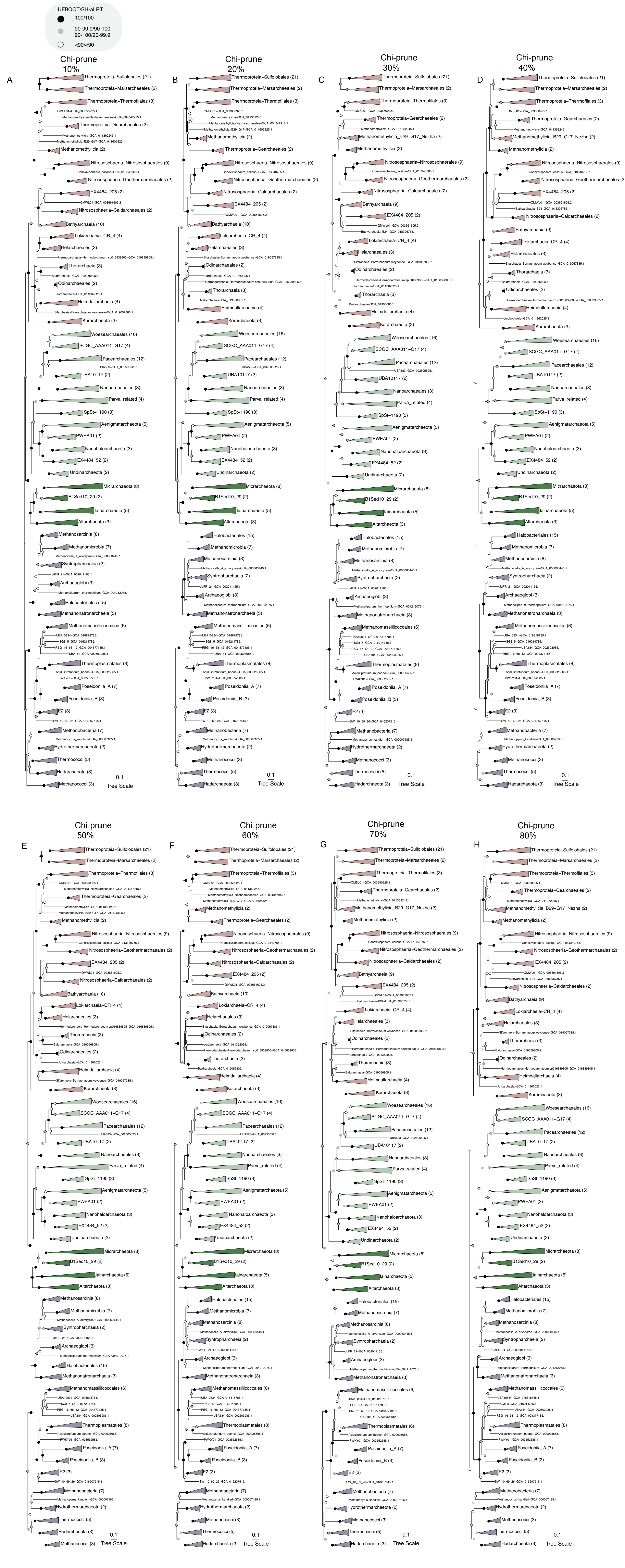


**Fig. S2: Maximum likelihood archaeal phylogeny under the best-fitting site-heterogeneous substitution model (LG+C60+F+R4) following the removal of 10%-80% most compositionally heterogeneous sites.** Note that the trees are manually rooted between Euryarchaea and all other archaea. The topologies are highly congruent with each other. and the branches representing the 15 rooting hypotheses are monophyletic when 10% to 80% of compositionally heterogeneous sites are removed. **A**. 10% of the most heterogeneous sites were removed, leaving 9348 sites out of 10386 sites. **B**. 20% of the most heterogeneous sites were removed, leaving 8309 sites out of 10386 sites. **C**. 30% of the most heterogeneous sites were removed, leaving 7271 sites out of 10386 sites. **D**. 40% of the most heterogeneous sites were removed, leaving 6232 sites out of 10386 sites. **E**. 50% of the most heterogeneous sites were removed, leaving 5193 sites out of 10386 sites. **F**. 60% of the most heterogeneous sites were removed, leaving 4155 sites out of 10386 sites. **G**. 70% of the most heterogeneous sites were removed, leaving 3116 sites out of 10386 sites. **H**. 80% of the most heterogeneous sites were removed, leaving 2078 sites out of 10386 sites. Scale bar: average number of substitutions per site.

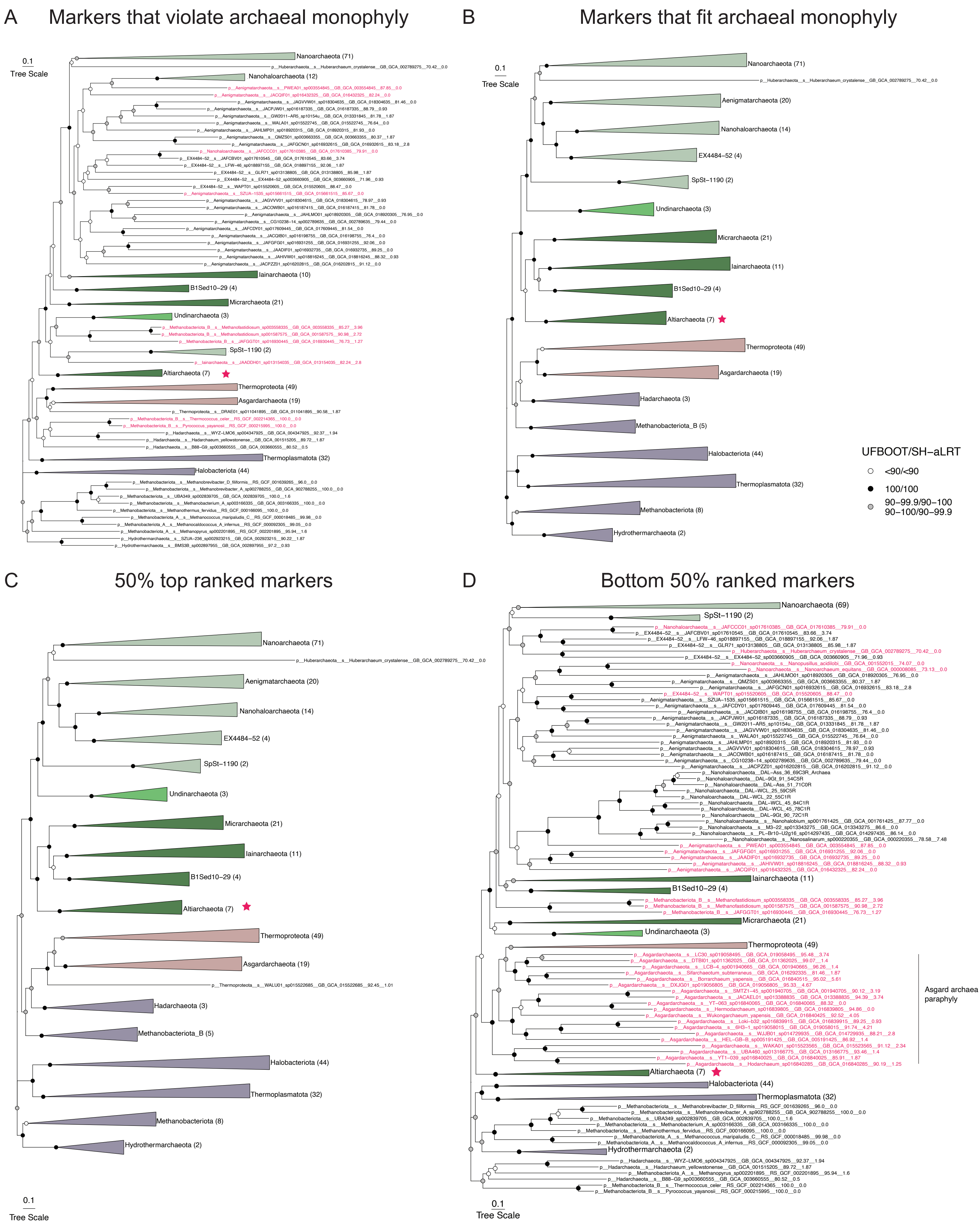


**Fig. S3: Maximum likelihood archaeal phylogenies under LG+C60+G models of different subsets of marker genes of the dataset from Baker et al., 2025.** Note that red stars indicate the position of Altiarchaeota. **A**. The phylogeny of the concatenation of 34 markers that violate archaeal monophyly. **B**. The phylogeny of the concatenation of 89 markers that conformed to archaeal monophyly. **C**. The phylogeny of the concatenation of 45 markers ranked above the 50th percentile among the 89 marker genes (See Supplementary Methods). **D**. The phylogeny of the concatenation of 45 markers ranked below the 50th percentile among the 89 marker genes (See Supplementary Methods). Scale bar: average number of substitutions per site.

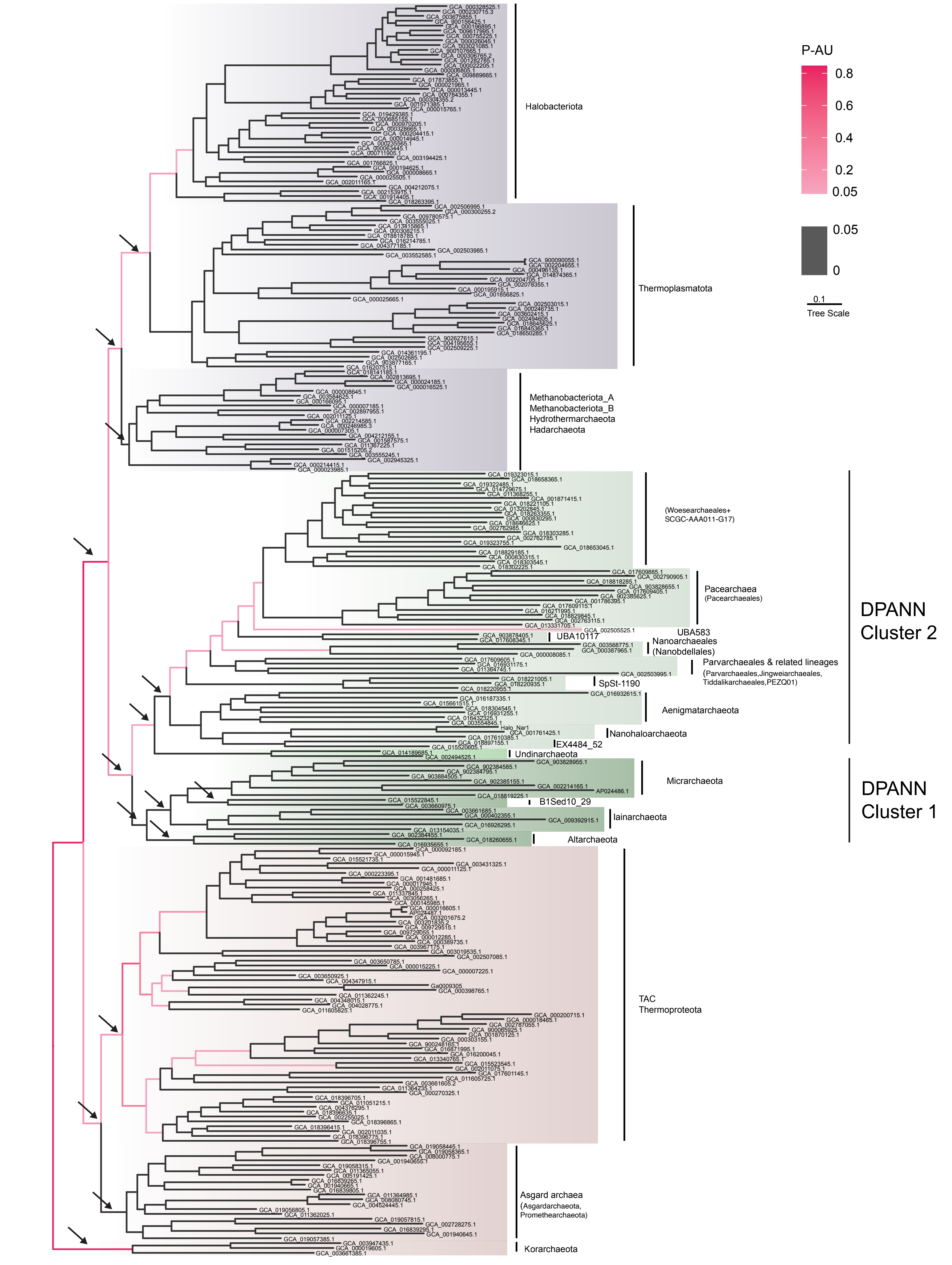


**Fig. S4: Maximum likelihood phylogeny of archaea using a non-reversible model.** The phylogeny was inferred with the NONREV+G model. This phylogeny placed the archaeal root between Korarchaeota and all other archaea (Korarchaeota root). However, an AU test of all branches in the tree revealed a wide range of statistically indistinguishable rooting positions (P > 0.05), colored in red, including the branch leading to the extremely genome-reduced lineage of Nanoarchaeales. The results of an AU test of rooting hypotheses overlapping with the main set are summarised in Table S6. Scale bar: average number of substitutions per site. Arrows indicates the roots overlapping with the main set.

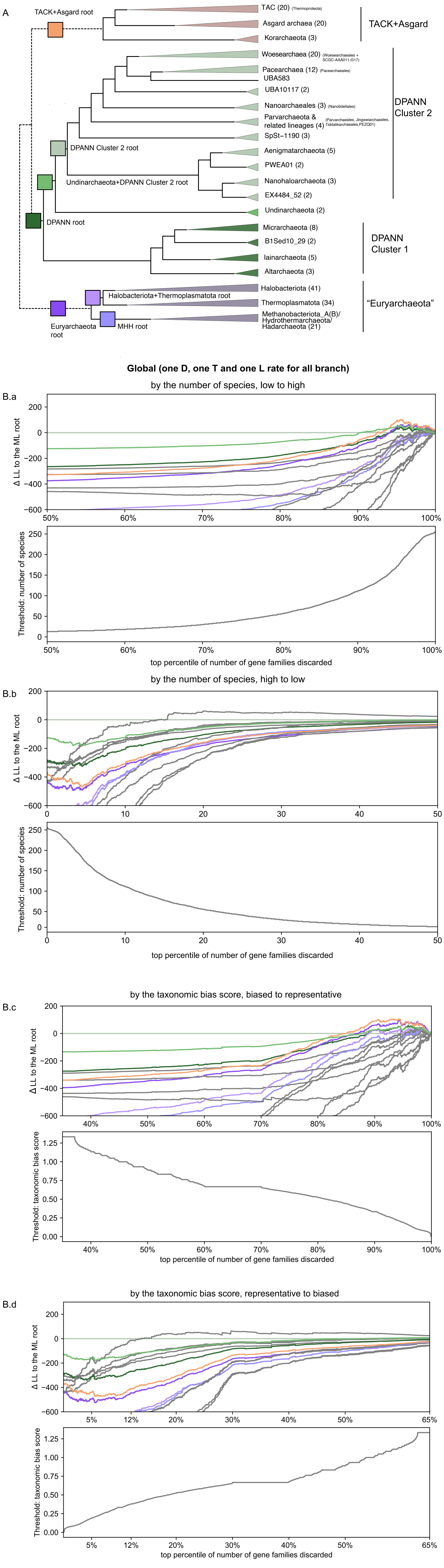
A
TACK+Asgard root
TAC (20)
Asgard archaea (20)
Korarchaeota (3)
TACK+Asgard
Woesearchaea (20)
Pacearchaea (12)
UBA583
UBA10117 (2)
Nanoarchaeales (3)
Parvarchaeota & related lineages (4)
SpSt−1190 (3)
DPANN Cluster 2
DPANN Cluster 2 root
Aenigmatarchaeota (5)
PWEA01 (2)
Undinarchaeota+DPANN Cluster 2 root
Nanohaloarchaeota (3)
EX4484_52 (2)
Undinarchaeota (2)
DPANN root
Micrarchaeota (8)
B1Sed10_29 (2)
Iainarchaeota (5)
Altarchaeota (3)
DPANN Cluster 1
Halobacteriota (41)
Halobacteriota+Thermoplasmatota root
Thermoplasmatota (34)
"Euryarchaeota"
Euryarchaeota root
MHH root
Methanobacteriota_A(B)/ Hydrothermarchaeota/ Hadarchaeota (21)
Global (one D, one T and one L rate for all branch)
by the number of species, low to high
B.a
Δ LL to the ML root
Threshold: number of species
top percentile of number of gene families discarded
by the number of species, high to low
B.b
Δ LL to the ML root
Threshold: number of species
top percentile of number of gene families discarded
by the taxonomic bias score, biased to representative
B.c
Δ LL to the ML root
Threshold: taxonomic bias score
top percentile of number of gene families discarded
by the taxonomic bias score, representative to biased
B.d
Δ LL to the ML root
Threshold: taxonomic bias score
top percentile of number of gene families discarded

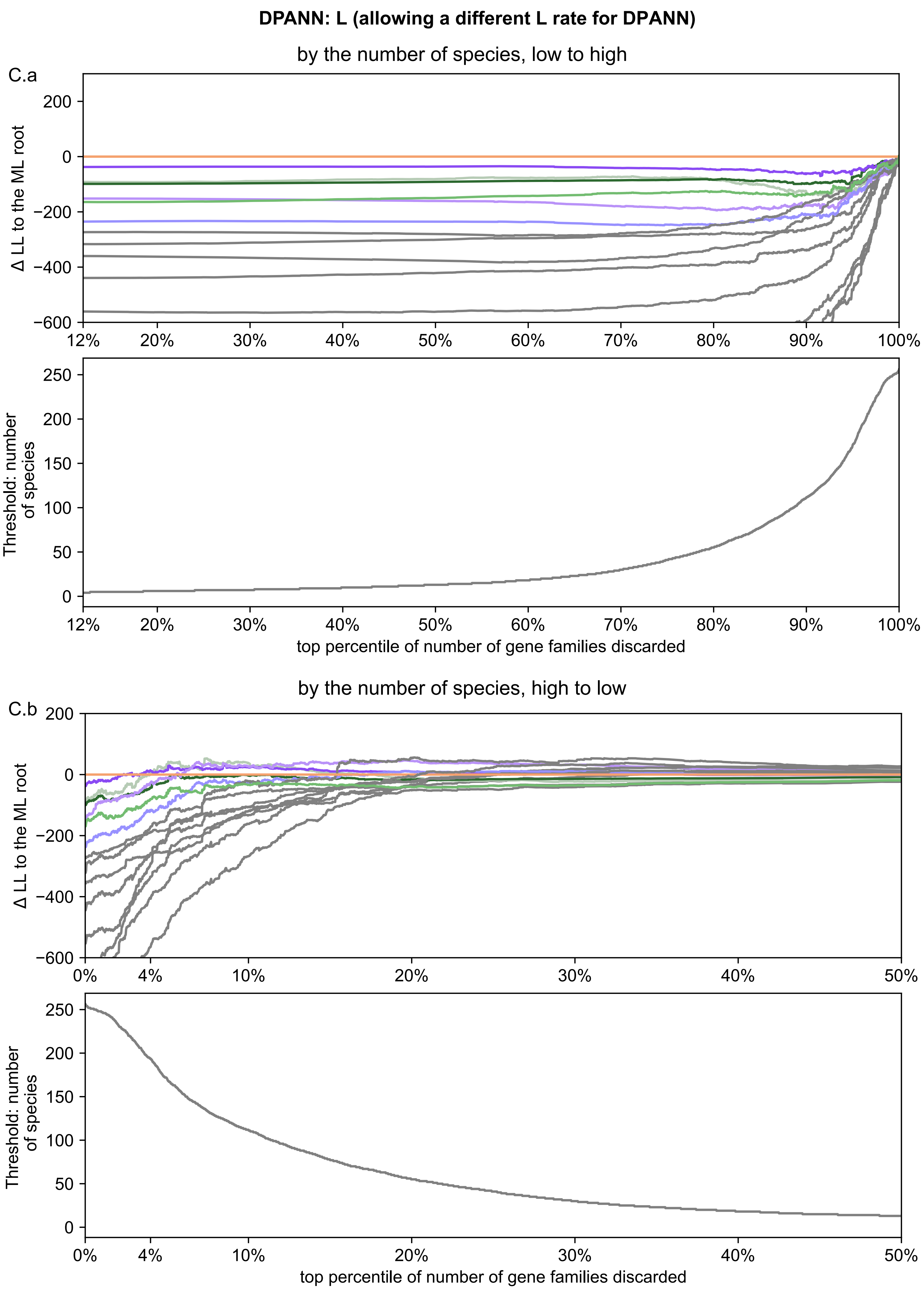
DPANN: L (allowing a different L rate for DPANN)
by the number of species, low to high
C.a
Δ LL to the ML root
Threshold: number of species
top percentile of number of gene families discarded
by the number of species, high to low
C.b
Δ LL to the ML root
Threshold: number of species
top percentile of number of gene families discarded

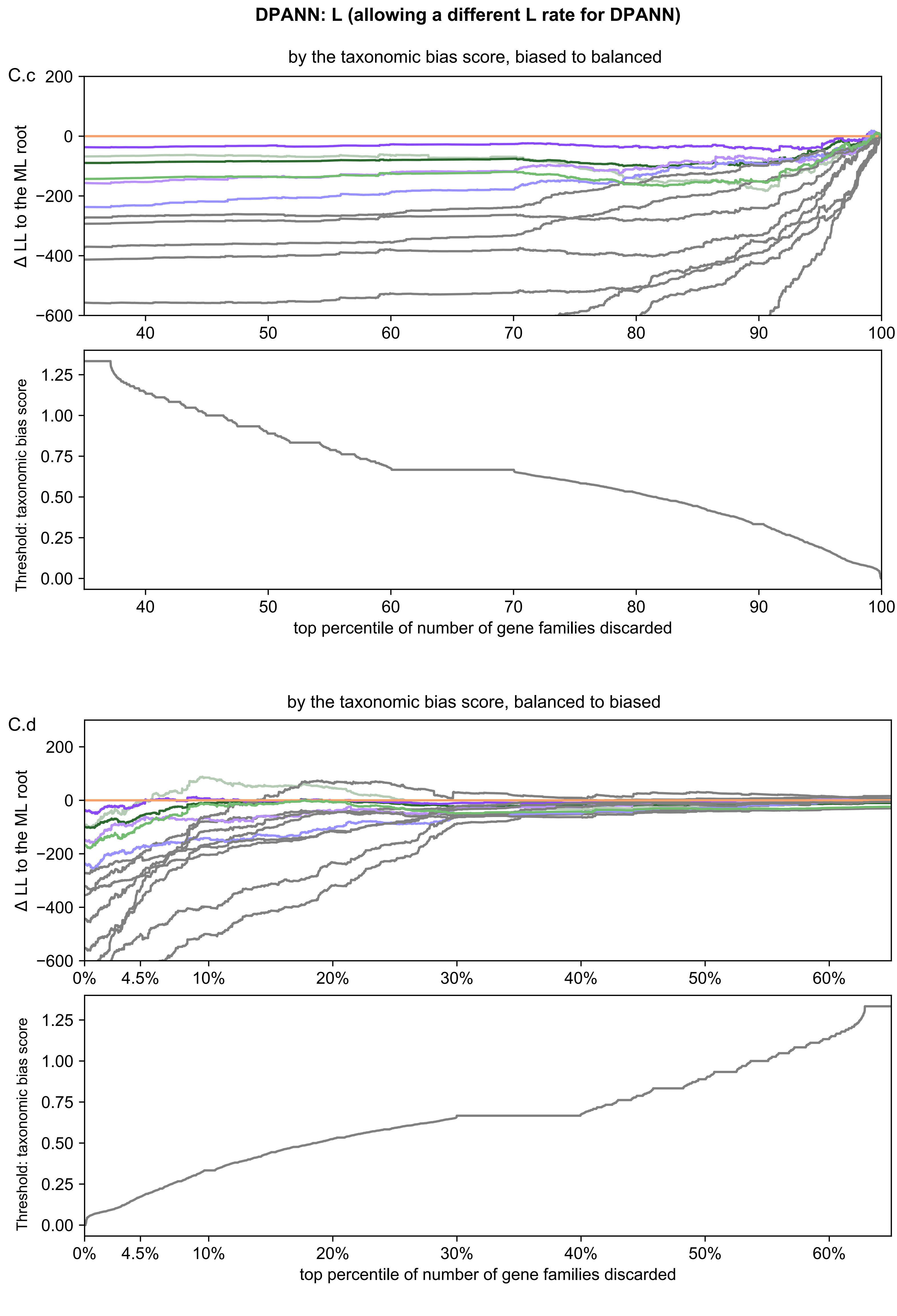
DPANN: L (allowing a different L rate for DPANN)
by the taxonomic bias score, biased to balanced
C.c
Δ LL to the ML root
Threshold: taxonomic bias score
top percentile of number of gene families discarded
by the taxonomic bias score, balanced to biased
C.d
Δ LL to the ML root
Threshold: taxonomic bias score
top percentile of number of gene families discarded

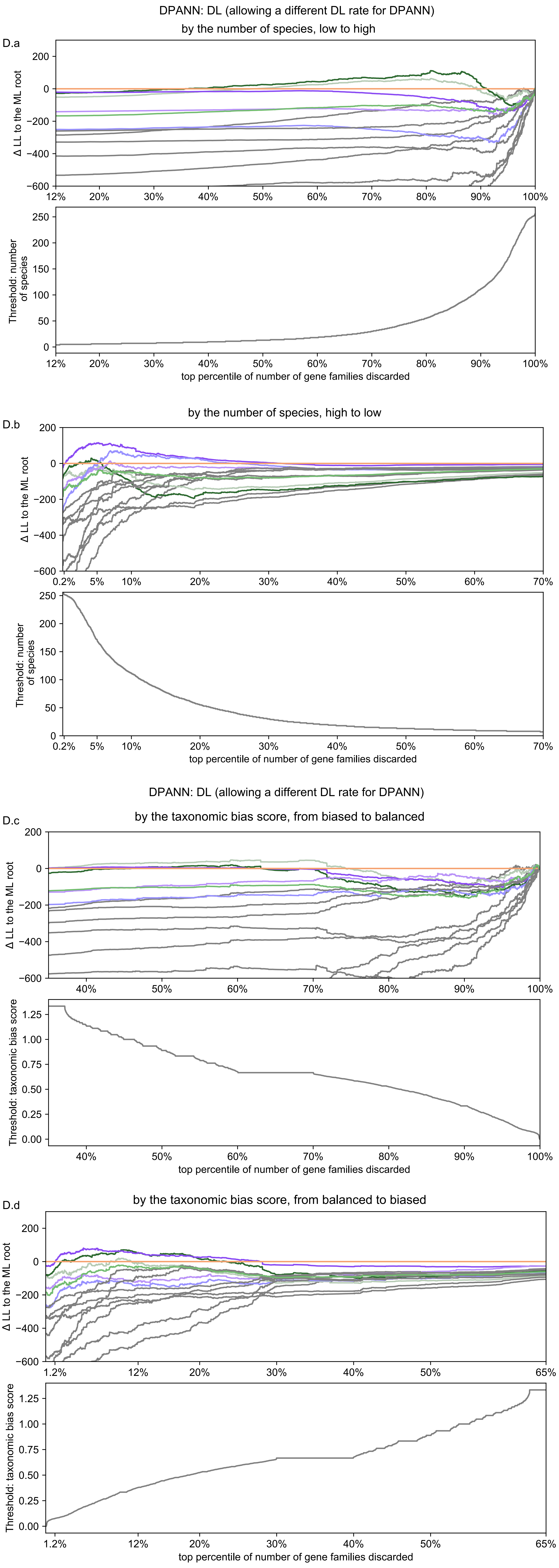
DPANN: DL (allowing a different DL rate for DPANN)
by the number of species, low to high
D.a
Δ LL to the ML root
Threshold: number of species
top percentile of number of gene families discarded
by the number of species, high to low
D.b
Δ LL to the ML root
Threshold: number of species
top percentile of number of gene families discarded
DPANN: DL (allowing a different DL rate for DPANN)
by the taxonomic bias score, from biased to balanced
D.c
Δ LL to the ML root
Threshold: taxonomic bias score
top percentile of number of gene families discarded
by the taxonomic bias score, from balanced to biased
D.d
Δ LL to the ML root
Threshold: taxonomic bias score
top percentile of number of gene families discarded

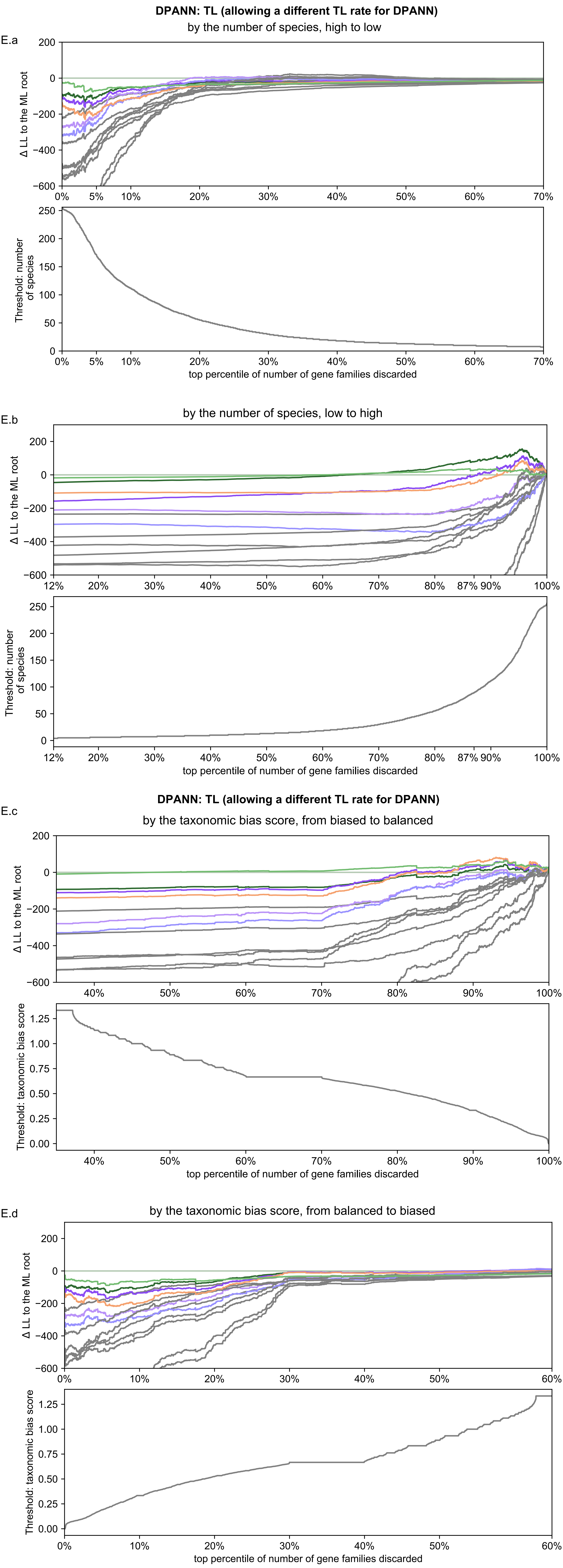

DPANN: TL (allowing a different TL rate for DPANN)
by the number of species, high to low
E.a
Δ LL to the ML root
200
0
−200
−400
−600
0%
5%
10%
20%
30%
40%
50%
60%
70%
Threshold: number of species
250
200
150
100
50
0
top percentile of number of gene families discarded
by the number of species, low to high
E.b
12%
80%
87%
90%
100%
DPANN: TL (allowing a different TL rate for DPANN)
by the taxonomic bias score, from biased to balanced
E.c
Threshold: taxonomic bias score
1.25
1.00
0.75
0.50
0.25
0.00
by the taxonomic bias score, from balanced to biased
E.d

**DPANN: DTL (allowing a different DTL rate for DPANN)**

F.a

by the number of species, low to high

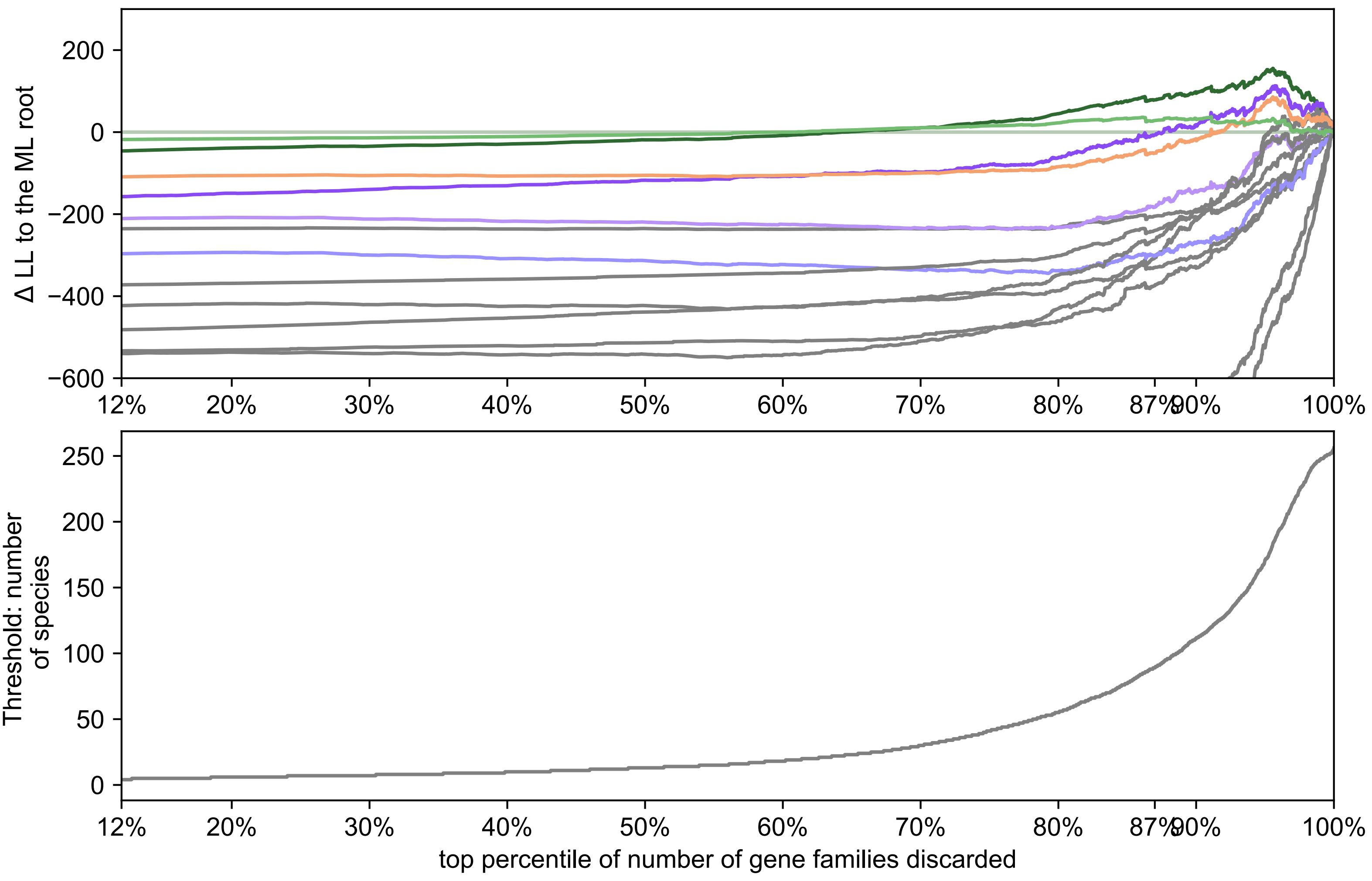


F.b

by the number of species, high to low

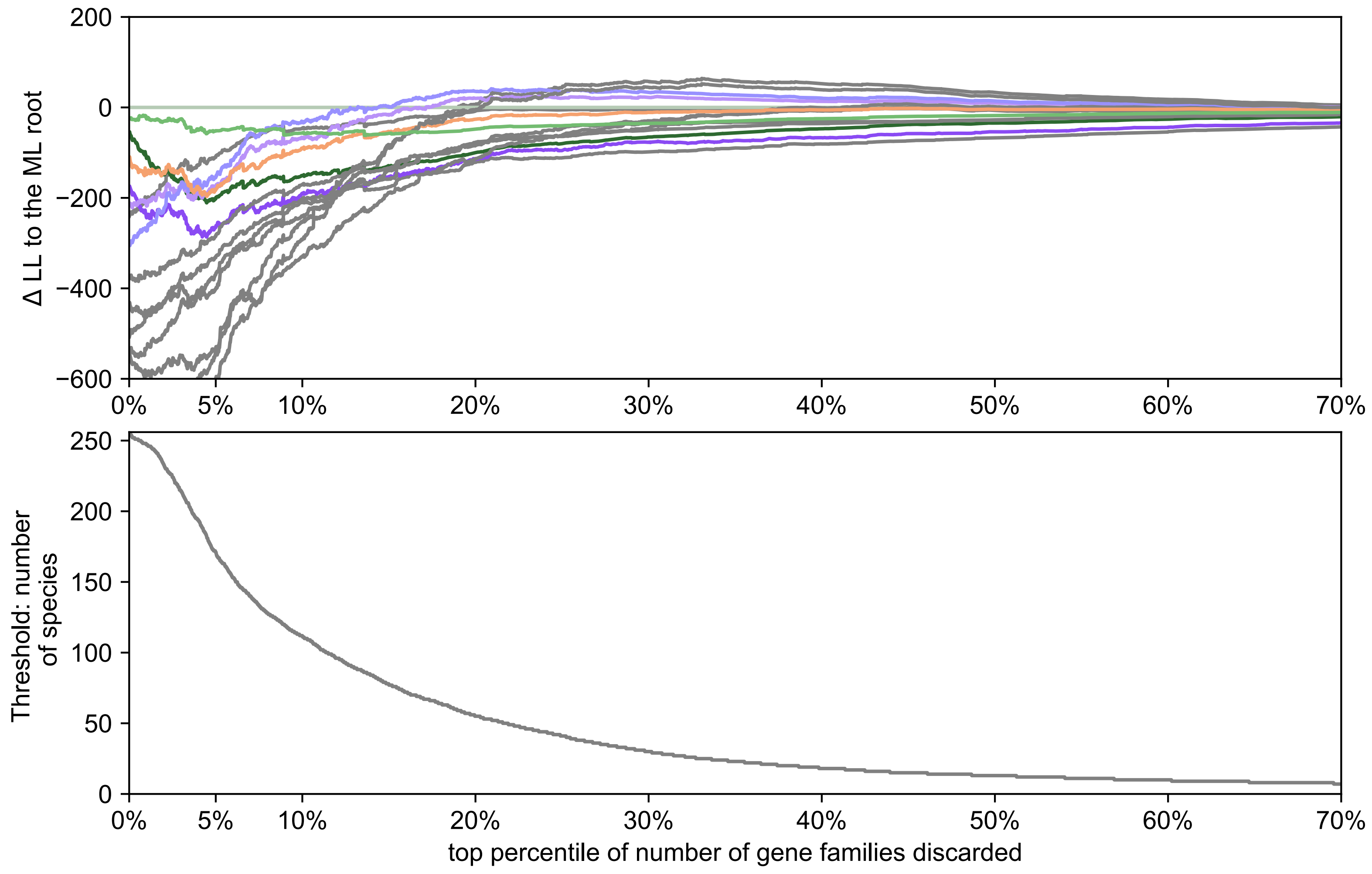


**DPANN: DTL (allowing a different DTL rate for DPANN)**

F.c

by the taxonomic bias score, from biased to balanced

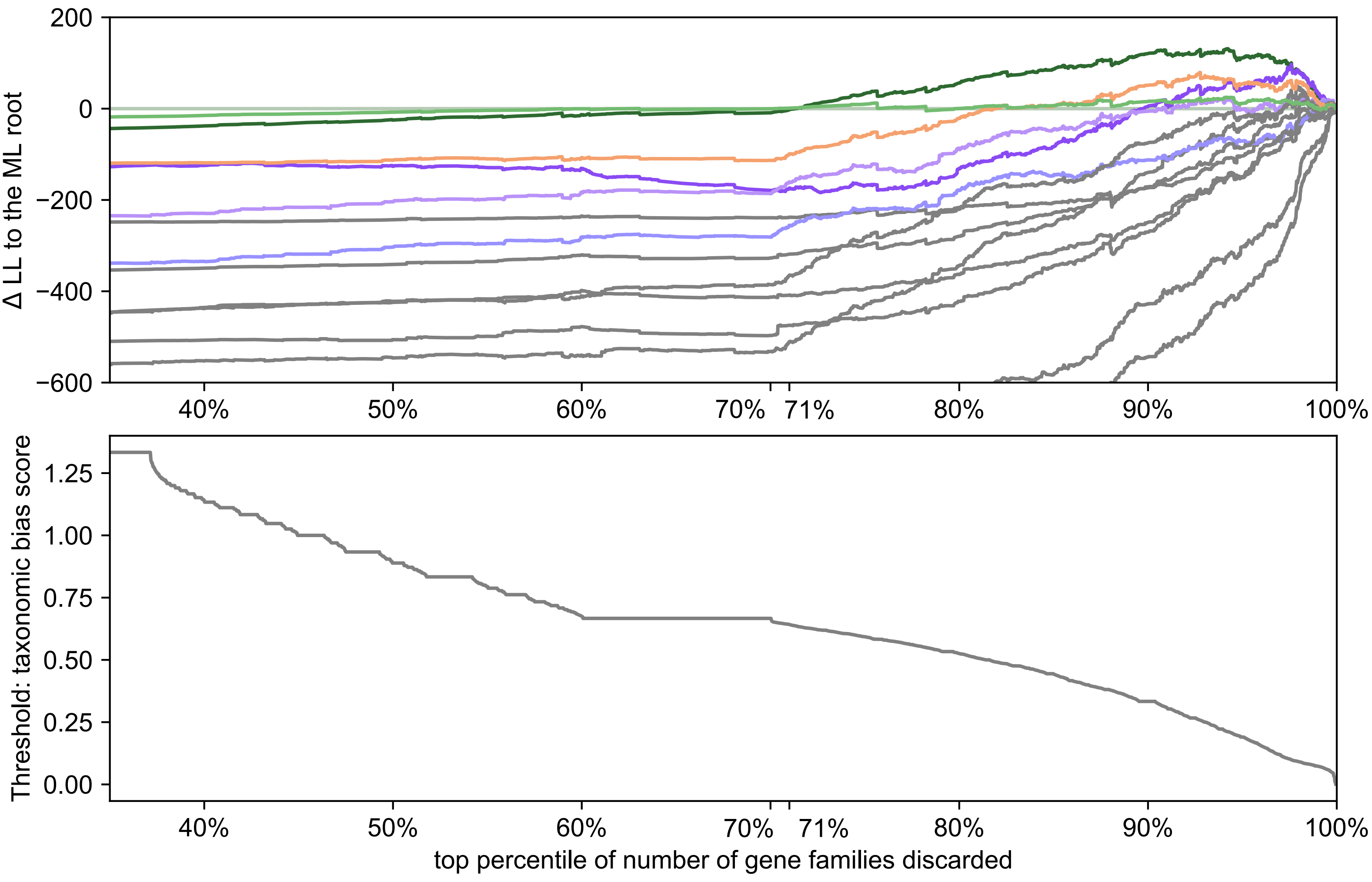


F.d

by the taxonomic bias score, from balanced to biased

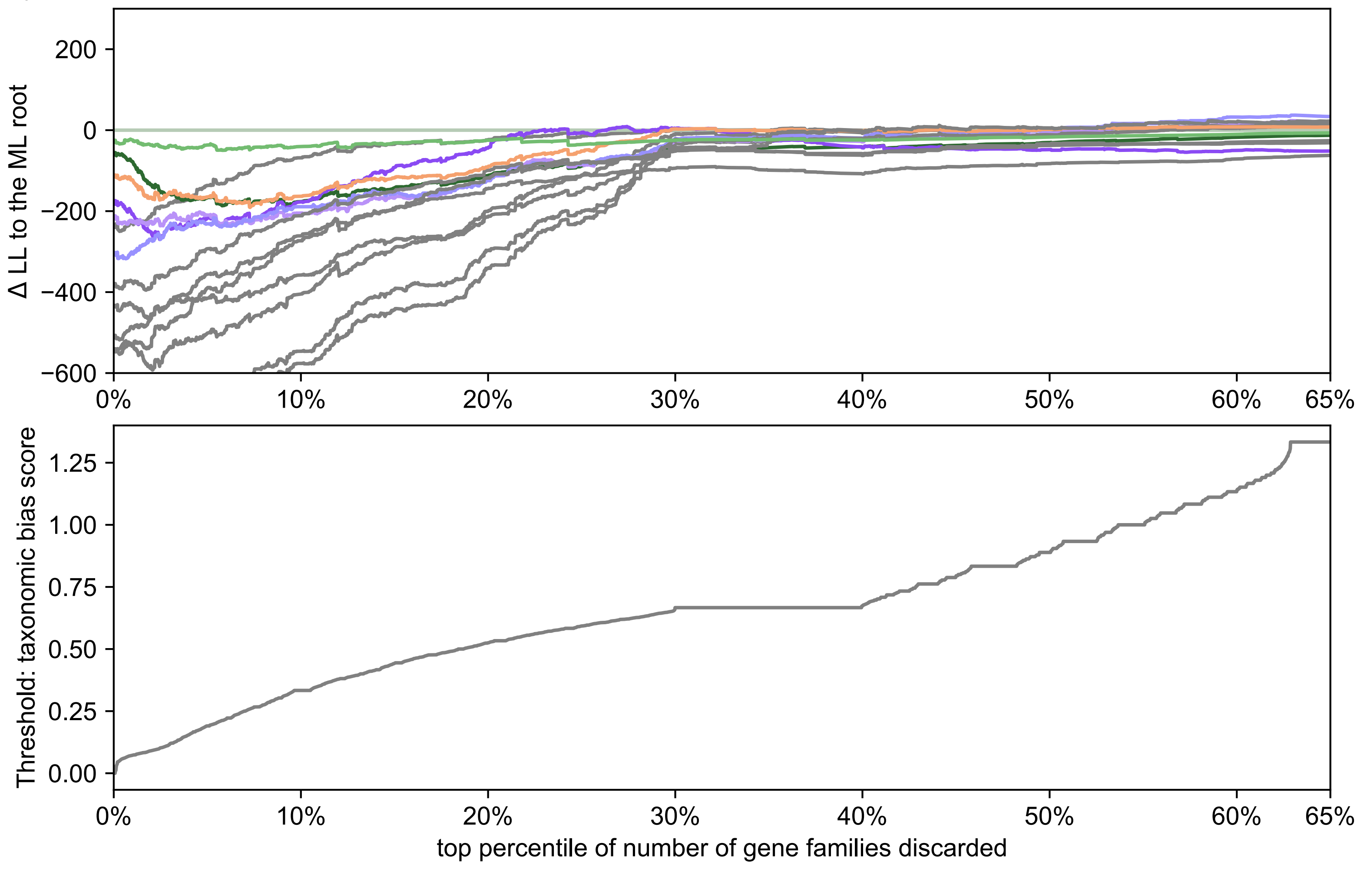

**DTL_br1**

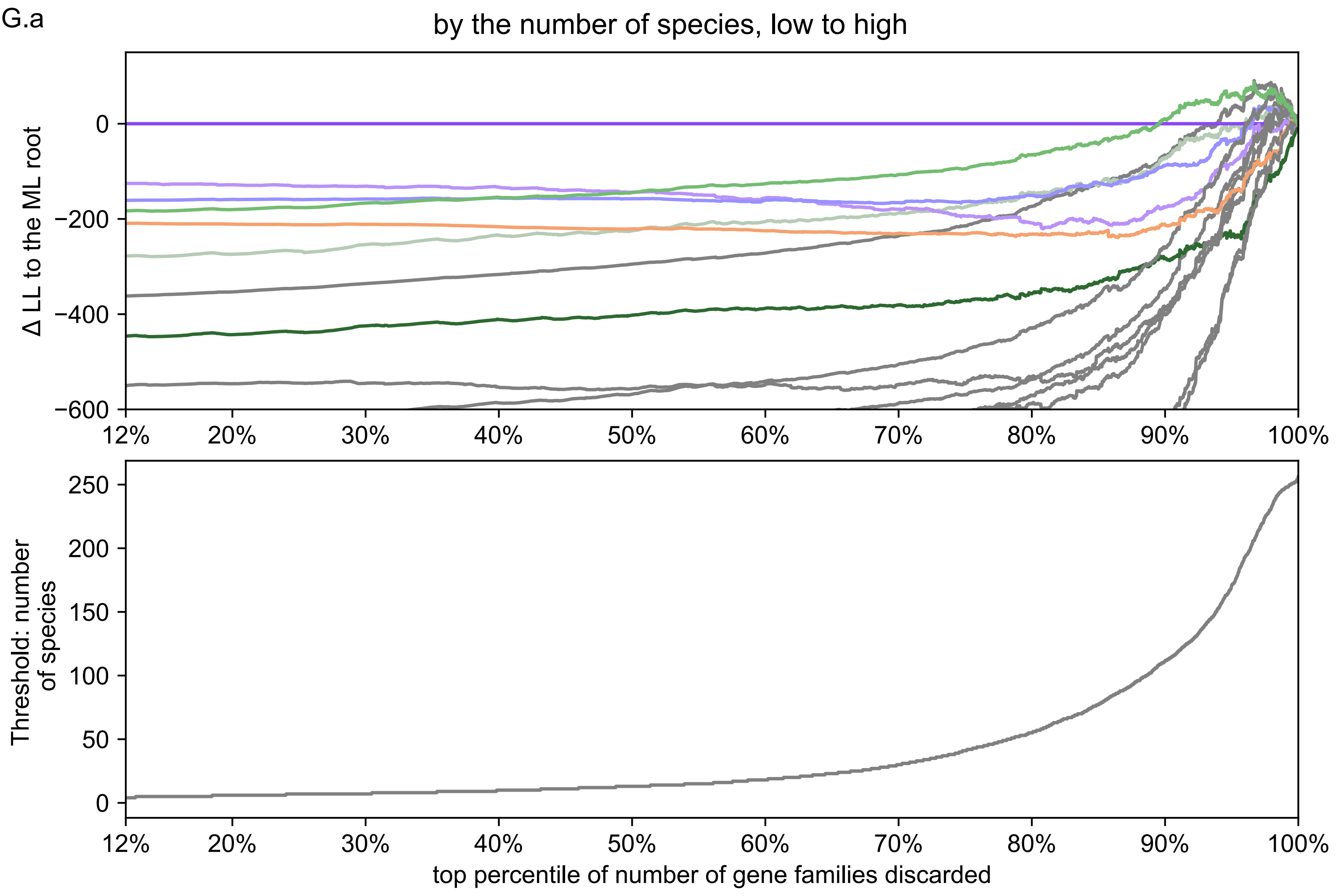


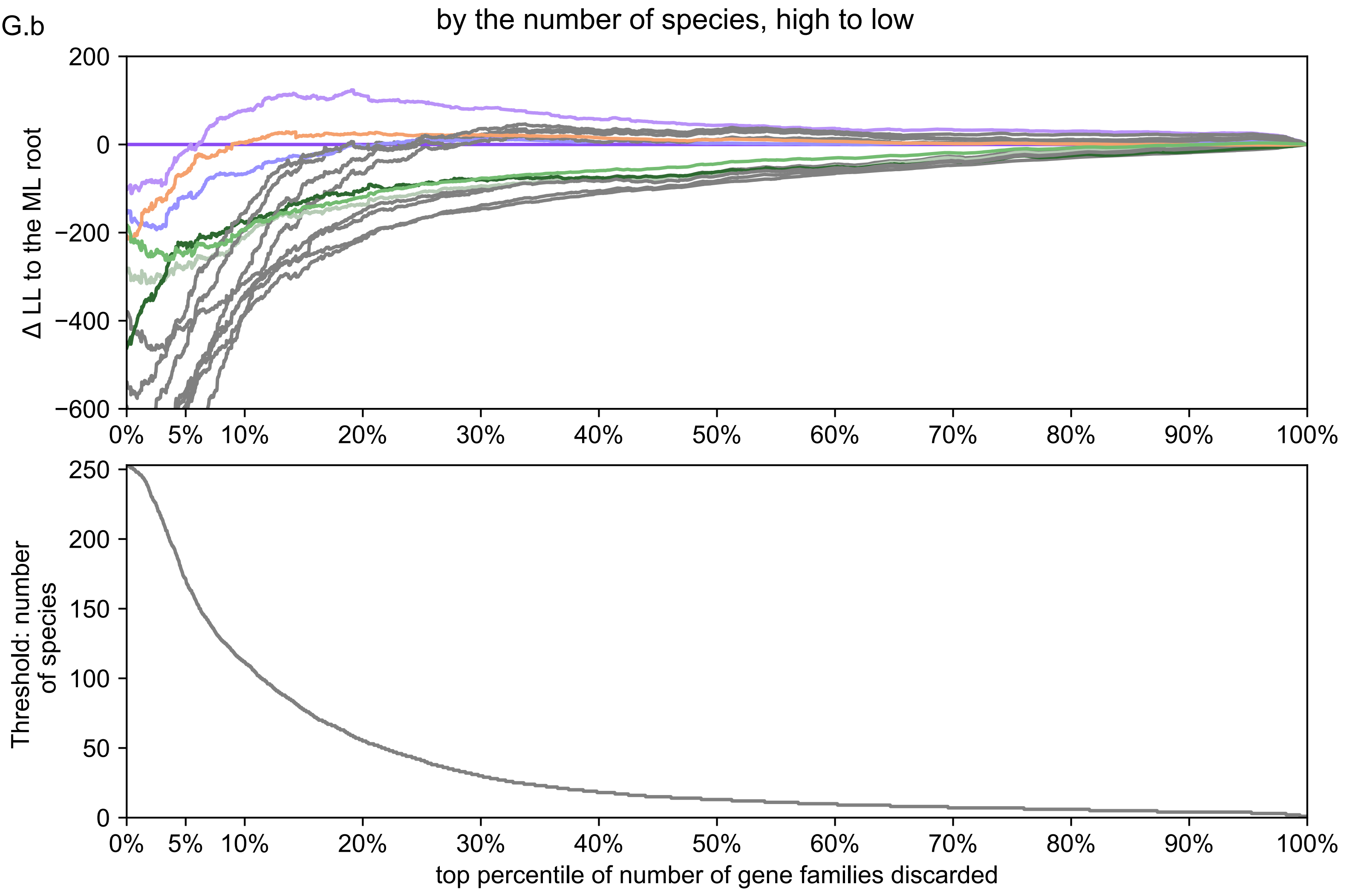


**DTL_br1**

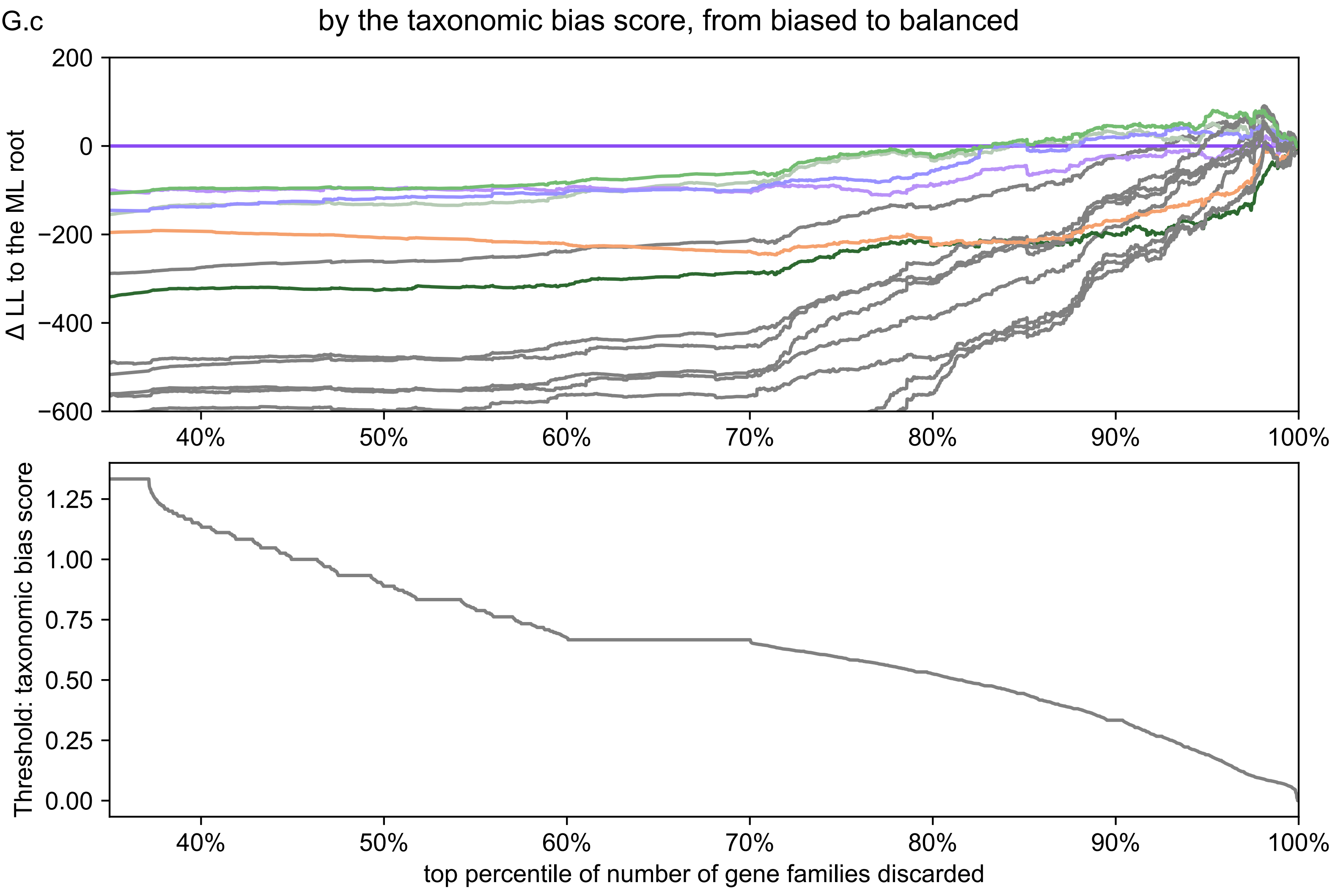


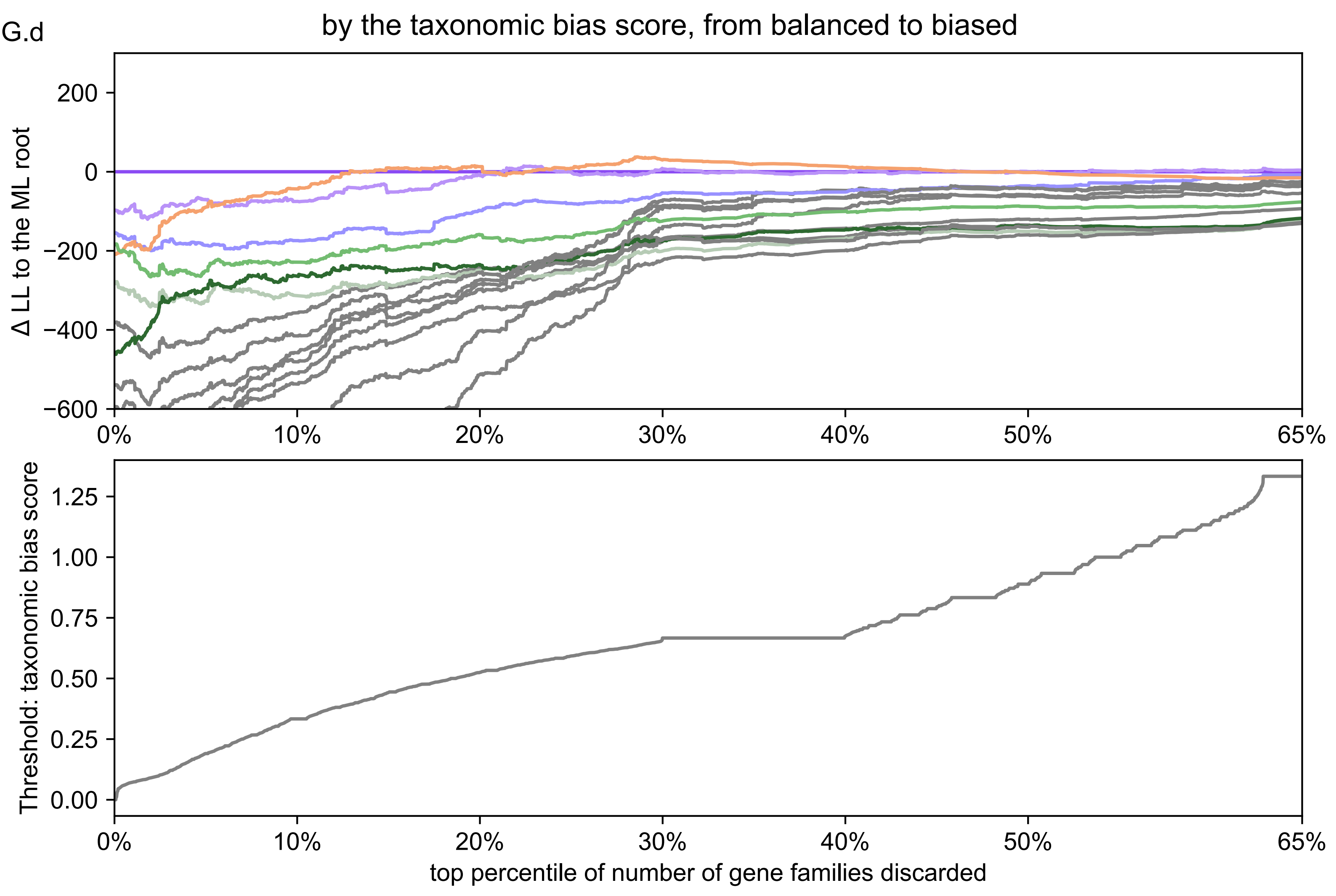

**DTL_br1_O**

H.a by the number of species, low to high

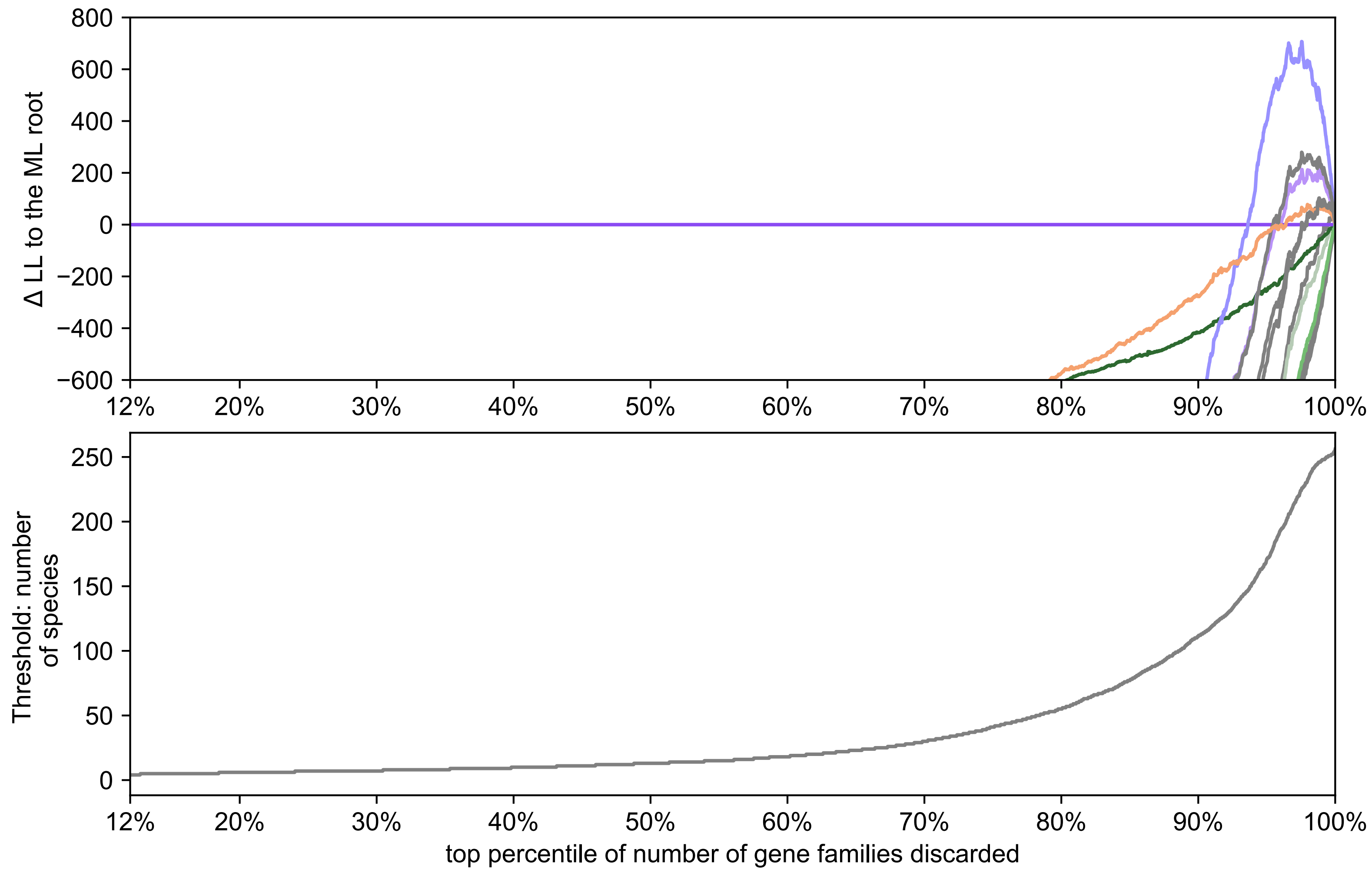


H.b by the number of species, high to low

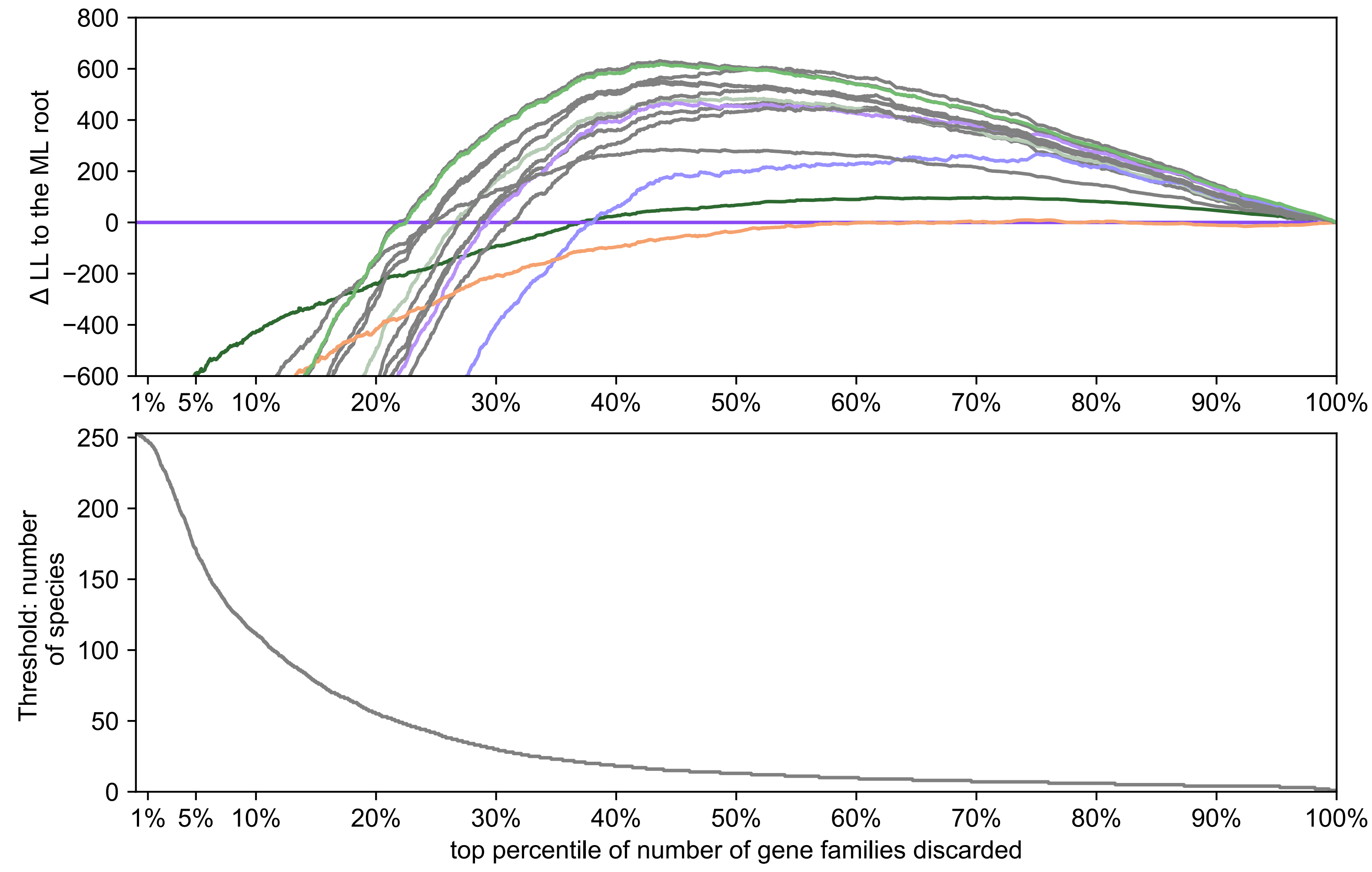


**DTL_br1_O**

H.c by the taxonomic bias score, from biased to balanced

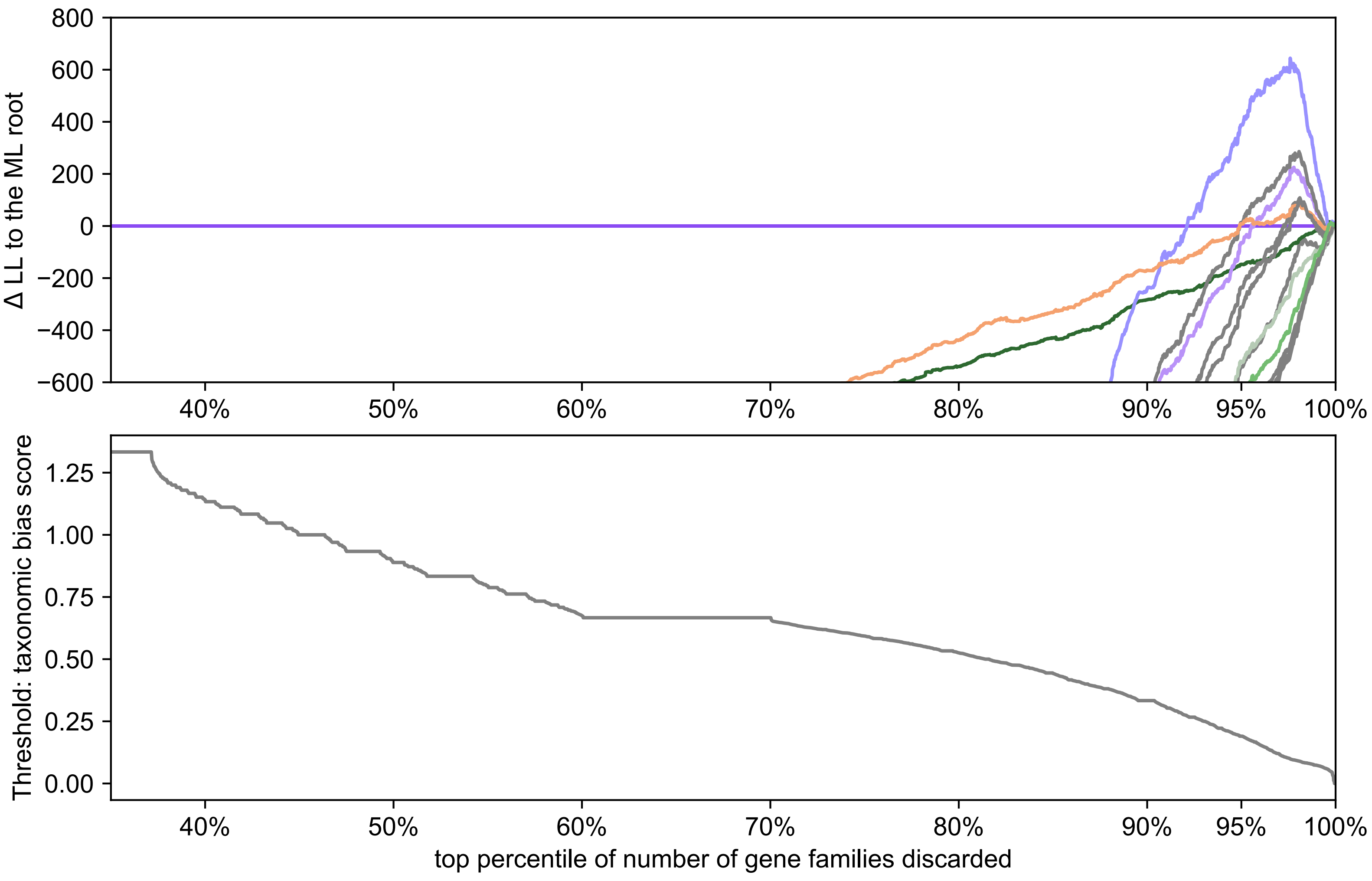


H.d by the taxonomic bias score, from balanced to biased

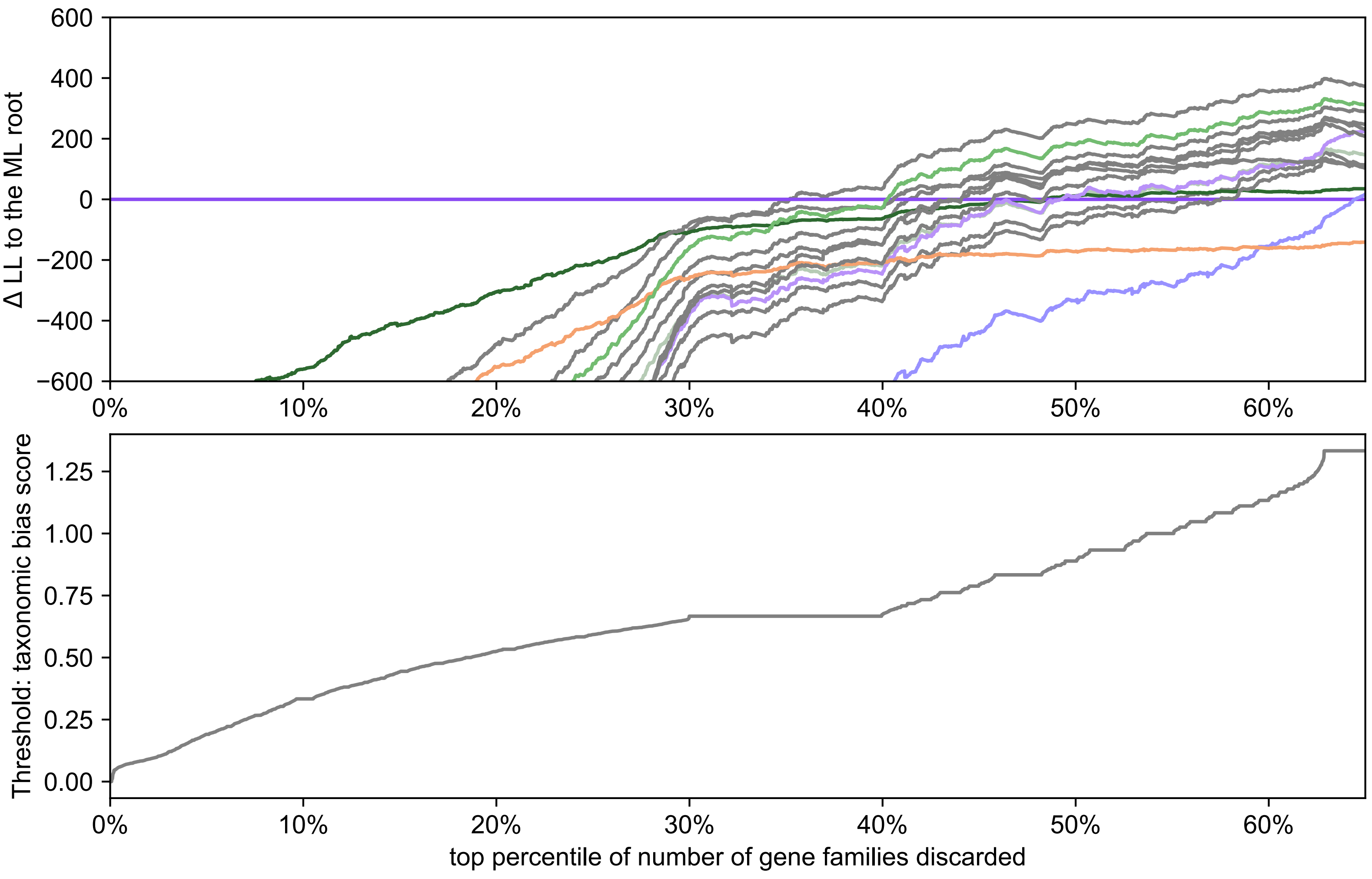

DTL_br2

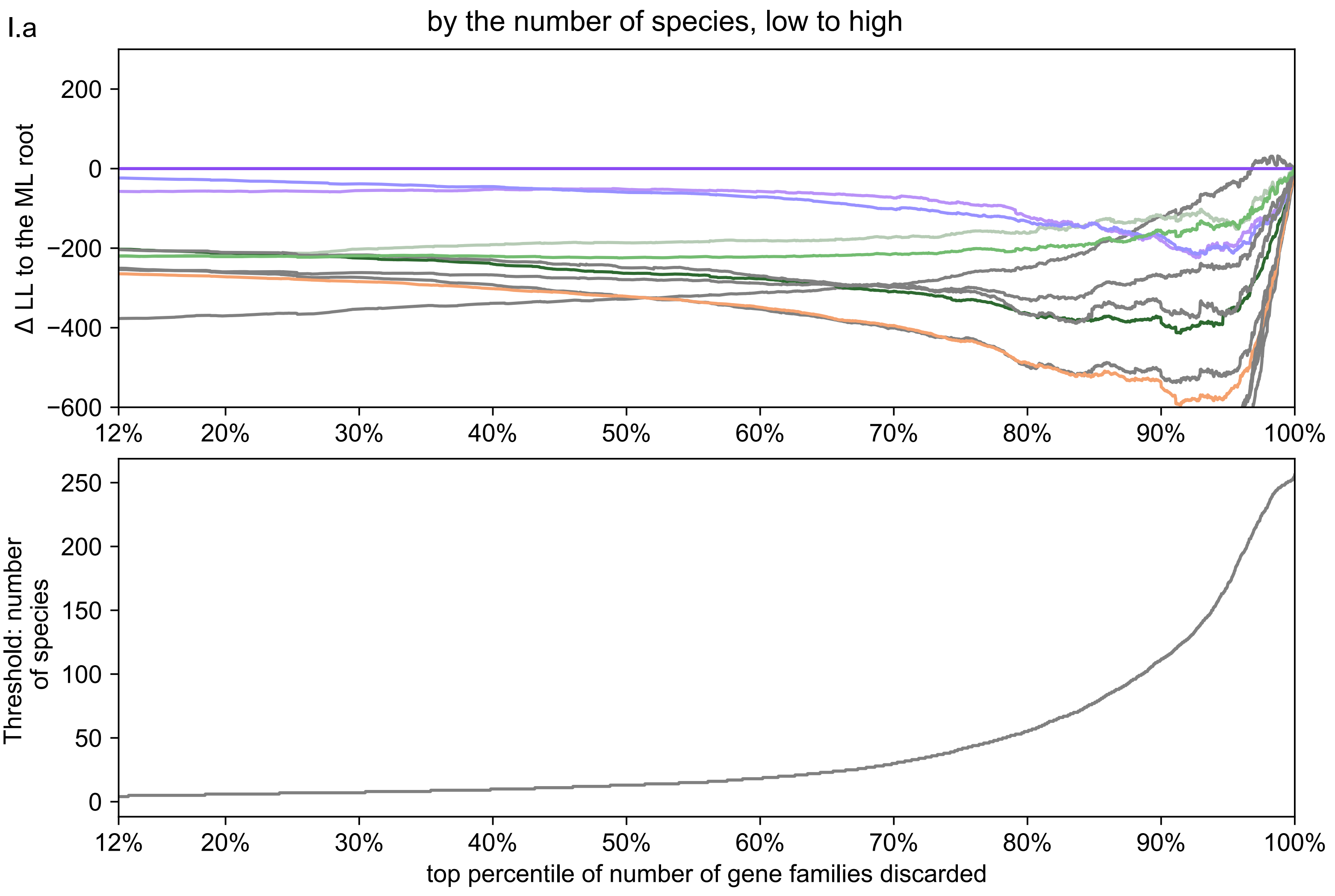


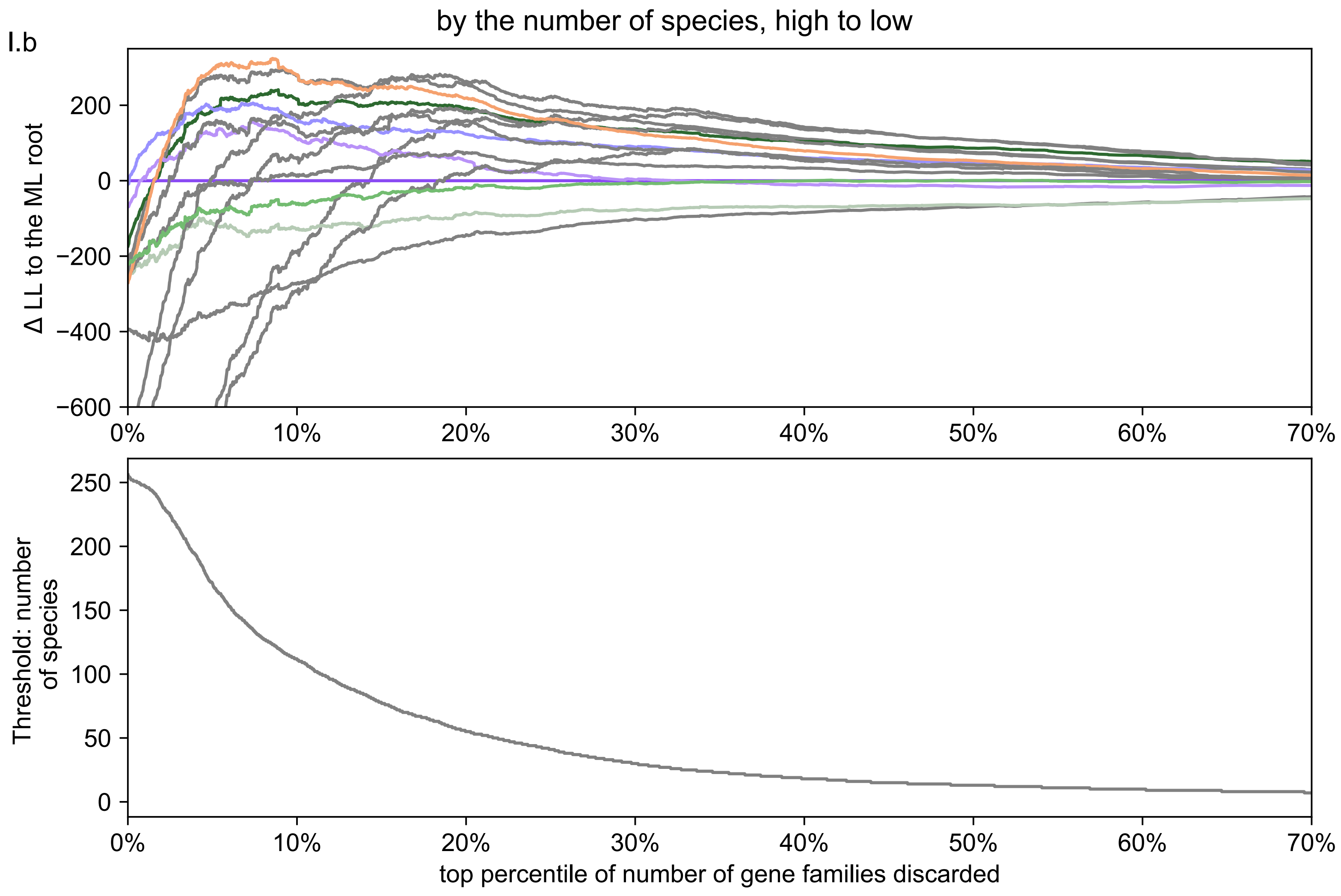


DTL_br2

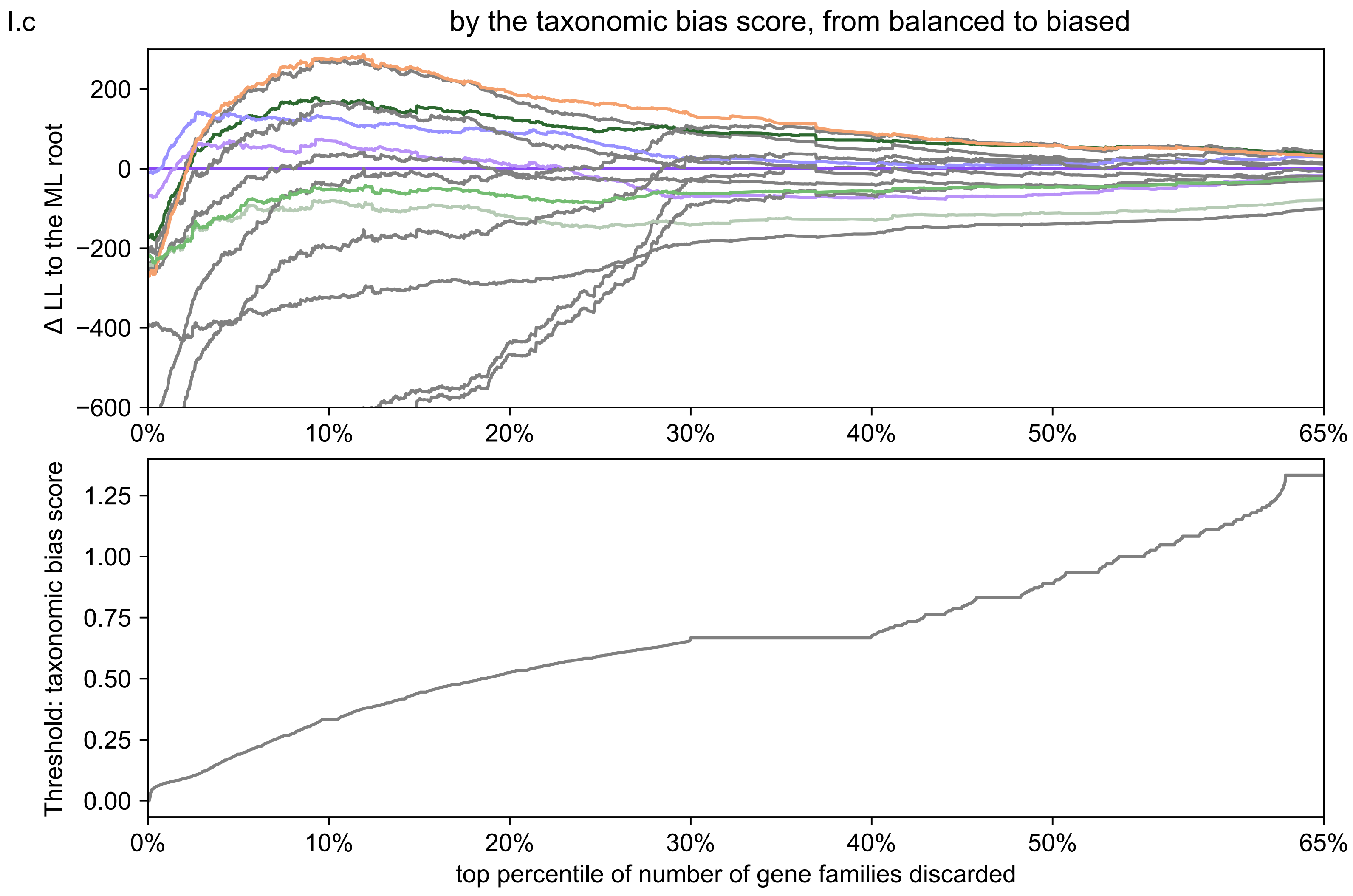


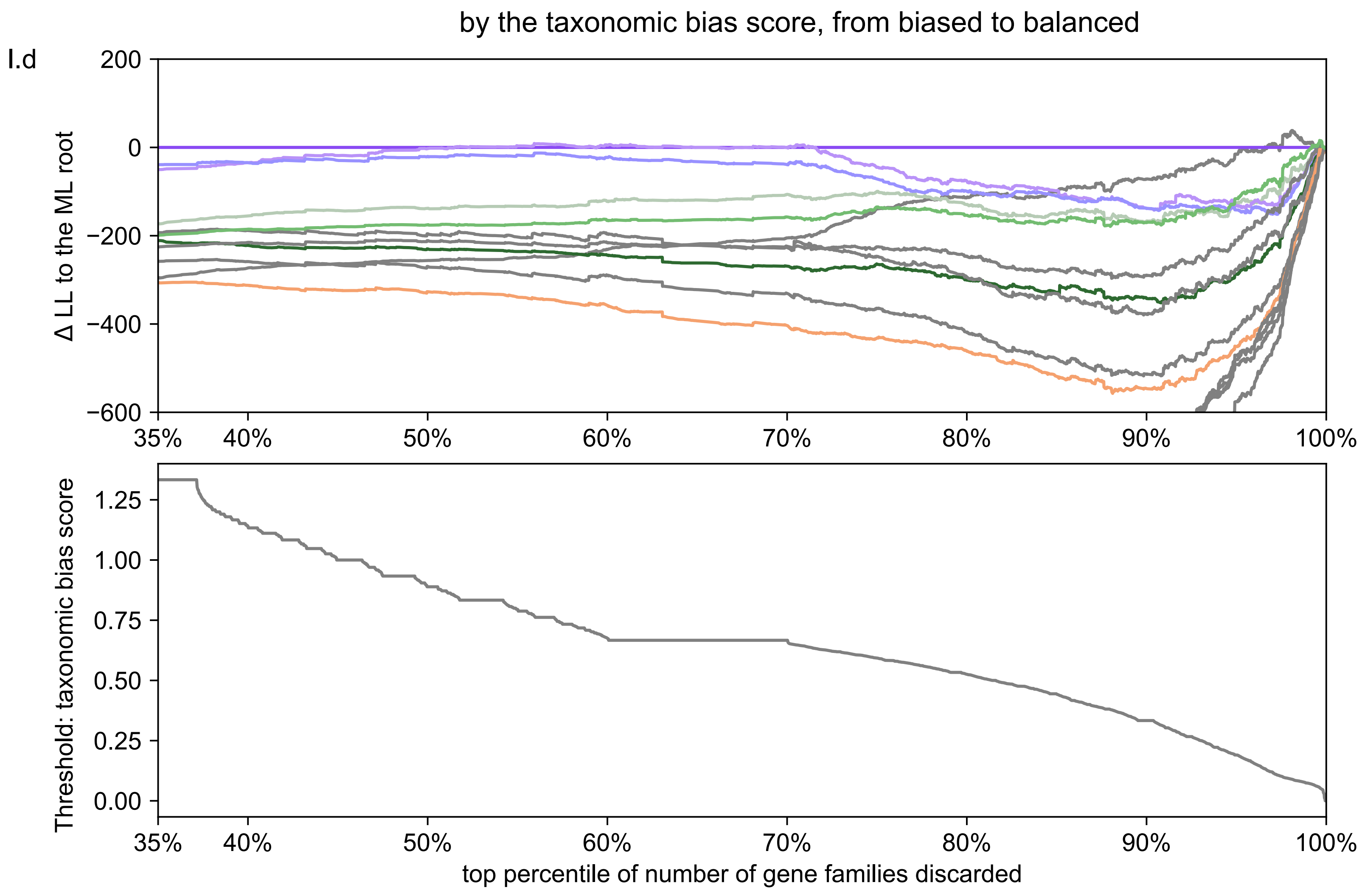


**Fig. S5: Impact of gene family filtering on root likelihoods of different reconciliation models.** We ranked gene families based on the number of species within each gene family and the taxonomic bias score. The lower the score is, the more taxonomically balanced the family is, hence more representative and informative. We then performed analyses to evaluate the impact on root likelihoods of excluding (filtering out) an increasing proportion of the top-ranked gene families for each criterion. The procedure was used in Coleman et al. (2021) and resembled site filtering in traditional phylogenetics, a useful approach to explore the types of gene families that support each root position. Each analysis was performed on the arCOG gene family set based on a different reconciliation model. For each analysis, the top panel plots ΔLL, the difference in log likelihood between a given root and the maximum likelihood root, as a function of filtered gene families. Positive values of ΔLL indicate that a given root has a better likelihood than the overall ML root at that percentile. The second panel displays the distribution of the test quantity and the value of the quantity at each percentile of the filtered gene families. Panel (**A**) maps the colours used in subsequent panels to the position of that root on a schematic unrooted tree; for example, the root between Euryarchaeota and the rest of Archaea corresponds to the dark purple line in panels (**B**) through (**I**). From high to low: families present with the most species are filtered first; from low to high: families present with the fewest species are filtered first. From biased to representative: families with the most biased representation are filtered first; from representative to biased: families with the most balanced representation are filtered first. Model names are given in the panels of each title, and model parameterisations are provided in Table S31.

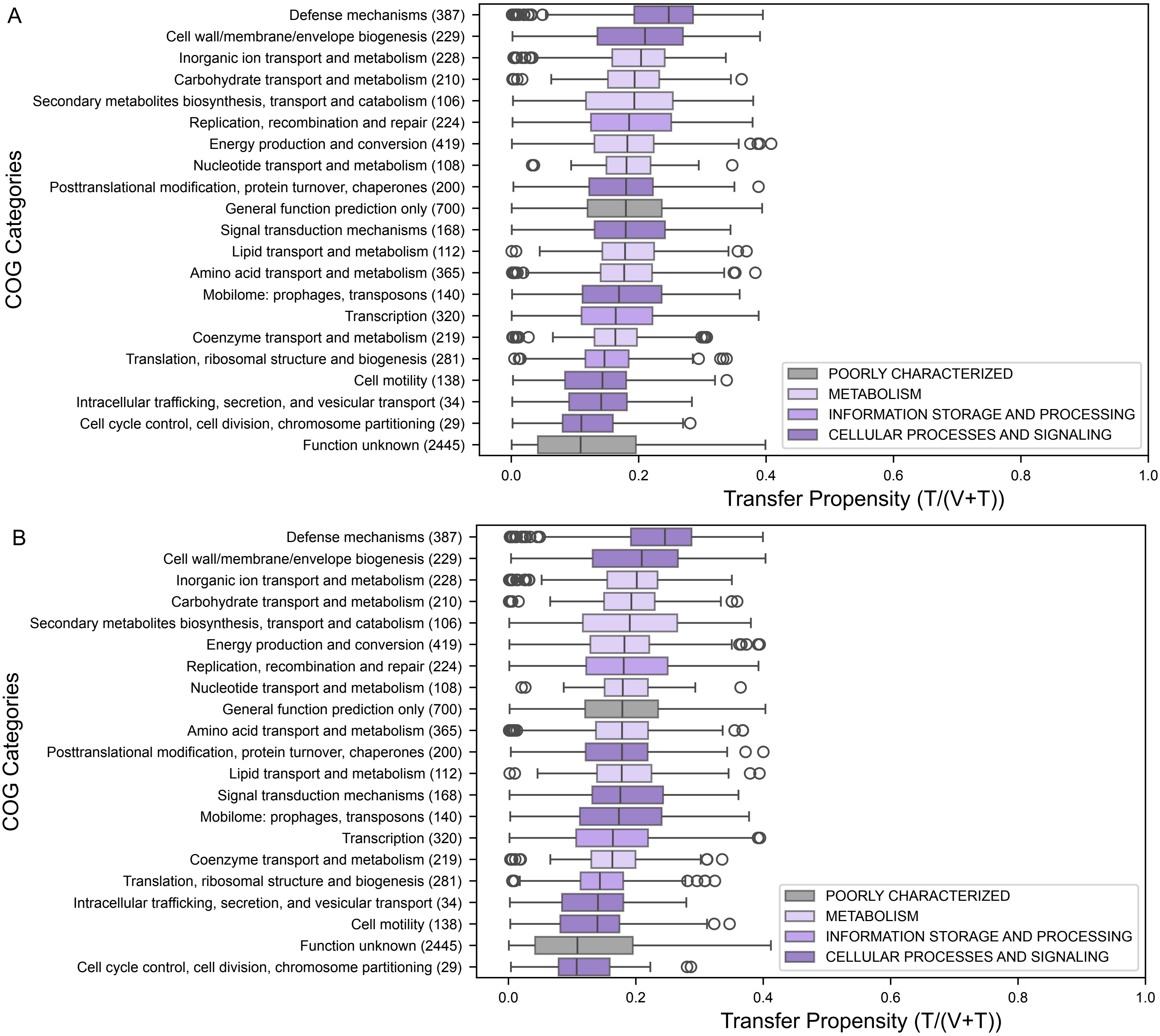


**Fig. S6: Summary of transfer propensity of gene families by COG categories under the Euryarchaeota root (A) and the MHH root (B).** Transfer propensity is the proportion of gene tree branches that are horizontal T/(V+T) for COG gene families. Summary statistics are provided in Table S18. Post-hoc Conover's tests among different gene families by COG categories are provided in Tables S16 and S17.

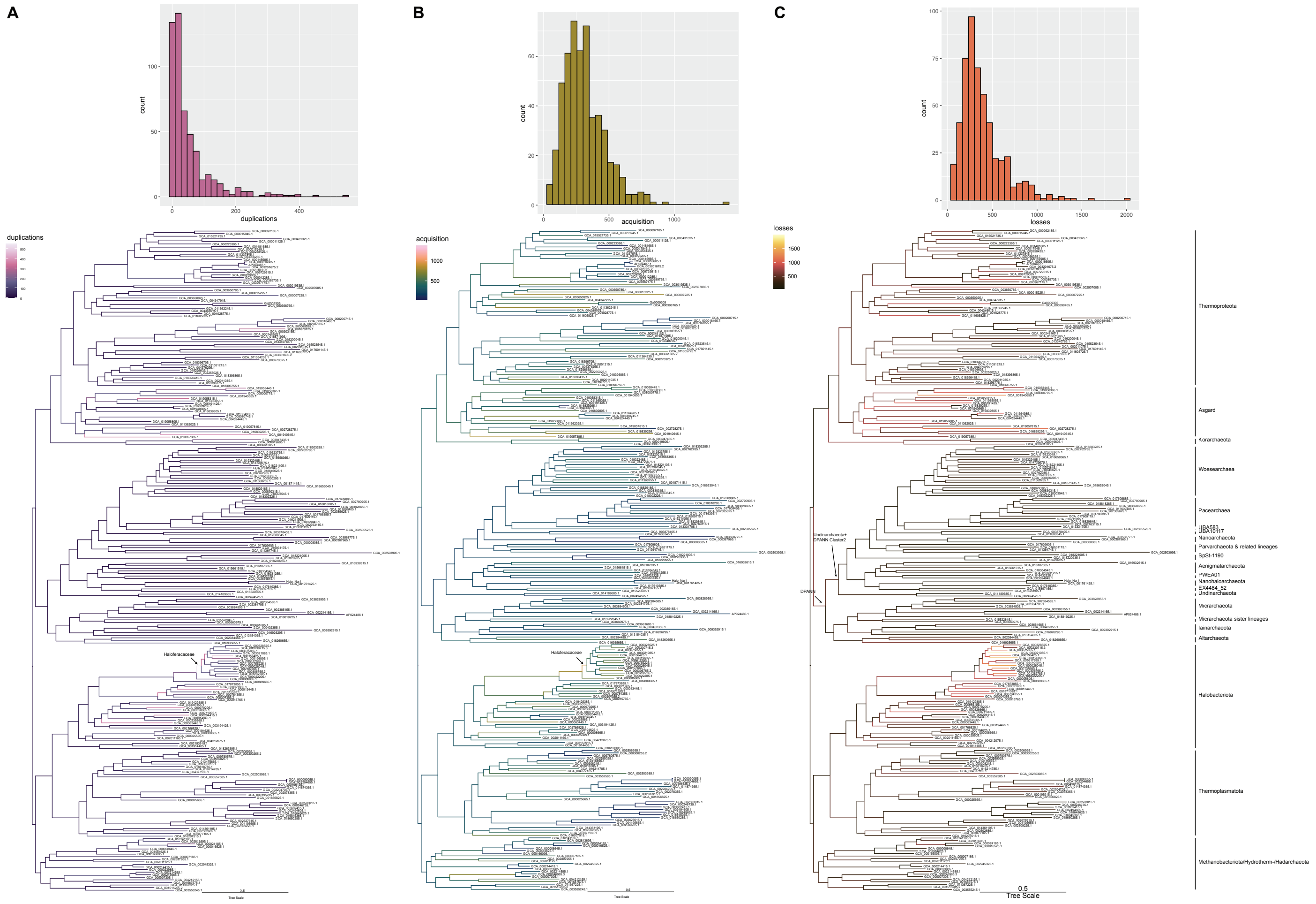


**Fig. S7: Dynamics of gene duplications, acquisitions (transfers + originations), and losses across the archaeal tree of life under the Euryarchaeota root.** (**A–C)** For each event class, the upper panel shows the distribution of per-branch counts (histogram), and the lower panel maps the inferred events onto the archaeal tree; branch colour encodes the number of events. (**A**) Duplications. (**B**) Acquisitions ( transfers + originations). (**C**) Losses.

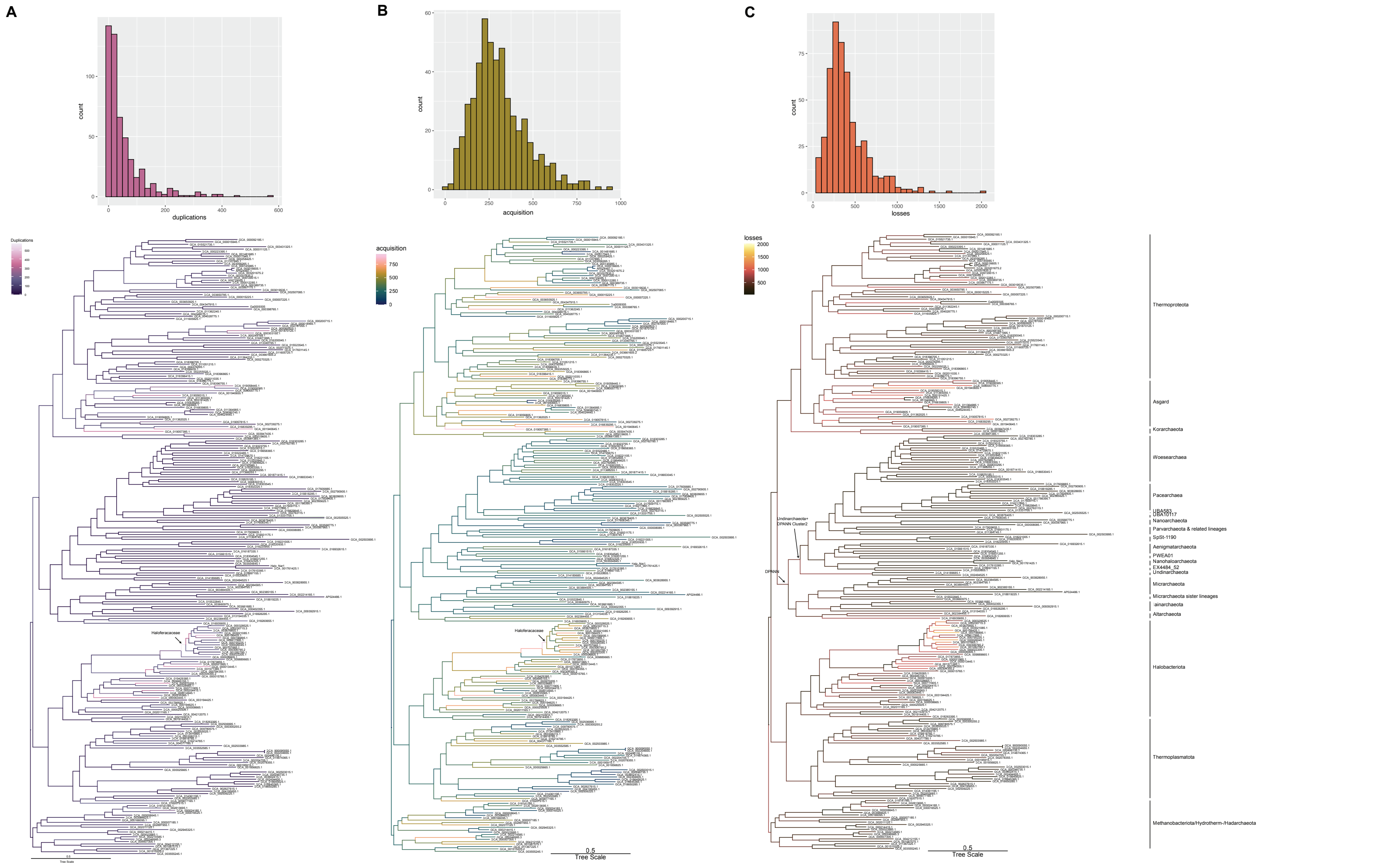


**Fig. S7: Dynamics of gene duplications, acquisitions (transfers + originations), and losses across the archaeal tree of life under the MHH root.** (**A–C)** For each event class, the upper panel shows the distribution of per-branch counts (histogram), and the lower panel maps the inferred events onto the archaeal tree; branch colour encodes the number of events. (**A**) Duplications. (**B**) Acquisitions ( transfers + originations). (**C**) Losses.

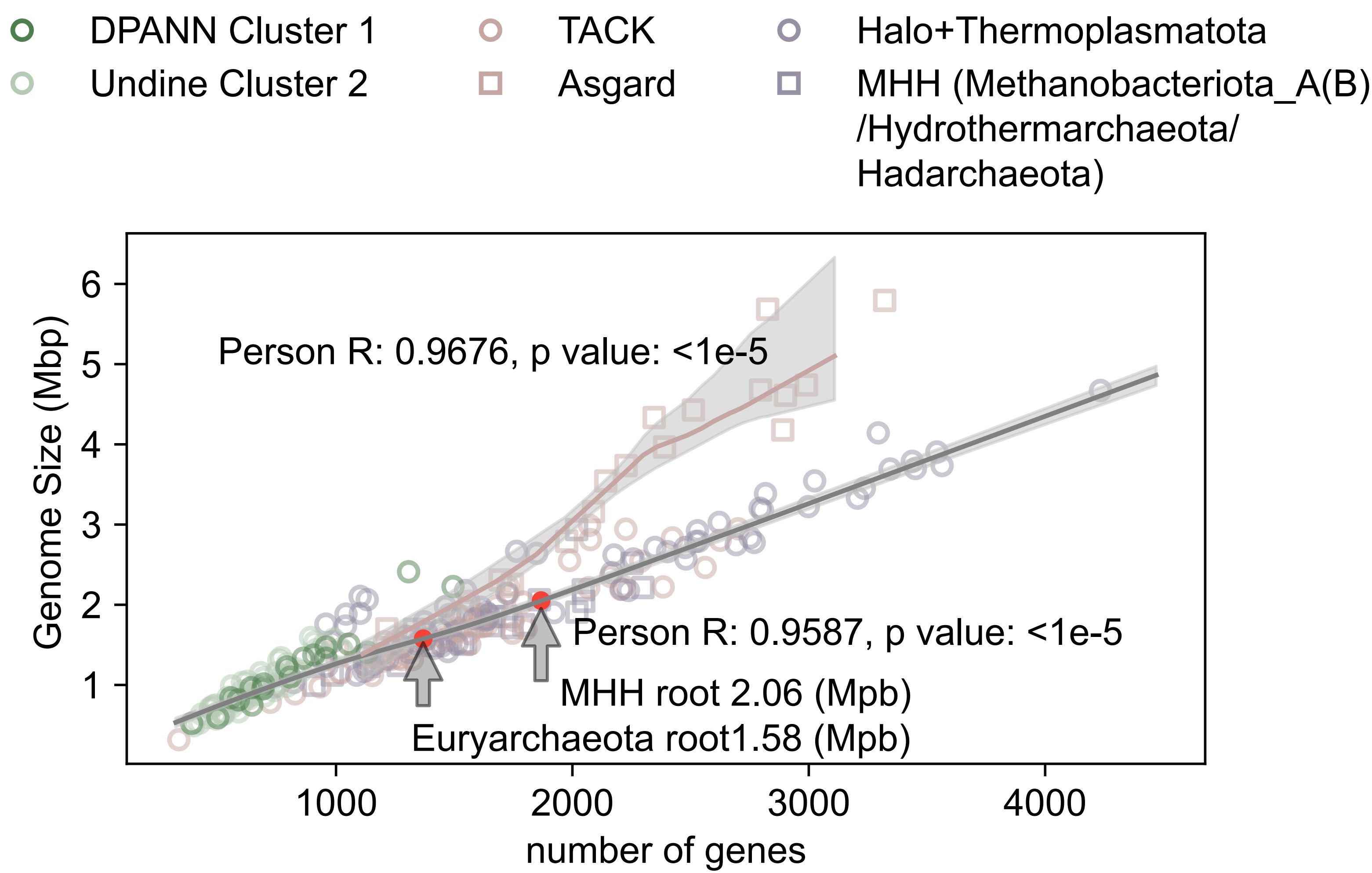


**Fig. S9: Correlation of arCOG gene families and genome size of extant archaea.** Each point is a genome. A locally weighted (LOESS) regression is fitted to non-Asgard archaea, and a separate regression to Asgard archaea, which deviate from the overall trend. These regression models were used to predict ancestral genome size along the archaeal tree of life. Estimates for the last archaeal common ancestor (LACA) under the Euryarchaeota-root and MHH-root placements are shown.

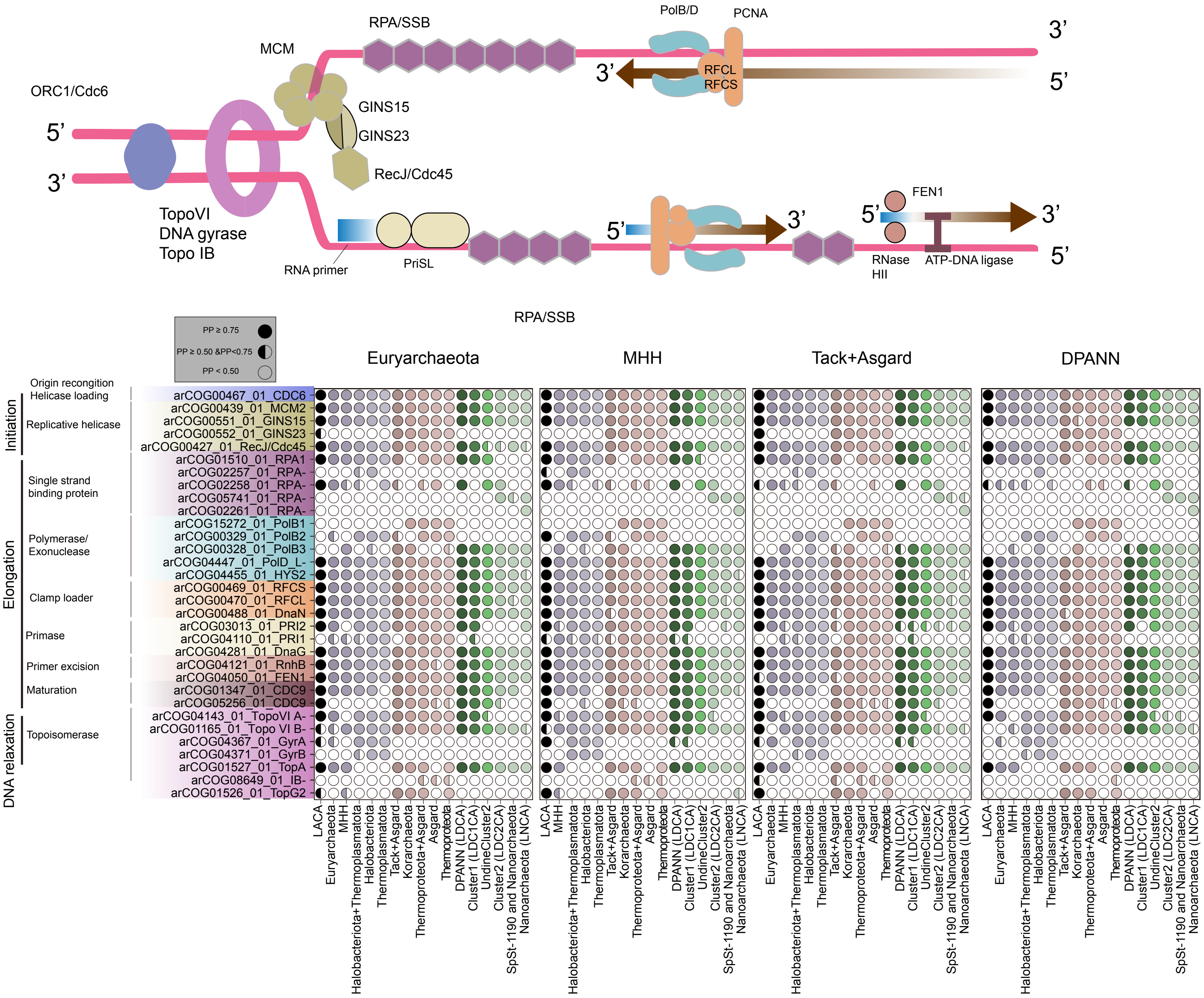


**Fig. S10: Presence posterior probabilities for DNA replication genes in LACA and key archaeal nodes under different roots.** Shown are posterior probabilities (PP) of gene presence for DNA replication machinery inferred for the last archaeal common ancestor (LACA) and selected nodes under four root placements: Euryarchaeota, MHH, TACK+Asgard, and DPANN. Symbols encode support: filled circles for PP ≥ 0.75 (strong), half circles for 0.5 ≤ PP < 0.75 (moderate).

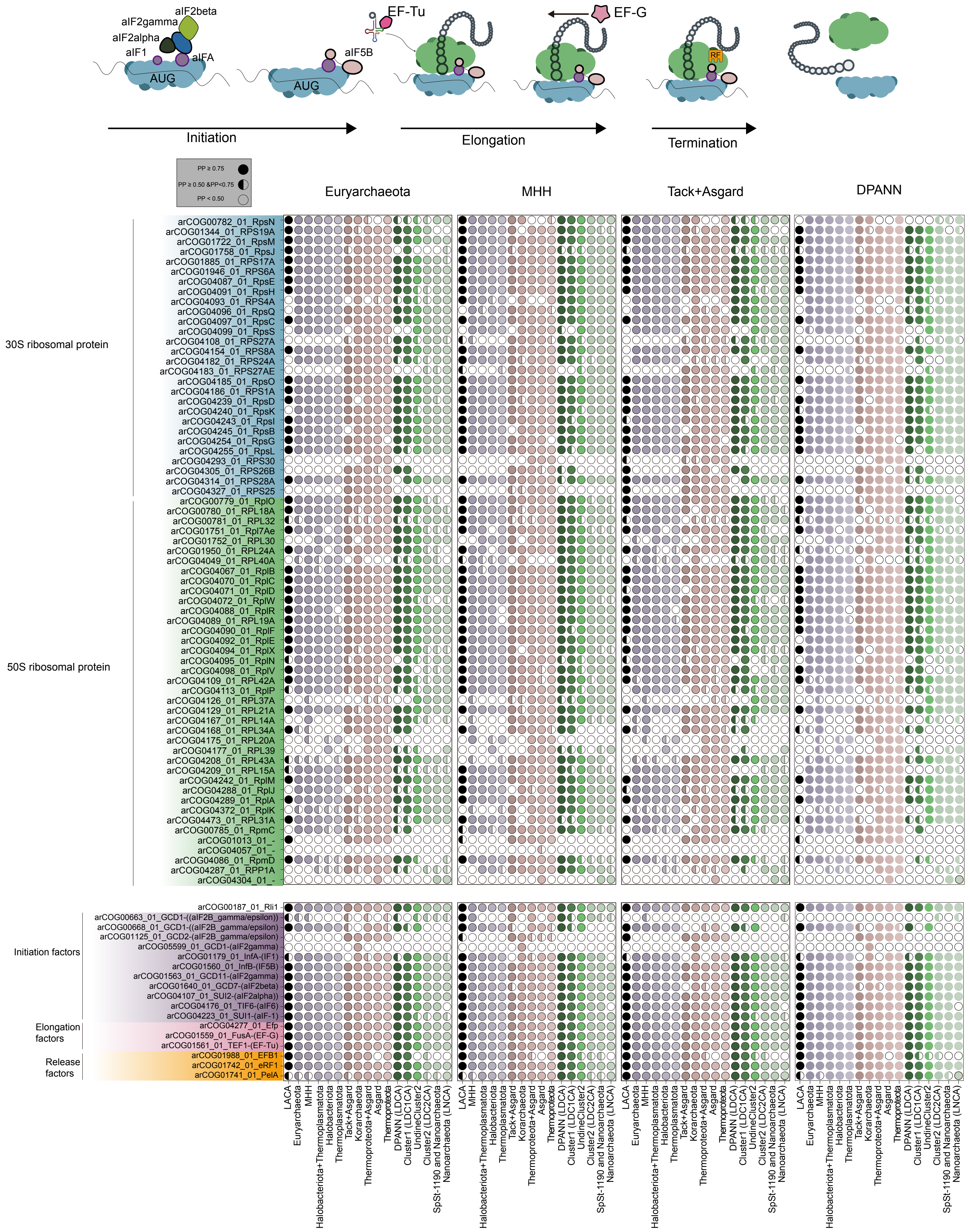


**Fig. S11: Presence posterior probabilities for translation genes in LACA and key archaeal nodes under different roots.** Shown are posterior probabilities (PP) of gene presence for translation inferred for the last archaeal common ancestor (LACA) and selected nodes under four root placements: Euryarchaeota, MHH, TACK+Asgard, and DPANN. Symbols encode support: filled circles for PP ≥ 0.75 (strong), half circles for 0.5 ≤ PP < 0.75 (moderate).

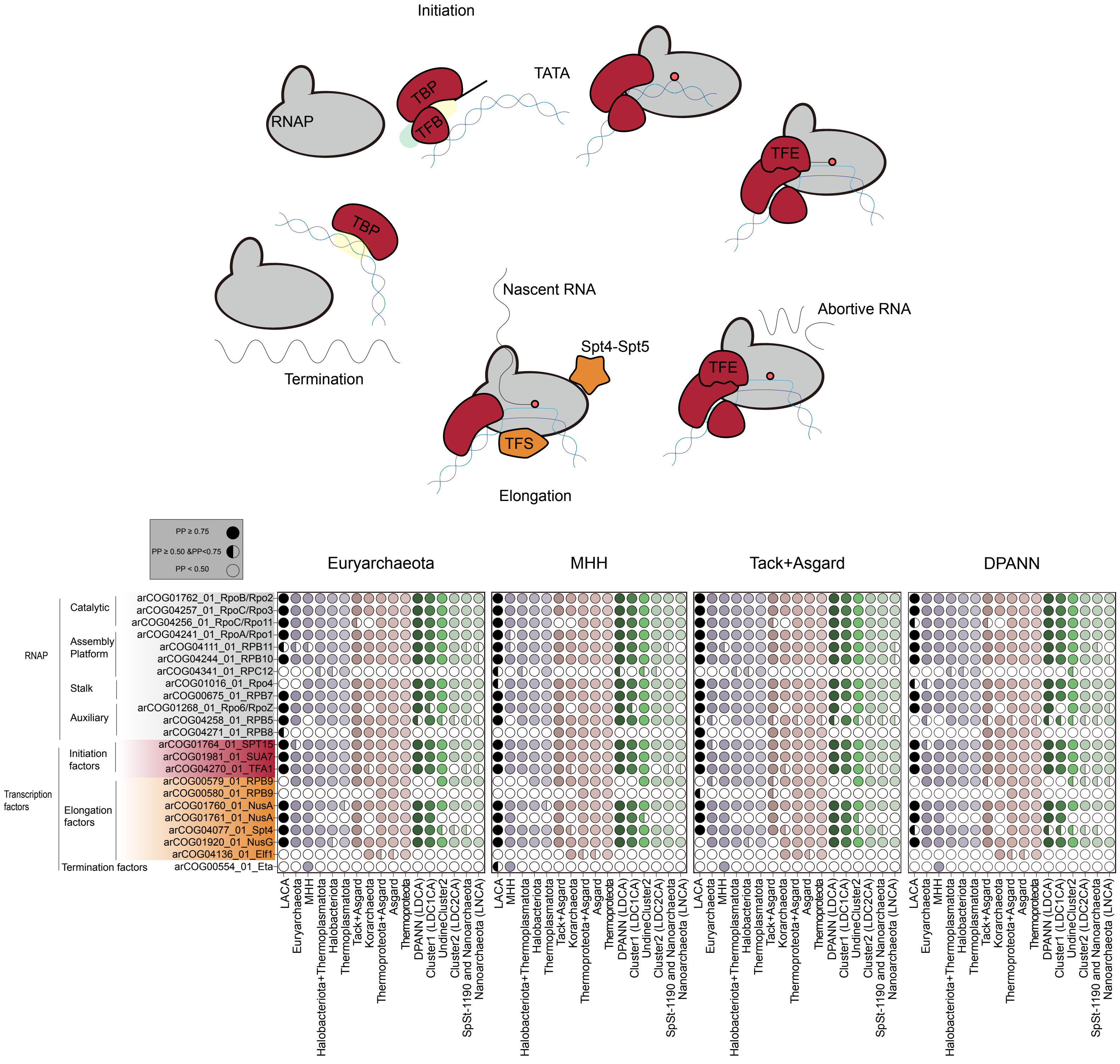


**Fig. S12: Presence posterior probabilities for transcription genes in LACA and key archaeal nodes under different roots.** Shown are posterior probabilities (PP) of gene presence for transcription inferred for the last archaeal common ancestor (LACA) and selected nodes under four root placements: Euryarchaeota, MHH, TACK+Asgard, and DPANN. Symbols encode support: filled circles for PP ≥ 0.75 (strong), half circles for 0.5 ≤ PP < 0.75 (moderate).

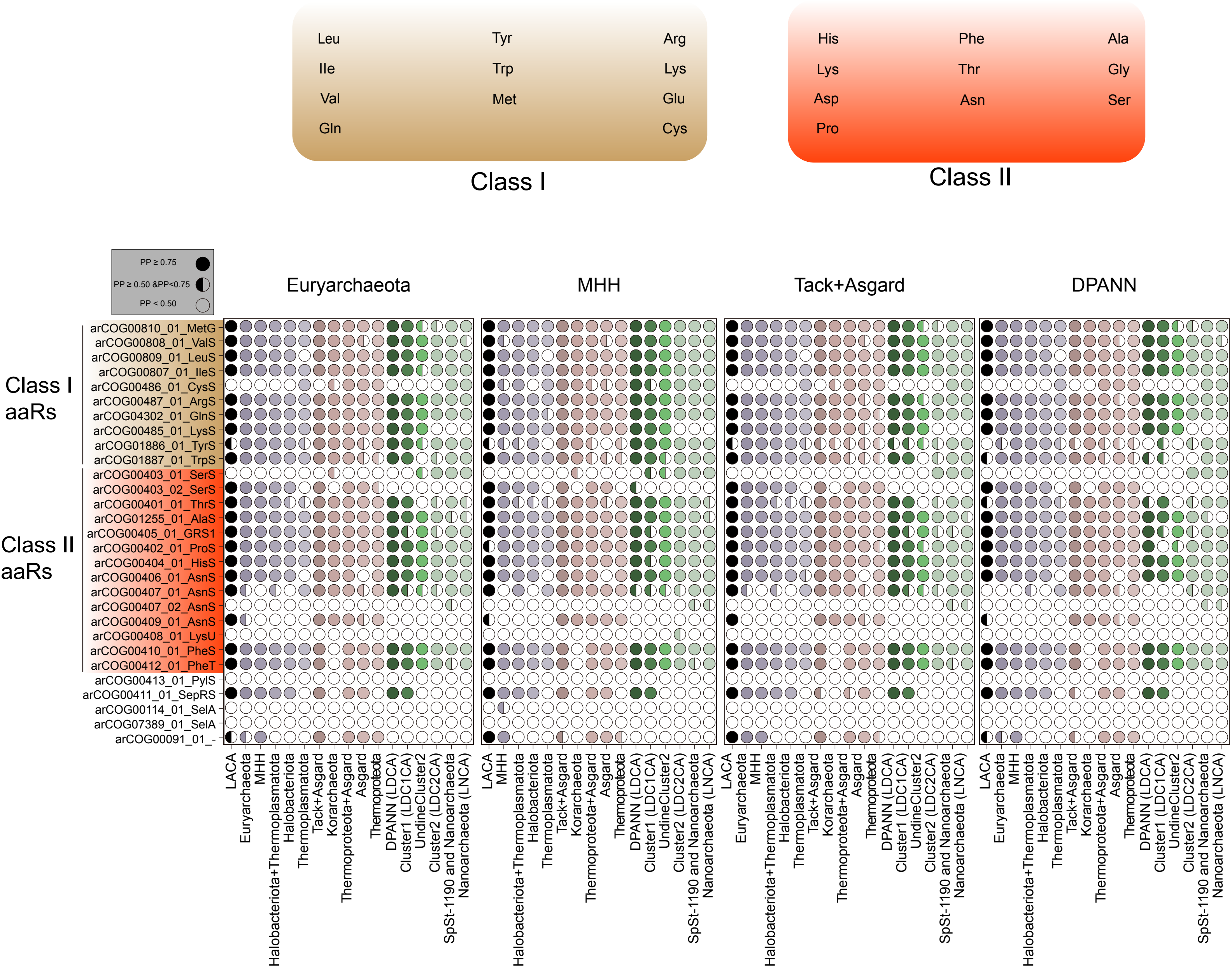


**Fig. S13: Presence posterior probabilities for tRNA synthetase genes in LACA and key archaeal nodes under different roots.** Shown are posterior probabilities (PP) of gene presence for tRNA synthetases inferred for the last archaeal common ancestor (LACA) and selected nodes under four root placements: Euryarchaeota, MHH, TACK+Asgard, and DPANN. Symbols encode support: filled circles for PP ≥ 0.75 (strong), half circles for 0.5 ≤ PP < 0.75 (moderate).

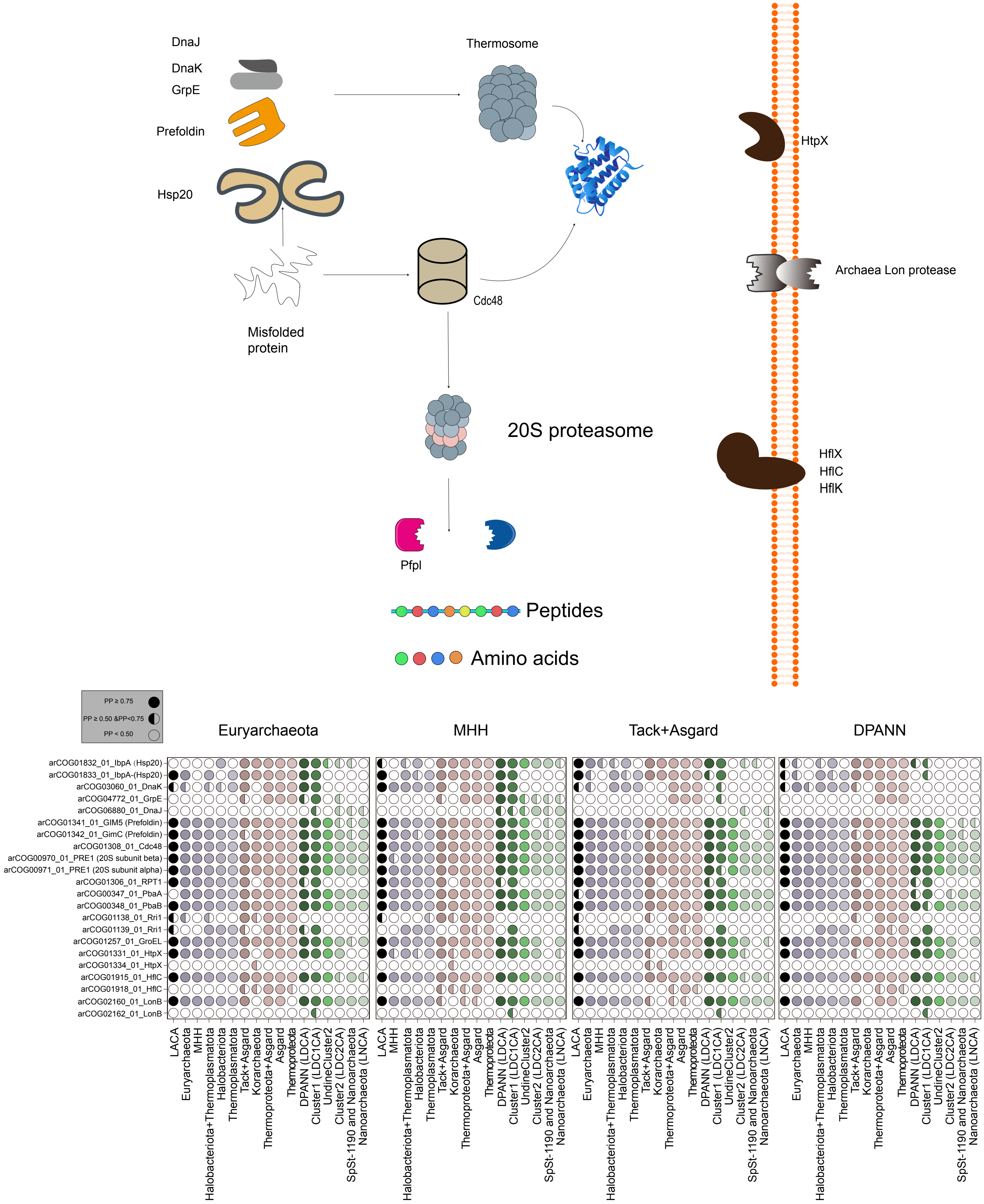


**Fig. S14: Presence posterior probabilities for post-translational genes in LACA and key archaeal nodes under different roots**. Shown are posterior probabilities (PP) of gene presence for post-translational genes inferred for the last archaeal common ancestor (LACA) and selected nodes under four root placements: Euryarchaeota, MHH, TACK+Asgard, and DPANN. Symbols encode support: filled circles for PP ≥ 0.75 (strong), half circles for 0.5 ≤ PP < 0.75 (moderate).

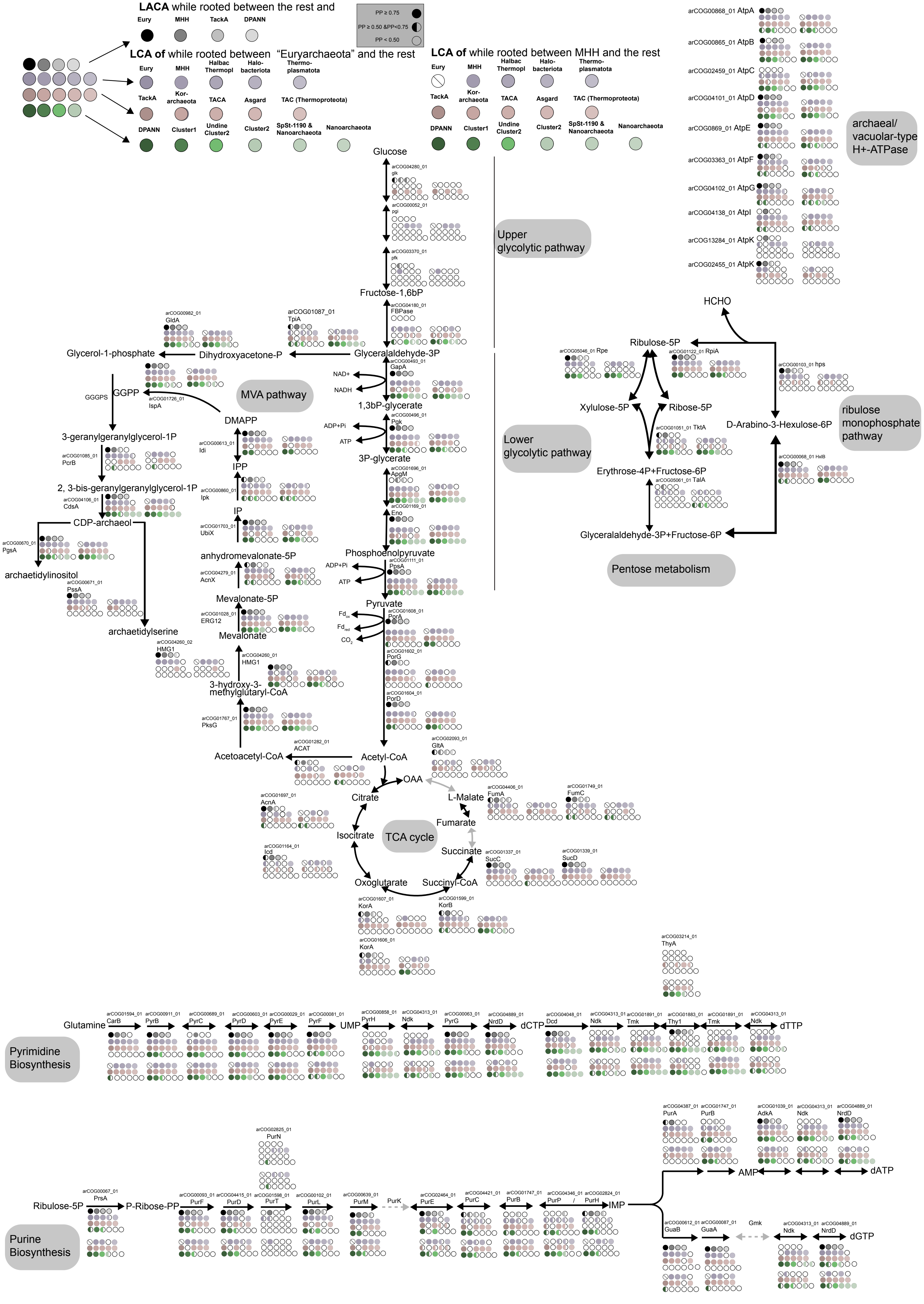


**Fig. S15: Presence posterior probabilities for key metabolic genes of interest in LACA and key archaeal nodes under different roots.** Shown are posterior probabilities (PP) of gene presence for metabolic genes of interest inferred for the last archaeal common ancestor (LACA) under four root placements: Euryarchaeota, MHH, TACK+Asgard, and DPANN. Symbols encode support: filled circles for PP ≥ 0.75 (strong), half circles for 0.5 ≤ PP < 0.75 (moderate). Other ancestors of interest were only shown under the Euryarchaeota and MHH root, which received support from the best-fitting reconciliation model.

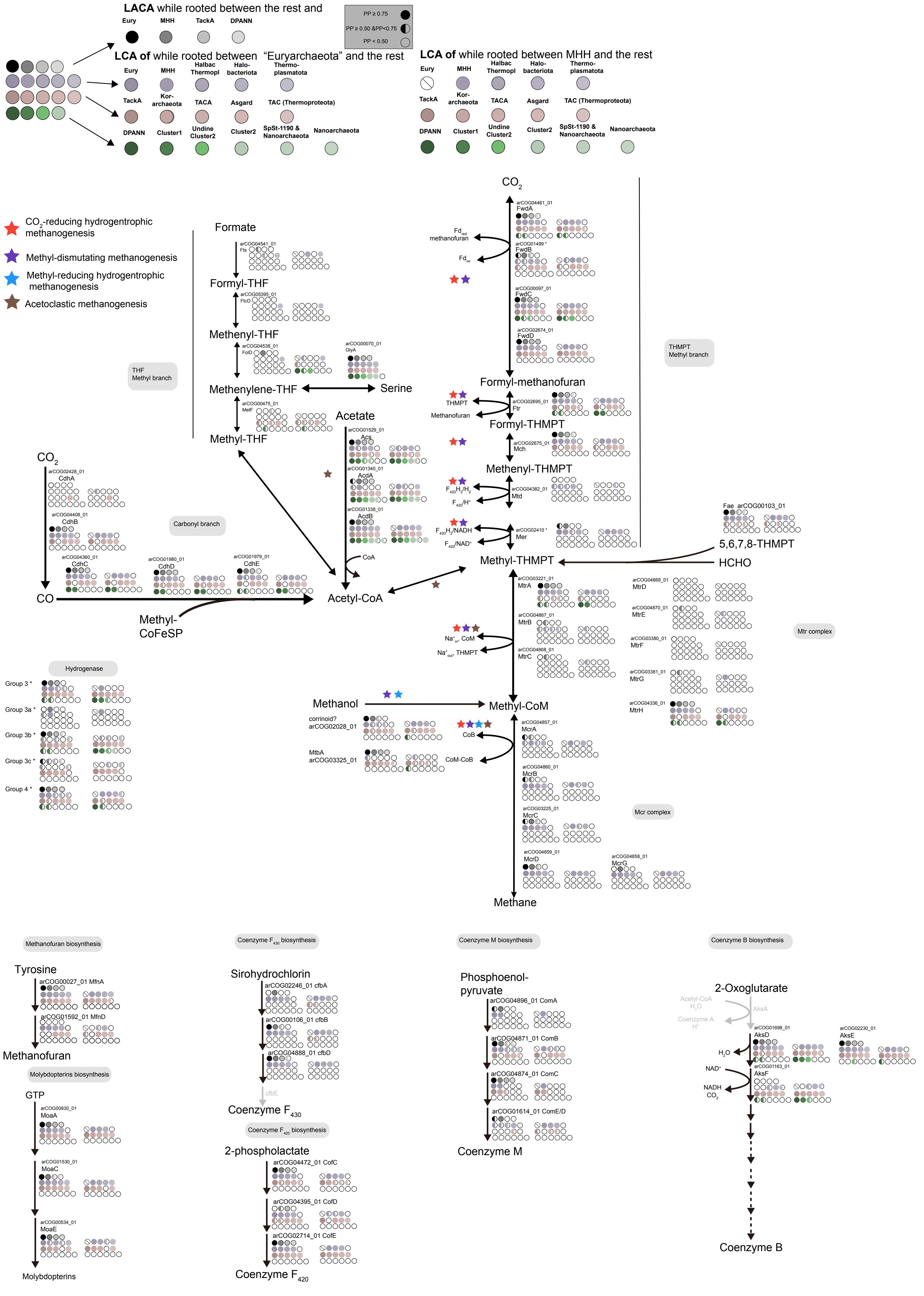


**Fig. S16: Presence posterior probabilities for genes related to the Wood-Ljundahl pathway in LACA and key archaeal nodes under different roots.** Shown are posterior probabilities (PP) of gene presence for metabolic genes of interest inferred for the last archaeal common ancestor (LACA) under four root placements: Euryarchaeota, MHH, TACK+Asgard, and DPANN. Symbols encode support: filled circles for PP ≥ 0.75 (strong), half circles for 0.5 ≤ PP < 0.75 (moderate). Other ancestors of interest were only shown under the Euryarchaeota and MHH root, which received support from the best-fitting reconciliation model. PP of arCOGs denoted with an asterisk can be found in Table S26-S27.

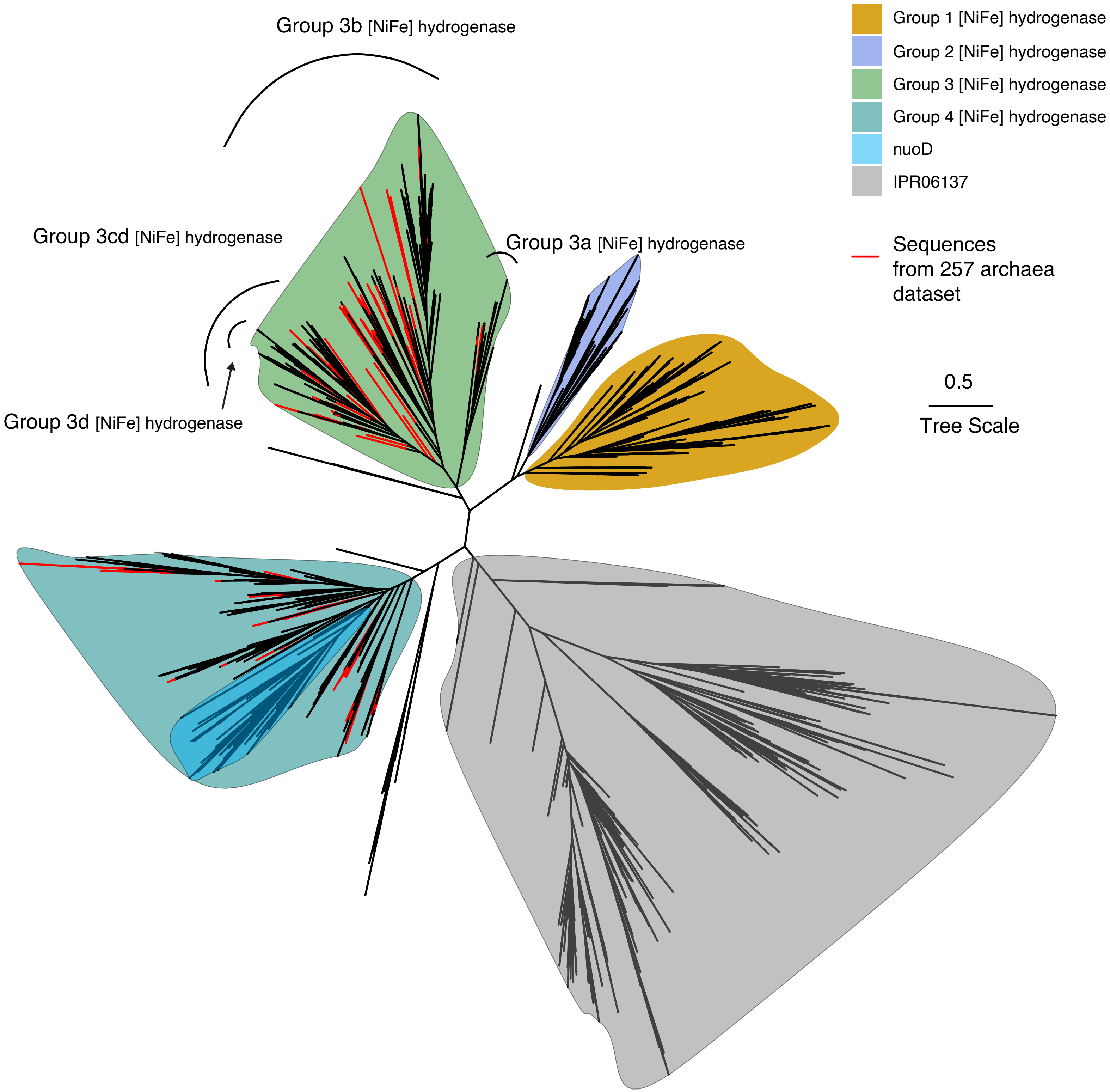


**Fig. S17: Phylogenetic analyses of homologs of [NiFe] hydrogenase under the best-fitting substitution model, LG+R10.** The alignment has 1818 sequences with 194 amino acid sites. Different groups of hydrogenase were assigned based on reference sets. Sequences coloured in red are from the 257-taxa set and were used for reconciliations.

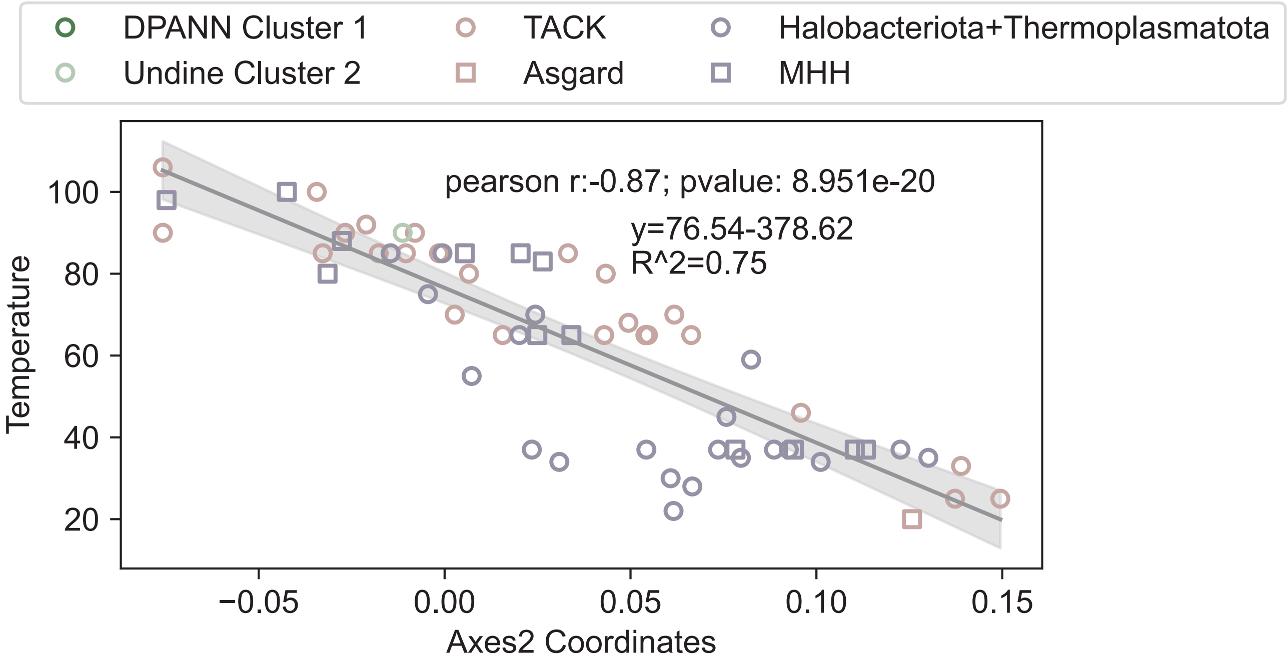


**Fig. S18: Correlation between optimal growth temperature and the second axis of a correspondence analysis on the amino acid composition of our concatenation.** Each point is an extant archaeal genome; the line shows the fitted regression (shaded 95% CI) between OGT and the second axis of a correspondence analysis on the amino-acid composition of the concatenated alignment. This calibrated relationship was then used to predict growth temperatures for ancestral nodes across the archaeal tree. Scale bar: average number of substitutions per site.

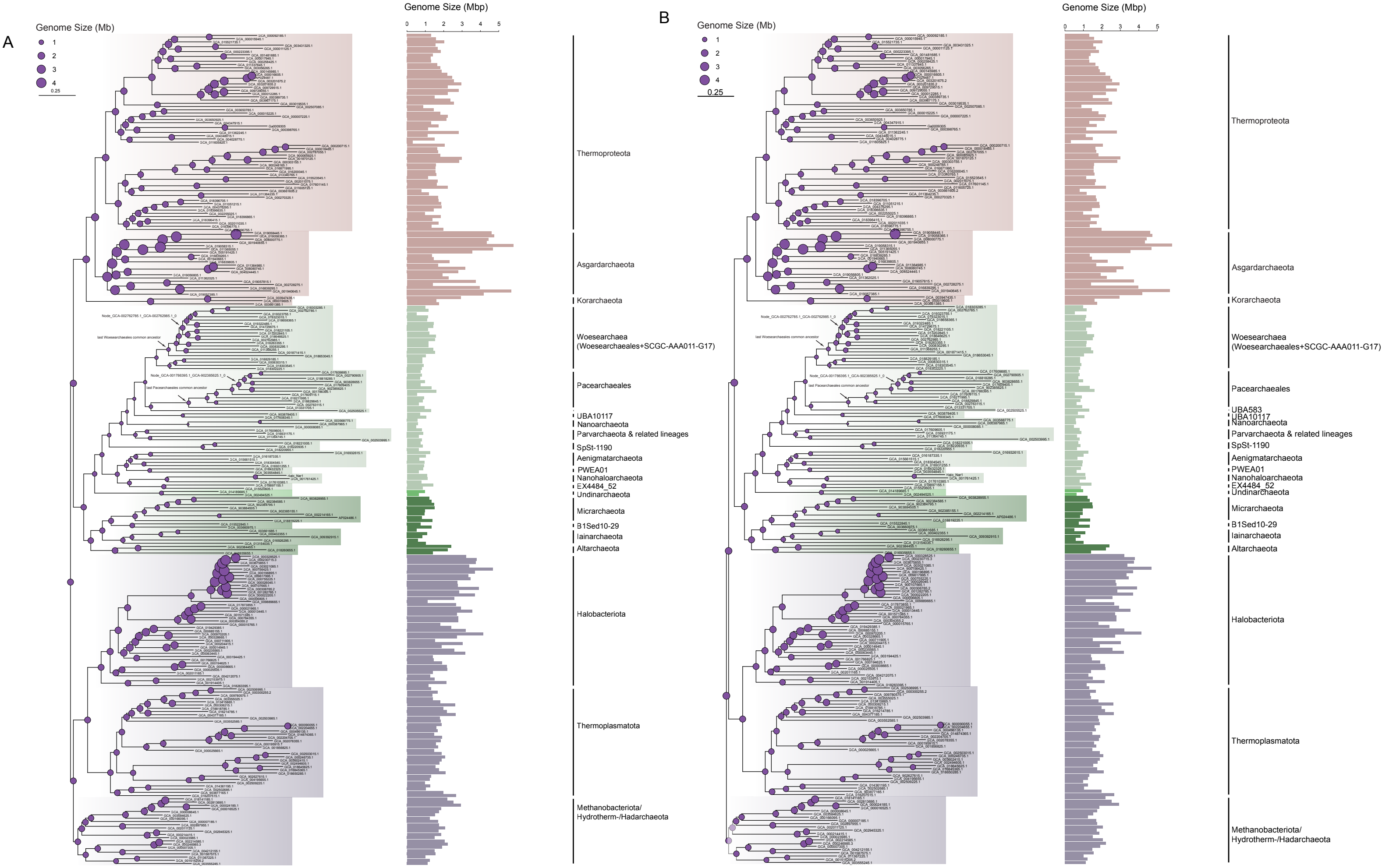


**Fig. S19: Evolution of the arCOG family and inferred genome size over the archaeal tree under the Euryarchaeota root (A) and the MHH root (B).** Genome sizes were predicted from the relationship between arCOG family members and genome size among extant Archaea (LOESS regression) in Fig. S9. The diameter of the circle is proportional to the inferred genome size. Barplot height scaled with the genome size of modern archaea. The largest increase in genome size of Woesearchaeales and Pacearchaeales is labelled.
